# Models to Estimate Genetic Gain of Soybean Seed Yield from Annual Multi-Environment Field Trials

**DOI:** 10.1101/2023.05.13.540664

**Authors:** Matheus D. Krause, Hans-Peter Piepho, Kaio O. G. Dias, Asheesh K. Singh, William D. Beavis

## Abstract

Genetic improvements of discrete characteristics such as flower color, the genetic improvements are obvious and easy to demonstrate; however, for characteristics that are measured on continuous scales, the genetic contributions are incremental and less obvious. Reliable and accurate methods are required to disentangle the confounding genetic and non-genetic components of quantitative traits. Stochastic simulations of soybean (Glycine max (L.) Merr.) breeding programs were performed to evaluate models to estimate the realized genetic gain (RGG) from 30 years of multi-environment trials (MET). True breeding values were simulated under an infinitesimal model to represent the genetic contributions to soybean seed yield under various MET conditions. Estimators were evaluated using objective criteria of bias and linearity. Results indicated all estimation models were biased. Covariance modeling as well as direct versus indirect estimation resulted in substantial differences in RGG estimation. Although there were no unbiased models, the three best-performing models resulted in an average bias of ±7.41 kg/ha^−1^/yr^−1^ (±0.11 bu/ac^−1^/yr^−1^). Rather than relying on a single model to estimate RGG, we recommend the application of multiple models and consider the range of the estimated values. Further, based on our simulations parameters, we do not think it is appropriate to use any single models to compare breeding programs or quantify the efficiency of proposed new breeding strategies. Lastly, for public soybean programs breeding for maturity groups II and III in North America from 1989 to 2019, the range of estimated RGG values was from 18.16 to 39.68 kg/ha^−1^/yr^−1^ (0.27 to 0.59 bu/ac^−1^/yr^−1^).

## 3 Introduction

The purpose of plant breeding is to improve genetic contributions to plant characteristics. As such, plant breeders want to demonstrate the return on investment for the funding to run the breeding program. For many discrete characteristics such as fruit and seed color, pubescence, herbicide and disease resistance, etc., the genetic improvements are obvious and easy to demonstrate. However, for most continuous characteristics, such as seed or grain yield, the genetic contributions are incremental and difficult to separate from management practices and variable, changing environments. Plant breeders have used various experimental and statistical methods to estimate incremental genetic improvements of quantitatively inherited traits that show continuous characteristics (e.g., Byrum *et al*. 2017; Rincker *et al*. 2014). However, with the exception of Rutkoski (2019b), the various proposed methods have not been compared using objective criteria. Herein we utilize a simulation approach to propose and investigate linear models to obtain estimates of genetic improvements to quantitative traits.

For over a century, we have recognized that traits evaluated on continuous scales (P) are composed of discrete polygenic effects (G), continuous environmental effects (E), and GE interaction effects (GEI) (Lynch and Walsh 1998; Sprague and Federer 1951; Mayr 1942; Fisher 1918). Consequently, the simple linear model P = G + E + GE has been successfully used to investigate responses to selection (Tabery 2008; Johannsen 1911). The genetic component of P is modeled as the sum of additive effects of alleles at multiple loci and non-additive genotypic (dominance and epistatic) effects. For diploid species, additive effects refer to effects associated with discrete alleles that can be inherited across generations, while non-additive genetic effects refer to genotypes that are not transmitted through inheritance (Singh *et al*. 2021). Thus, cycles of selection and reproduction affect genetic variability through inherited alleles, also known as breeding values (Falconer 1960; Kempthorne 1957). In other words, realized genetic gain (RGG) is accomplished by the accumulation of alleles with additive effects through recurrent cycles of selection (Bernardo 2020; Walsh and Lynch 2018; Lynch and Walsh 1998).

Applications of this theory were originally demonstrated using population improvement methods (Jenkins 1940). An original sample drawn from an unselected population is known as the founder population and is referred to as cycle zero (C0). Based on evaluated phenotypic values, a subset from C0 is selected to be randomly inter-crossed. The progeny generated by inter-crossing the members of the selected group represents a Mendelian sample of all possible progeny, and is referred to as the C1 population. The process used to create C1 is reiterated with a sample from the C1 population to create C2, which is used to create C3, etc. The process is continuous, and superior individuals are generated in each cycle due to transgressive segregation (Rieseberg *et al*. 1999).

Within a cycle of population improvement, the difference between the average phenotypic value of the population and of a selected sub-group is known as the selection differential (*S*). The difference between the average phenotypic values representing two consecutive cycles of recurrent selection is known as the realized response to selection (*R*). Thus, under Fisher’s additive (linear) model (Fisher 1918), *R* equals the mean breeding value of members that are selected for inter-crossing (Walsh and Lynch 2018), and the relationship between *S* and *R* is given by the breeder’s equation: *R* = *h*^2^*S*, where *h*^2^ is the narrow-sense heritability (Lush 1937). Since only breeding values, i.e., additive allelic effects, are transmitted from one cycle to the next for diploid species, the ratio *R/S* assumes values between zero and one and is known as the realized heritability 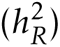 (Bernardo 2020; Hallauer and Miranda 1988). The 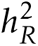 associated with *R* and *S* can be used to obtain an estimate of RGG. Alternatively, it has been suggested that RGG can be estimated from the slope of the regression line of average breeding values on breeding cycles across time (Fritsche-Neto *et al*. 2023; Rutkoski 2019a; Eberhart 1964).

While RGG has been used to compare various population improvement methods in maize (Dudley and Lambert 2004; Hallauer and Miranda 1988; Jenkins 1940), since about 1980, these methods have not been used routinely to develop competitive hybrids. Rather, various types of cultivar development methods have been developed for maize and most commodity crops (Singh *et al*. 2021). Nonetheless, Duvick and co-workers demonstrated that previously sold maize hybrids could be used to assess progress in commercial hybrid maize breeding programs (e.g., Duvick 2005, 1984, 1977). Their approach consisted of sampling a few maize hybrids to represent time periods (eras) in which the hybrids were grown by farmers, and evaluating these in replicated field trials conducted in a common set of environments (location-year combinations). The estimated genotypic values for hybrids were regressed across the years when the hybrids were released to obtain an estimated trend (slope). As with maize, RGG in soybean has been estimated using “era trials” (Milioli *et al*. 2022; Bruce *et al*. 2019; Felipe *et al*. 2016; Rogers *et al*. 2015; Rincker *et al*. 2014; Fox *et al*. 2013; Ustun *et al*. 2001; Wilcox 2001).

Because era trials use designed experiments in which released cultivars represent a “treatment” replicated across the same set of environments, treatments are not confounded with environments. However, inferences from era trials are limited because there are only a select few cultivars representing each era and they are evaluated at only a few locations in a few years. Thus, changes in the genotypic component across time represent an estimated commercial gain, which is not directly interpreted as an estimate of RGG (see Discussion). Further, the years in which the trials are conducted will favor newer cultivars that are more likely to be adapted to the environments in which the era trial is conducted (Rizzo *et al*. 2022). For example, older cultivars were replaced with cultivars that were resistant to emerging pests (Piepho *et al*. 2014). Thus, there can be an inherent bias in favor of recently released cultivars.

Alternatively, historical field data from routine annual multi-environment trials (MET) have been used to assess changes across time for crops including common bean (de Faria *et al*. 2018), potato (Ortiz *et al*. 2022), rice (Streck *et al*. 2018; Breseghello *et al*. 2011), rye (Laidig *et al*. 2017), sugarcane (Ellis *et al*. 2004), sunflower (de la Vega *et al*. 2007), wheat (Gerard *et al*. 2020; Crespo-Herrera *et al*. 2018), among other commercial crops and forage grasses (Piepho *et al*. 2014; Laidig *et al*. 2014). The advantages of using historical MET data are that (*i*) the datasets are usually already available and (*ii*) the data represent a larger number of sampled environments. However, historical MET usually have low connectivity because most experimental genotypes are culled annually. Thus, the disadvantages are that (*i*) environmental effects are mostly estimated from the small sample of connected check cultivars, and (*ii*) new breeding programs usually do not have large historical datasets (Covarrubias-Pazaran 2020).

The most widely used method for partitioning genetic and non-genetic effects is the linear mixed model (LMM, Henderson 1949, 1950; Henderson *et al*. 1959). For a LMM, Best Linear Unbiased Estimators (BLUE) are used to obtain estimates of fixed effects, and Best Linear Unbiased Predictors (BLUP) are used to obtain predicted values for random effects. In state-of-the-art software, both estimators utilize Residual Maximum Like-lihood (REML, Patterson and Thompson 1971) algorithms to obtain empirical values, i.e., eBLUE and eBLUP values. In general, genetic trends are computed from the regression of the eBLUE or eBLUP values of genotypes on the first year of testing or the year of germplasm release. Alternatively to the application of LMM, algorithmic modeling (Byrum *et al*. 2017) or a combination of algorithmic and linear modeling have been proposed to remove environmental effects in order to estimate RGG (Bornhofen *et al*. 2018; Oury *et al*. 2012; Brisson *et al*. 2010).

A simulation approach designed to evaluate the accuracy and precision of estimators of RGG using historical data from MET in plant breeding was conducted by Rutkoski (2019b). In this work, simulation parameters were obtained from medium to low-budget cultivar development programs conducted by the International Rice Research Institute. In total, the author simulated 80 *indica*-type rice breeding programs assuming two levels of heritability (low and high), the inclusion of either a positive or negative non-genetic trend linked to the calendar year (i.e., years of breeding operation), and two breeding schemes. The RGG was then estimated for a quantitative trait (1,000 loci) from era and yield trials with different modeling strategies based on LMM. The study concluded that (*i*) the evaluated estimators were inaccurate, (*ii*) the error associated with the estimates was dependent on the breeding scheme, non-genetic trend, and heritability, and that (*iii*) if the goal is to only determine if there is RGG, some indicators like the expected rate of genetic gain and the equivalent complete generations are useful (Boichard *et al*. 1997). Herein, we use a similar simulation approach as Rutkoski (2019b) to evaluate estimators of RGG for public soybean breeding programs. It is evident the genotypic sampling space in a breeding program is complex due to multiple parental genotypes, families in early trials, check cultivars, experimental genotypes, and introductions of germplasm from external sources. In this context, the first step toward estimating RGG in a cultivar development programs is to define the inference space - We define the breeding population as consisting of evaluated homozygous genotypes (i.e., purlines) that will be used as parents in crossing blocks. Estimates of genetic trends across time based on MET data are associated with selected genotypes that were or will be hybridized to create a new population of genotypes for evaluation. As such, the breeding population represents the tail of a distribution with twice the additive genetic variance expressed in the F2 generations of newly created progeny (Bernardo 2020; Lynch and Walsh 1998). Thus, the estimated genetic trends across years of MET is a function of the breeding values; and therefore, is interpreted herein as an estimate of RGG.

Simulations were based on knowledge of the organization of public soybean breeding programs for Maturity Groups II and III in North America, as well as previously analyzed MET data generated by public soybean breeders (Krause *et al*. 2022). These data consist of 4,257 genotypes, 63 locations, and 31 years, resulting in 591 observed environments from 1989 to 2019. The true RGG from simulation was computed according to our novel definition of RGG for complex breeding programs. We also implemented in the simulator an alternative way to simulate non-genetic gain based on empirical frequencies of locations used in the historical MET. Thus, the simulator was built to (*i*) evaluate estimators of RGG based on realistic data routinely collected by public soybean breeders, (*ii*) investigate the bias of several models used to estimate RGG according to the true simulated RGG, and (*iii*) determine if there is any RGG, regardless of estimated bias. Lastly, based on the best-performing models, we reported estimates of RGG for the soybean empirical dataset from Krause *et al*. (2022).

## 4 Material and Methods

### 4.1 Soybean stochastic simulations

A soybean breeding program with 46 years of operation was simulated using AlphaSimR (Gaynor *et al*. 2021) and functions developed in the R programming environment (R Core Team 2021). The simulation code was run in parallel (3 cores per replicate) in a bash shell under Linux. Each simulated run took, on average, four hours to be completed in computer nodes with 30 GB of RAM and Intel processors with speeds ranging from 3.20 to 3.50 GHz. Statistical analyses were performed using Asreml-R version 4.1 (Butler *et al*. 2017) and R base functions. The simulation parameters (e.g., initial trait mean, variance components, number of crosses and genotypes, etc.) were obtained from the analysis of historical MET soybean data from Krause *et al*. (2022).

#### 4.1.1 Founder population

A set of 499 pureline soybean genotypes from maturity groups II and III previously developed by public soybean breeders in the North Central Region of the United States were used as the founder population. The pureline genotypes were genotyped using the SoySNP6K BeadChip (Song *et al*. 2020). Single nucleotide polymorphism markers (SNP) were removed if missing scores were *>* 20%, or if the minor allele frequency was *<* 0.05. Genotypes were removed if heterozygosity *>* 0.0625 and/or *>* 20% of the markers were missing data. The final number of SNP markers was 5,279, ranging from 204 to 349 markers per chromosome. A principal components analysis of the additive genomic relationship matrix G_M_ (Endelman and Jannink 2012) do not show evidence of any grouping pattern among this set of founders (Figure A1).

#### 4.1.2 Breeding values

Simulated breeding values were obtained assuming Fisher’s infinitesimal model to approximate polygenic effects on soybean seed yield. Eligible additive quantitative trait loci (QTL) were randomly assigned to 1,000 loci distributed across 20 chromosomes, according to the genetic linkage map (2145.5 cM) estimated from the Soybean Nested Association Mapping population (Ramasubramanian and Beavis 2020). Further details are given in Appendix B.

#### 4.1.3 Crossing nurseries and generation of experimental lines

Every year 50-80 biparental crosses between 20-30 selected breeding lines were made to produce a population of F_1_ individuals. Each F_1_ individual was self-pollinated to create fifty to eighty families of F_2_ seeds, each representing the bi-parental cross (to provide information for cross-pollination crop species we are including the ‘S’ symbol for selfing generation; S_0_ generation). From each F_2_ individual plant, 2-3 F_3_ seeds were created to represent the S_1_ generation. Each F_3_ individual was self-pollinated to create F_4_ seeds representing the S_2_ generation. All of the self-pollinated seed from each F_4_ individual was combined to represent an F_4:5_ experimental line. Selections were not performed during inbreeding generations.

#### 4.1.4 Selection of experimental lines

Phenotypic values of F_4:5_ experimental lines were simulated for evaluation in an unreplicated field trial at a single location, designated as breeders trial 1 (BT1). Self pollinated seeds (F_4:6_) from selected F_4:5_ experimental lines were subsequently evaluated at two locations (BT2). Self pollinated seeds (F_4:7_) from the selected F_4:6_ experimental lines were then evaluated at three locations (BT3). The best F_4:7_ experimental lines selected from the BT3 were evaluated in regional trials. The first year of regional trials consisted of F_4:8_ experimental lines evaluated at eight locations, and was referred to as the preliminary yield test (PYT). If selected, F_4:8_ experimental lines were evaluated in the uniform regional test (URT) at 12 locations. Experimental lines in PYT and URT (advanced MET) can be thought of as candidate varieties (Figures 1B and 1C).

**Figure 1:**
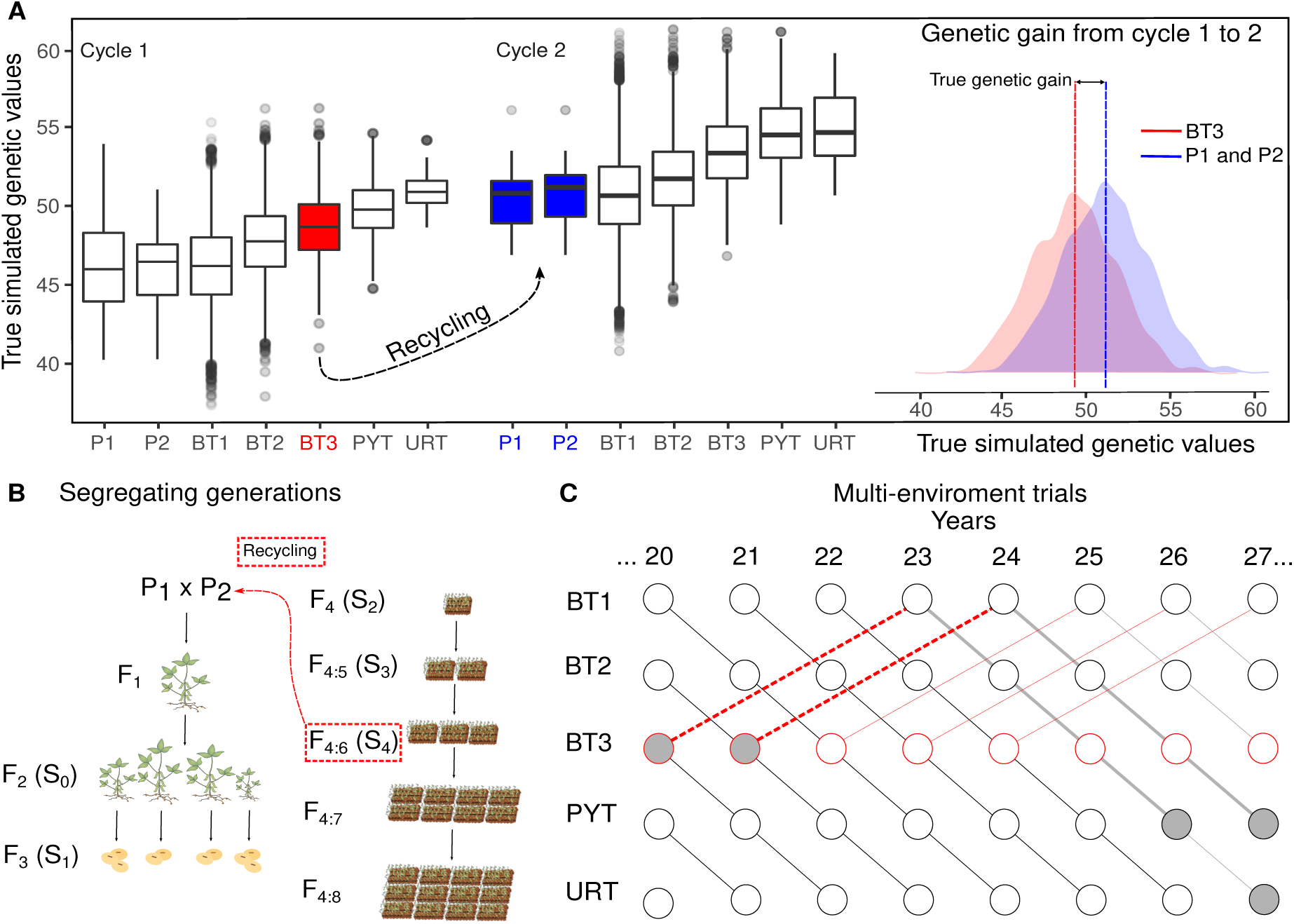
Description of the simulated RGG between cycles 1 and 2 of line development. Experimental lines selected from BT3 of cycle 1 are crossed in a nursery to create a new population that will be used to create experimental lines for evaluation in cycle 2 (A). Experimental lines evaluated in PYT and URT include experimental lines selected from BT3 for crossing (B). For example, breeding lines selected in year 20 and evaluated *per se* in the PYT of year 21, will have progeny developed as experimental lines evaluated in the PYT of year 26 (C).

Ten percent of the BT1 experimental lines were advanced to BT2. For the remaining trials, a 20% selection intensity was considered. Yearly selections were carried out by ranking the predicted eBLUP values of genotypic means obtained only from the data of the current year (Model C2, Appendix C). For example, genotypes that advanced from BT2 to BT3 were ranked according to their BT2 data. Although molecular markers were not used for selection (e.g., genomic selection), some of the LMM used to estimate RGG need information about covariance among relatives. We did not include genomic selection to provide applicability for a wide variety of breeding programs globally, with varying resources. Thus, the experimental lines in BT1 phase of development were genotyped with an SNP chip consisting of 2,400 SNP markers. A full description of the MET simulation models is given in Section 4.1.6.

Field trials included three to six check cultivars, the actual number being randomly determined for each simulation run (Figure 2A and A2). The number of locations in each trial was fixed. However, actual locations were replaced across years to mimic the actual practice of occasionally changing locations in a region from year to year (Figure 2B and A3). The field plot design was assumed to be completely randomized with a single replicate for BT1, two replicates for BT2, and three replicates for the remaining trials. A random percentage from zero to 12% of individual field plots were considered to be missing data within each trial. Data from single trials were analyzed with Model C3 (Appendix C).

**Figure 2:**
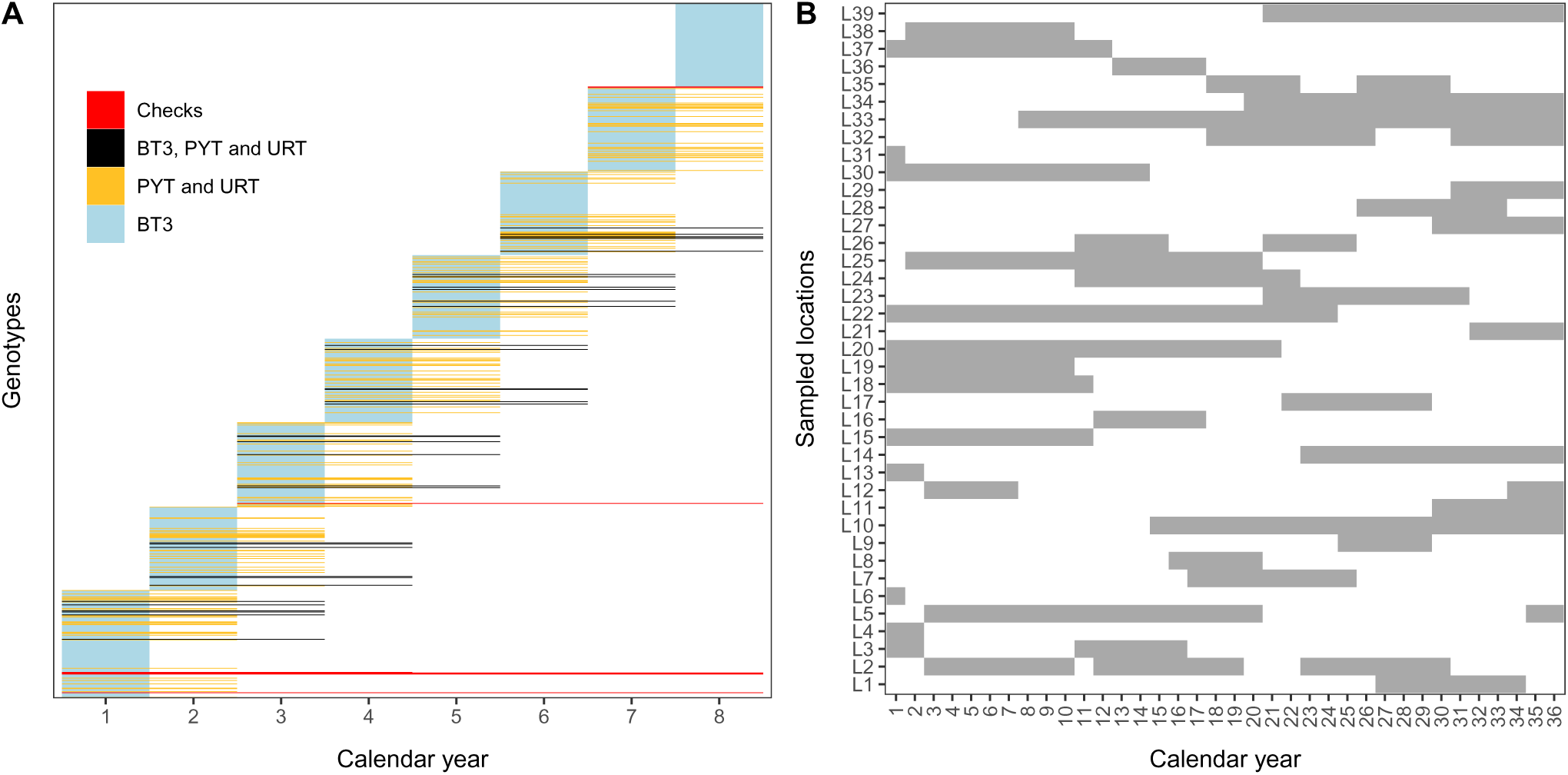
Representation of the simulated (A) selection process in MET (B) and the rate of locations replacement across years.

#### 4.1.5 Selection of breeding lines and cycle of line development

BLUP selection of parental or breeding lines were performed in BT3 with phenotypic data from BT1, BT2, and BT3 (Model C1, Appendix C). The time per cycle of line development was assumed to be five years based on the assumption that the creation of the experimental lines could be conducted at off-site continuous nurseries. Thus, it will take five growing seasons, 2 years, from initial crosses to create F1 seed to creating F_4:5_ experimental lines for evaluation on-site in BT1. And assuming that there is only one growing season for evaluating the crop, each of the BT’s will require 1 year (Figure 1B). Breeding lines were not selected in the first five years of the simulated breeding programs (i.e., not enough data). Rather the simulated crosses were randomly sampled from the founder population. Moreover, these initial years were not used to estimate RGG (Figure A4).

**Figure 4:**
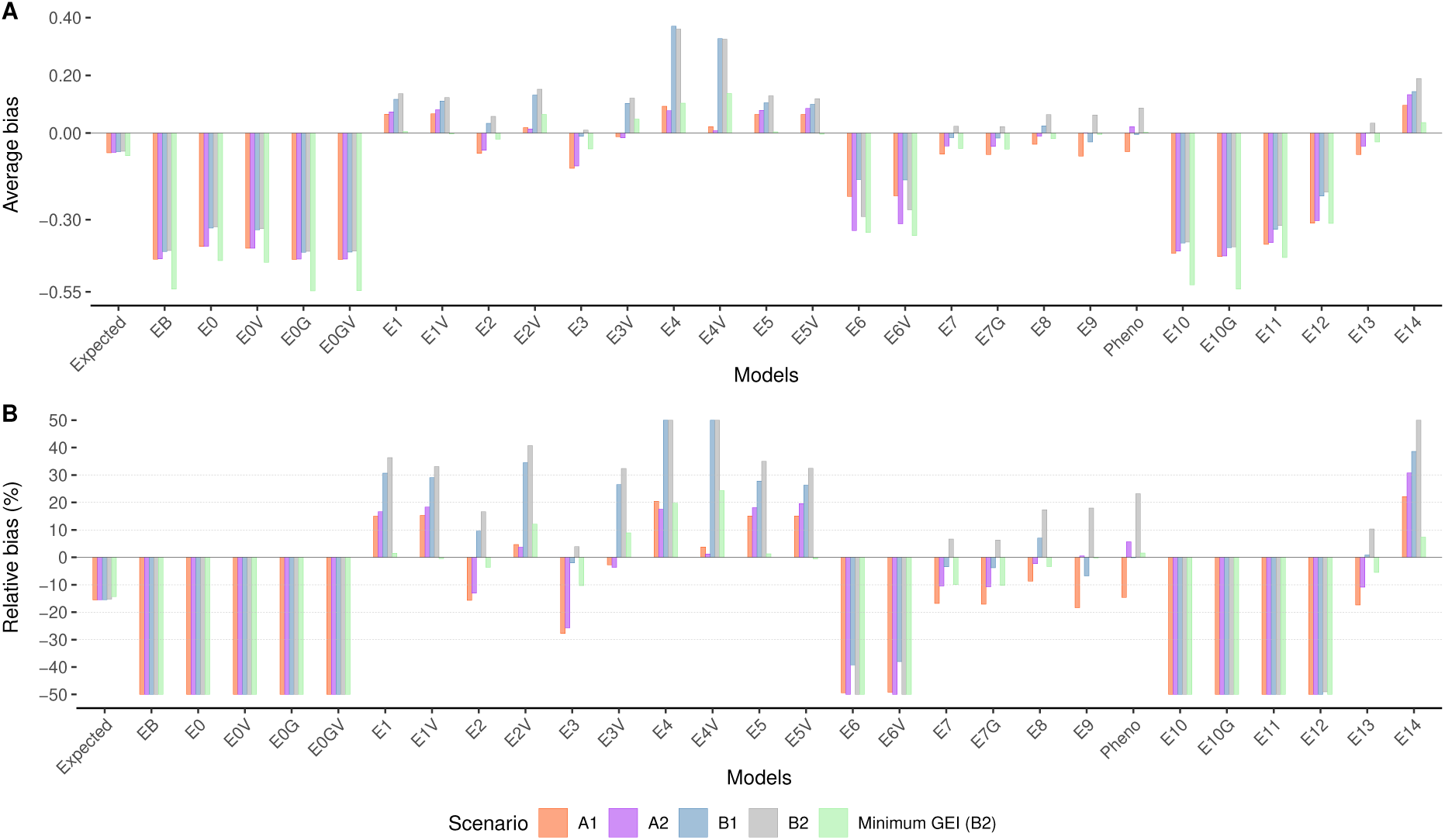
Average bias in bu/ac^−1^/yr^−1^ (A) and relative bias (B) across simulations. The y-axis in B was limited to ±50% to enhance visualization.

#### 4.1.6 Simulations of multi-environment trials

Genotype by environment effects were defined as the sum of genotype × location (*GL*), genotype × year (*GY*), and genotype × location × year (*GLY*) interaction effects, i.e., *GEI* = *GL* + *GY* + *GLY* . Genotype refers to the experimental lines and QTL effects assigned to genotypes at the beginning of simulation runs and retained through all stages of line development. The simulation of the main genotypic effects (*G*) was accomplished using AlphaSimR as described in Section 4.1.2, with the matrix of additive QTL genotypes being carried further to calculate GEI effects. Individual plot phenotypic values were simulated according to the following general linear model:

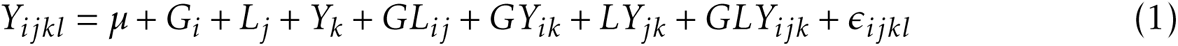

where *Y_ijk_* is the simulated phenotype for the *ijk^th^* genotype (line) × location × year combination in the *l^th^* plot, *µ* is the intercept or mean trait value, and *ɛ_ijkl_* is the residual plot to plot variability associated with the simulated phenotypic value. The remaining terms *L* and *Y* represent the locations and years’ main effects, respectively. All model terms except *L* were considered random effects sampled from various distributions described below (Section 4.1.6.1). Locations were simulated as fixed effects to incorporate estimated values from empirical data (Figure A5). Individual trial effects were not simulated.

##### 4.1.6.1 Specific simulation models

We simulated six conditions for MET and label them A1, A2, B1, B2, B2-M, and B2-R. The “A” conditions represent simple genetic effects models where random effects from Model 1 were simulated as independent with homogeneous variances. The “B” conditions represent complex models where each simulation run (*s* = 1*, . . ., S* = 225) had a unique 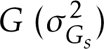 and 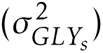 variance component. Simulation runs represent independent, stochastically simulated breeding programs. Also, the simulations of the B conditions consisted of correlated additive QTL effects across locations and years, for the GL and GY interaction effects, respectively. Simulations labeled B2-M are a modification of B2 where the GL, GY, and GLY sampled variances were divided by an *ad-hoc* factor of 10 (i.e., minimum GEI). Lastly, simulations labeled B2-R represent B2 where a random sample of experimental and breeding lines were retained. Heterogeneous residual variances represented plot-to-plot variability within trials and were sampled from a Log-Logistic distribution (Tables 1 and 2).

**Table 1:**
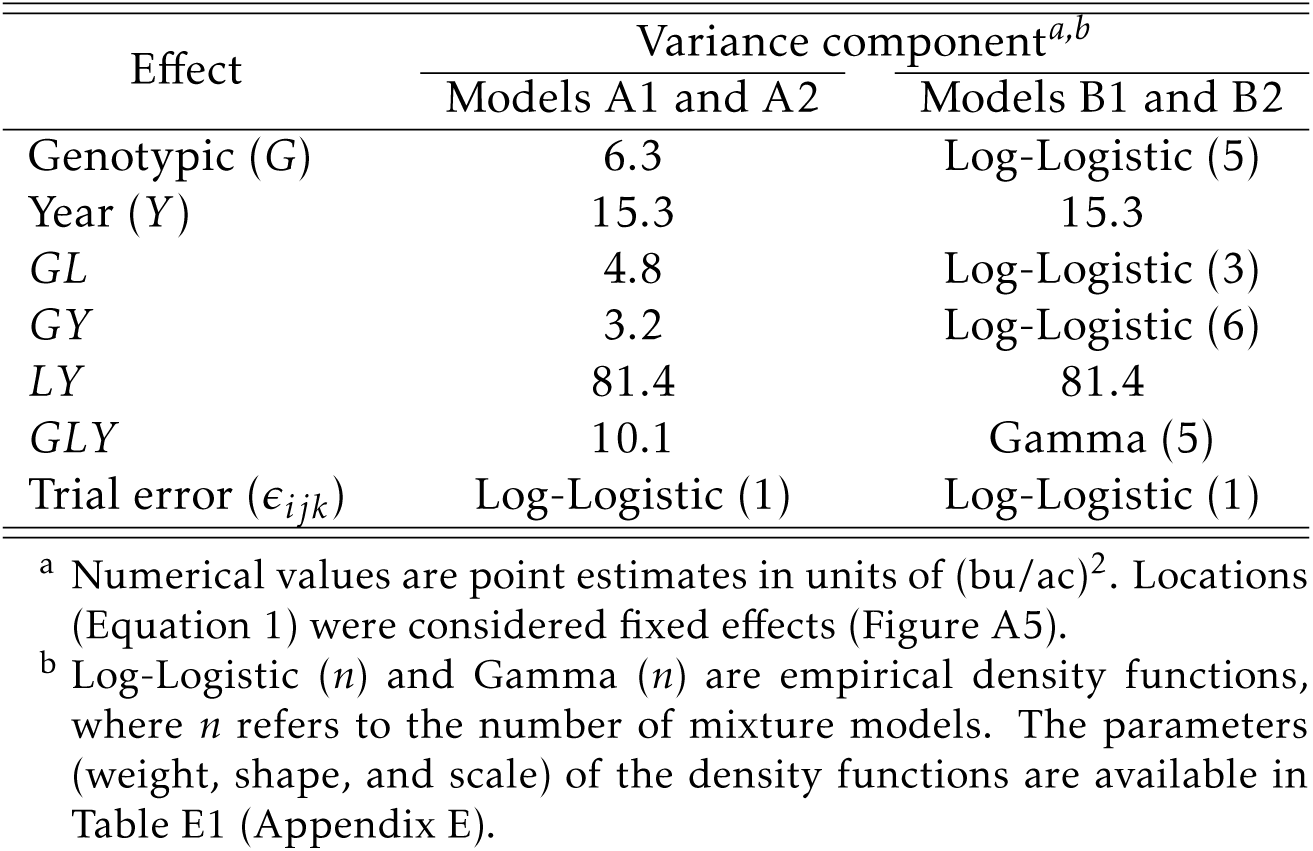
Variance component values and probability distributions implemented in the simulator.

The GEI simulation approach for the “B” simulation models aimed to generate data that have a similar structure to the empirical data analyzed by Krause *et al*. (2022). A step-by-step description of the simulation of the corrected QTL effects is given in Appendix D. Briefly, let us define *_I_* Θ_1000_ as the matrix of QTL dosages (0, 1, and 2) simulated by AlphaSimR. The matrix Θ has dimension *I* ×1, 000, where *I* (*i* = 1*, . . ., I*) represents the number of genotypes and *q* = 1, 000 is the number of simulated QTL. The sampled effects are defined in matrix notation as 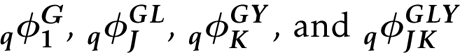 for model terms *G, GL, GY*, and *GLY*, respectively. Their dimensions are represented by *I, J, K* for geno-types, locations (*j* = 1*, . . ., J*), and years (*k* = 1*, . . ., K*), respectively. Note 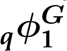 is a vector of length 1, 000 as defined in Section 4.1.2. This notation is general and reflects the sample size associated with the model parameters in each trial (Tables 1 and 2).

**Table 2:** Fixed effect and distributional assumptions for random effects implemented in the simulator. G, L, and Y represent genotypic (line), location, and year factors. Note the sampling for the genotypic-related terms G, GL, GY, and GLY were at a QTL level.

| Model terms | Simple effects<sup>a</sup> | Complex effects<sup>a</sup> |
| --- | --- | --- |
|  | Simulation models A1 and A2<sup>b</sup> | Simulation models B1 and B2<sup>b</sup> |
| Intercept ($\mu$) | 46 | 46 |
| Genotypic (G) | $I\Theta_q \cdot q\phi_1^G$, where $q\phi_1^G \sim N(\mathbf{0}_q, \sigma_G^2 \mathbf{I}_q)$ | $I\Theta_q \cdot q\phi_1^G$, where $q\phi_1^G \sim N(\mathbf{0}_q, \sigma_{G_s}^2 \mathbf{I}_q)$ |
| Location (L) | Fixed | Fixed |
| Year (Y) | $N(\mathbf{0}_K, \sigma_Y^2 \mathbf{I}_K)$ | $N(\mathbf{0}_K, \sigma_Y^2 \mathbf{I}_K)$ |
| GL | $I\Theta_q \cdot q\phi_J^{GL}$, where $q\phi_J^{GL} \sim N(\mathbf{0}_J, \sigma_{GL}^2 \mathbf{I}_J)$ | $I\Theta_q \cdot q\phi_J^{GL}$, where $q\phi_J^{GL} \sim N(\mathbf{0}_J, \Sigma_{J_s})$ |
| GY | $I\Theta_q \cdot q\phi_K^{GY}$, where $q\phi_K^{GY} \sim N(\mathbf{0}_K, \sigma_{GY}^2 \mathbf{I}_K)$ | $I\Theta_q \cdot q\phi_K^{GY}$, where $q\phi_K^{GY} \sim N(\mathbf{0}_K, \Sigma_{K_s})$ |
| LY | $N(\mathbf{0}, \sigma_{LY}^2 \mathbf{I}_{JK}) + (z_{jk} \times \eta_j)$ | $N(\mathbf{0}, \sigma_{LY}^2 \mathbf{I}_{JK}) + (z_{jk} \times \eta_j)$ |
| GLY | $I\Theta_q \cdot q\phi_{JK}^{GLY}$, where $q\phi_{JK}^{GLY} \sim N(\mathbf{0}_{JK}, \sigma_{GLY}^2 \mathbf{I}_{JK})$ | $I\Theta_q \cdot q\phi_{JK}^{GLY}$, where $q\phi_{JK}^{GLY} \sim N(\mathbf{0}_{JK}, \sigma_{GLY_s}^2 \mathbf{I}_{JK})$ |
| Trial error<sup>c</sup> | $N(\mathbf{0}, \oplus_{b=1}^B \sigma_{\epsilon_b}^2 \mathbf{I}_{n_b})$ | $N(\mathbf{0}, \oplus_{b=1}^B \sigma_{\epsilon_b}^2 \mathbf{I}_{n_b})$ |

<sup>a</sup> The matrix $\Theta$ has dimension $I \times 1,000$ , where $I(i = 1, \dots, I)$ represents the number of genotypes and $q = 1,000$ is the number of simulated QTL. The sampled effects are defined in matrix notation as $q\phi_1^G$ , $q\phi_J^{GL}$ , $q\phi_K^{GY}$ , and $q\phi_{JK}^{GLY}$ for model terms G, GL, GY, and GLY, respectively. Their dimensions are represented by $I, J, K$ for genotypes, locations ( $j = 1, \dots, J$ ), and years ( $k = 1, \dots, K$ ), respectively. Note $q\phi_1^G$ is a vector of length 1,000 and it was simulated as defined in Section 4.1.2. $N$ represent the Multivariate Normal distribution and · matrix multiplication. The matrices $\Sigma_{J_s}$ and $\Sigma_{K_s}$ are positive-definite covariance matrices for GL and GY effects, respectively. $\mathbf{0}$ is a vector of zeros and $\mathbf{I}$ an identity matrix with their respective dimensions.
<sup>b</sup> The interaction LY simulates the inclusion of a positive non-genetic trend in models A2 and B2, according to the mapping variable $z_{jk}$ described in Equation 2. In scenarios A1 and B1, the non-genetic trend was not simulated, hence $z_{jk}$ was absent. For details, see Section 4.1.6.2.
<sup>c</sup> Trials ( $b = 1, \dots, B$ , with a sample size of $n_b$ ) have heterogeneous residual variances ( $\sigma_{\epsilon_b}^2$ ), where $\oplus$ is the direct sum operator.

##### 4.1.6.2 Simulation of the non-genetic trend

Simulation models A2 and B2 mimic the inclusion of the positive non-genetic trend, whereas models A1 and B1 do not (Tables 1 and 2). To simulate a positive non-genetic trend, we propose that the location × year interaction term (*LY_jk_*) from Model 1 be further split into two components:

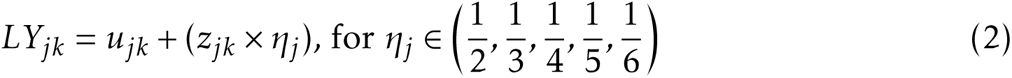

where 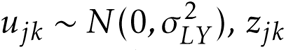 is a location-year covariate mapping if the *j^th^* location was observed in the *k^th^* and (*k* + 1)*^th^* years, and *η_j_* is a constant of the non-genetic gain randomly chosen with equal probability. Equation 2 was designed to simulate a (cumulative) positive non-genetic trend by adding increments of *η_j_* units (bu/ac) every time a specific location was observed across consecutive years. This mimics improvements in management practices in field trials that are continuously used by the breeding program, which is linked to the location × year effect. For example, location “L1” is used for phenotyping in the current year, and the farmer/researcher identifies a source of variation in the field due to poor fertility. The issue will be addressed for the next year/trial, and the yield in that same piece of land will improve. For clarity, an example of the covariate mapping is provided in Tables 3 and 4.

**Table 3:** Hypothetical example of the covariate mapping (*z_jk_*) used to simulate a cumulative (*z_jk_* × *η_j_*) rate (*η_j_*) of non-genetic gain.

| Calendar year | Is the $j^{th}$ location used? | $z_{jk}$ | $\eta_j$ | How many years has the $j^{th}$ location been used? | Cumulative non-genetic gain ($z_{jk} \times \eta_j$) |
| --- | --- | --- | --- | --- | --- |
| 1 | Yes | 0 | 0 | 1 | 0 |
| 2 | No | - | - | 1 | - |
| 3 | Yes | 1 | 1/2 | 2 | 1/2 |
| 4 | Yes | 2 | 1/2 | 3 | 1/2+1/2 |
| 5 | No | - | - | 3 | - |
| 6 | No | - | - | 3 | - |
| 7 | Yes | 3 | 1/2 | 4 | 1/2+1/2+1/2 |

**Table 4:** Hypothetical example of *L_j_* + *Y_k_* + *LY_jk_* from Model 1 when non-genetic trend is considered in the simulation of field trials.

|  |  |  |  |  |  |  |  |
| --- | --- | --- | --- | --- | --- | --- | --- |
| $L_j + Y_k + LY_{jk} \Rightarrow$ | Fixed effect | + | $Y_k \sim N(0, \sigma_Y^2)$ | + | $LY_{jk} \sim N(0, \sigma_{LY}^2)$ | + | $(z_{jk} \times 0.5)$ |
| ⇒ | +3 | | -2.27 | | -17.42 | + | $(0 \times 0.5)$ [Year 7] |
| ⇒ | +3 | | +2.79 | | -10.40 | + | $(1 \times 0.5)$ [Year 8] |
| ⇒ | +3 | | +3.22 | | +4.23 | + | $(2 \times 0.5)$ [Year 9] |
| | ⋮ | | ⋮ | | ⋮ | | ⋮ |
| ⇒ | +3 | | +9.03 | | +5.53 | + | $(15 \times 0.5)$ [Year 22] |

Annual effects (*Y_k_*) were assumed to be random: there are favorable (positive increments in yield) and unfavorable (negative increments in yield) years. This was simulated by randomly sampling year effects from a Normal distribution (Table 1). If the cumulative non-genetic trend is not simulated, the term *z_jk_* does not appear, and therefore Equation 2 is reduced to *LY_jk_* = *u_jk_*. In this case, the location-year effects are only a function of the random sample from the Normal distribution (*u_jk_*).

### 4.2 Estimation of RGG

The simulated RGG was calculated from the genetic values of breeding (parental) lines. Estimates of RGG from MET were determined with two approaches: (*i*) using data from PYT and URT, and assuming information of breeding lines is not available, and (*ii*) using data from BT3, PYT, and URT, and assuming information from breeding lines is available. The first approach reflects the majority of datasets available. The main difference between these two approaches is that RGG estimated from (*i*) is computed from all experimental lines in advanced MET, and from (*ii*) only from experimental lines used as breeding lines (i.e., pedigree information is available). The estimated RGG values were compared with the simulated RGG to assess the analytic models’ ability to estimate RGG. Estimation models are presented in Tables 6 and 7. Check cultivars were included in the models but not considered to estimate RGG; rather they provide information for the non-genetic trends across years.

**Table 5:**
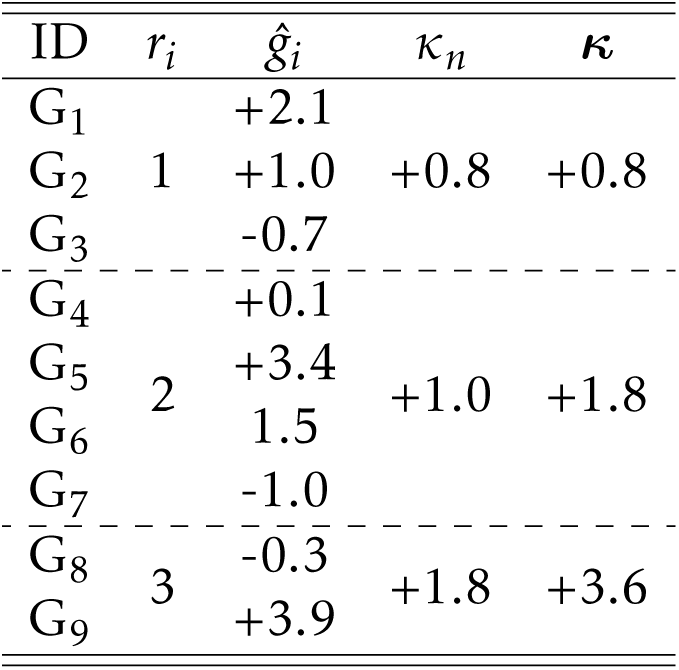
Hypothetical example of the cumulative sum (***κ***) of the average (*κ_n_*) eBLUP values (ĝ*_i_*) linked to the first year of trial (*r_i_*) for nine experimental lines (ID). Note ***κ***_(*s*)_ is a generalization of the expected gain from selection (Rutkoski 2019b; Piepho and Mohring 2007). In addition, it is worthwhile emphasizing ***κ***_(*s*)_ is computed with a full, G×L×Y model. Hence, the variations from year to year (i.e., GY, GLY, LY, and Y) are accounted for.

#### 4.2.1 Simulated RGG from breeding lines

The simulated RGG was calculated as the slope (*β_T_*_(*s*)_) of the regression line of true genetic values of breeding lines (*g_Ti_*_(*s*)_) on the year they were used in crossing blocks (*w_i_*_(*s*)_):

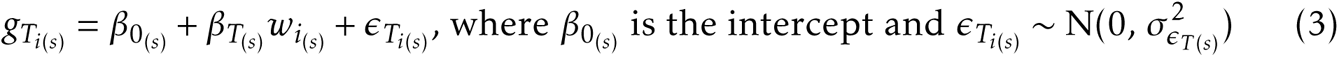

The slope *β_T_*_(*s*)_ represents the rate of accumulation of beneficial (additive) alleles among breeding lines across years of breeding operation. Also, although a breeding line could be crossed multiple times (hub network), it was used only in a single crossing nursery.

#### 4.2.2 Estimates of RGG using PYT and URT

Direct versus indirect estimation of RGG were tested as they differs on how the main genotypic effect (*G_i_*) is decomposed. The direct estimate was computed by replacing *G_i_*with *β_g_*_(*s*)_ *r_i_*_(*s*)_, where *β_g_*_(*s*)_ is a heterogeneous regression coefficient of RGG for experimental lines and check cultivars, and *r_i_*_(*s*)_ is the year of first testing. The year of first testing is a designated year the *i^th^* experimental line is first available in the dataset (e.g., PYT in this case). In addition, note *β_g_*_(*s*)_ is considered heterogeneous to isolate checks’ contributions to RGG. The indirect estimation was computed in two steps. The first step consisted of estimating/predicting genotypic main effects (ĝ*_i_*_(*s*)_), which can be eBLUE or the eBLUP values. The second step estimates RGG as the slope (*β_R_*_(*s*)_) of the linear regression between ĝ*_i_*_(*s*)_ and *r_i_*_(*s*)_, as follows:

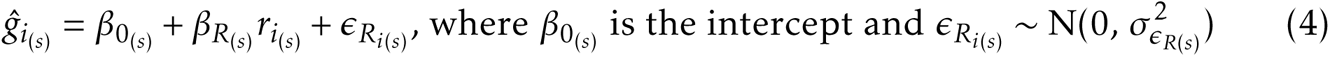

Alternatively, when ĝ*_i_*_(*s*)_ are eBLUP values, RGG was indirectly estimated with the cumulative sum of the average ĝ*_i_*_(*s*)_ values of all lines (***κ***_(_*_s_*_)_) according to the year of first testing (*r_i_*_(*s*)_):

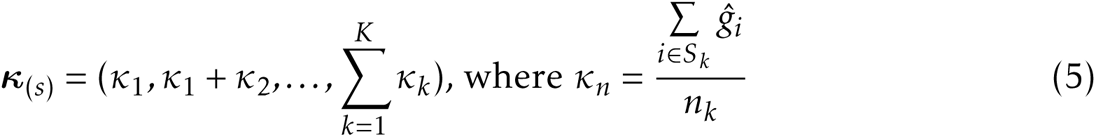

where *K* is the number of years in the dataset, ĝ*_i_* is the predicted eBLUP value of the *i^th^*experimental line, *S_k_* is the set of lines first tested in the *k^th^* year, and *n_k_* is the number of experimental lines evaluated in *S_k_*. The RGG was then estimated by regressing ***κ***_(*s*)_ on *r_i_*_(*s*)_ . An example is presented in Table 5.

#### 4.2.2.1 Estimation models with advanced MET

We applied 21 analytic models to estimate RGG from advanced MET. The first G×L×Y Model EB (benchmark) indirectly estimated RGG with eBLUP values. Model E0 uses the same framework of EB, but RGG was estimated with ***κ***_(*s*)_. Model E1 (Mackay *et al*. 2011) estimated RGG with eBLUE values. Model E2 (Piepho *et al*. 2014) directly estimated RGG with a fixed regression coefficient, and Model E3 with a random regression coefficient. Models E4 and E5 are variations of Models E0 and E1, respectively, where only experimental lines from URT were considered. Model E6 (control population, Rutkoski 2019b) directly estimated RGG (*β_t_*) by contrasting checks and experimental lines with a fixed regression term (*β_t_t_k_*), where *t_k_* is a continuous covariate for the calendar year. Model E7 was designed to assess if environmental (the combination of location-year) effects will be properly modeled using check cultivars. An initial model is applied to the data only with checks to obtain predicted values (i.e., eBLUP) of environmental effects. These predicted values are subsequently used as a fixed continuous covariate in a second analysis model. In the second model, the eBLUP values of experimental lines are predicted and used to estimate RGG indirectly. Model E8 is a variation of E7, in which RGG was directly estimated in the second step (Table 6).

Different covariance structures were utilized to model GL and GY interactions. Models E0, E1, E2, E3, E4, E5, and E6, assumed independent effects with homogeneous variance, whereas their counterparts E0V, E1V, E2V, E3V, E4V, E5V, and E6V, involved correlated random effects with heterogeneous variances. It must be emphasized we only consider diagonal and factor-analytic models of first order to avoid convergence issues across simulation runs. For Models E0G, E0GV, and E7G, we also investigated the RGG estimation with genomic estimated breeding values (Table 6).

Lastly, Model E9 takes advantage of both experimental lines and checks replicated between consecutive pairs of years: (1) an initial model was fit within years to obtain eBLUP values for experimental lines (*g_ik_*); and (2) the *g_ik_* values are then corrected for the “year effect” according to the “reference year (R)”. For example, in a MET dataset of 10 years, if R = year 5, the corrections for the year effects will be performed like 1 ← 2 ← 3 ← 4 ← 5 → 6 → 7 → 8 → 9 → 10, with a forward-backward regression algorithm along years. If R = 1, only a forward process (1 → · · · → 10) is used, and if R = 10, only a backward process (1 ← · · · ← 10) is used. The methodology is presented with references in Appendix G. In addition to the described models, RGG was also computed from raw phenotypes at a location level (“Pheno”) to access the impact of estimating RGG without explicit models (Table 6).

**Table 6:**
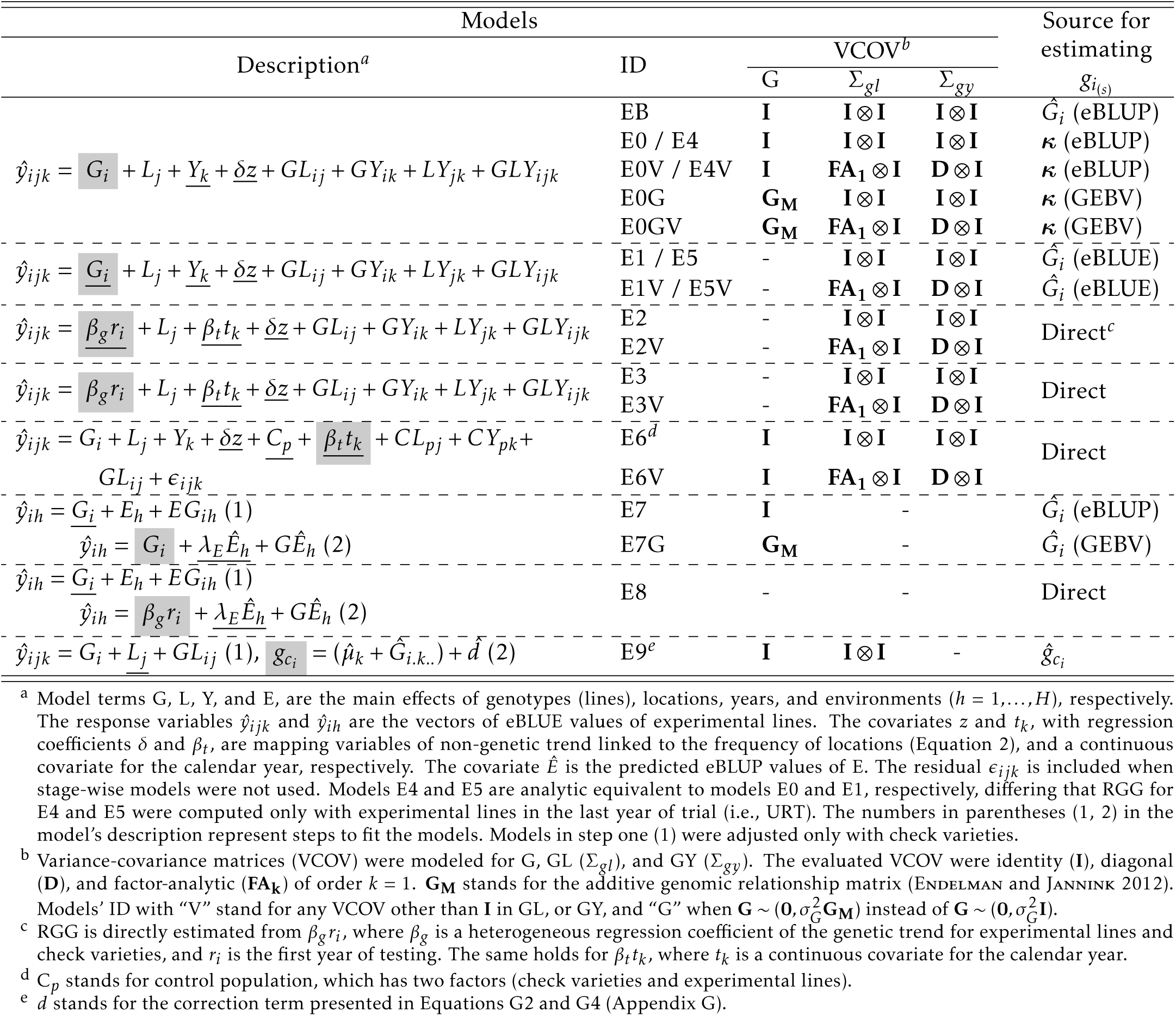
Models for advanced MET. Fixed effects are underlined, and the RGG was estimated with model terms highlighted in gray.

### 4.2.3 Estimates of RGG using BT3, PYT and URT

In addition to exploring data from BT3, here we assume information about lines used in crosses is available. All data were included to fit the evaluated models (Table 7); however, the RGG was computed from breeding lines. The indirect RGG estimate was computed with a modified version of Equations 4 and 5, where ĝ*_i_*_(*s*)_ and ***κ***_(*s*)_ are now restricted to breeding lines, and *r_i_*_(*s*)_ was replaced by *w_i_*_(*s*)_ .

**Table 7:**
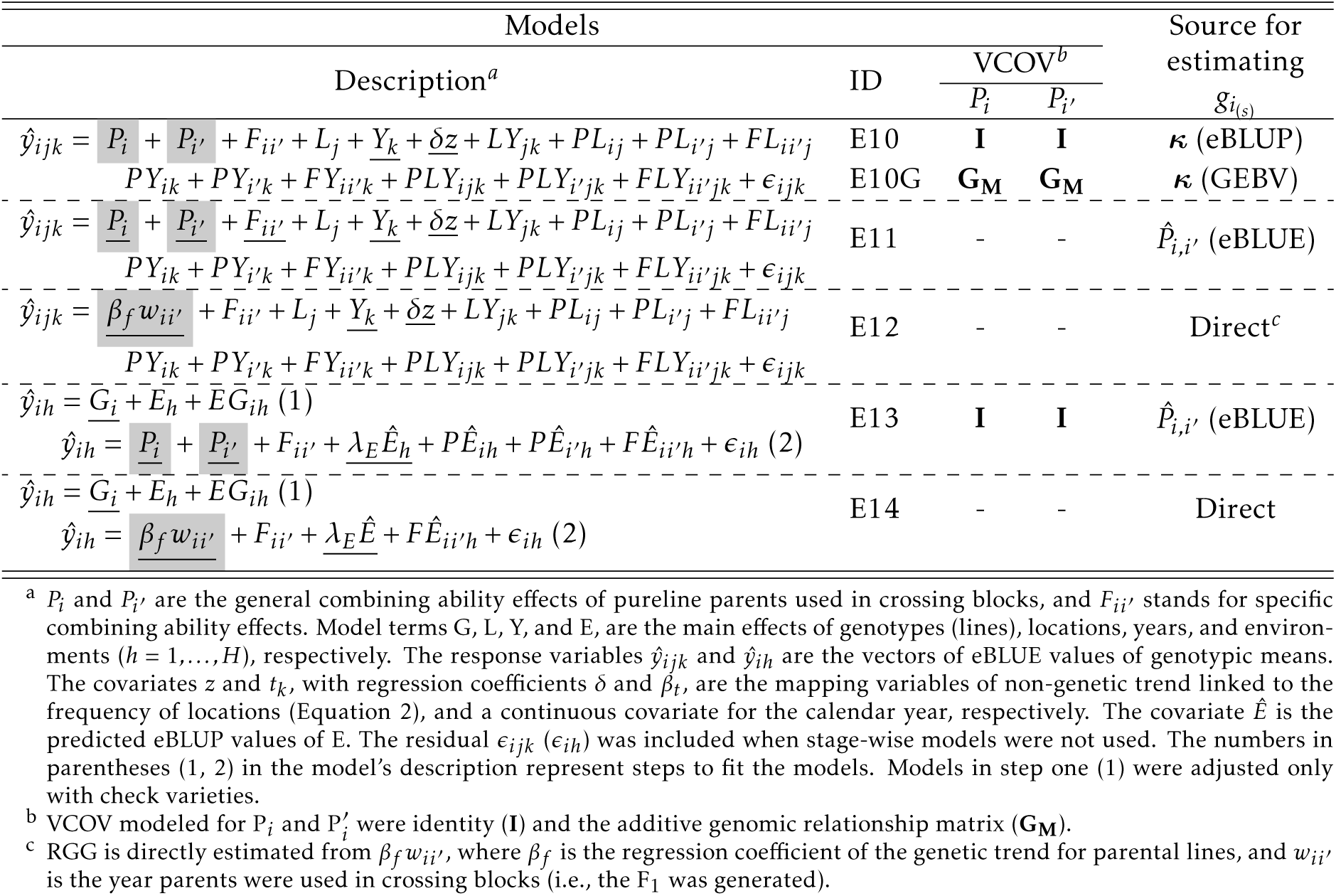
Models with information from breeding lines used in crosses. Fixed effects are underlined, and the RGG was estimated with model terms highlighted in gray.

| Models |  |  |  | Source for estimating |
| --- | --- | --- | --- | --- |
| Description<sup>a</sup> | ID | VCOV<sup>b</sup> | | $\mathcal{G}_{i(s)}$ |
| | | $P_i$ | $P_{i'}$ | |
| $\hat{y}_{ijk} = \underline{P_i} + \underline{P_{i'}} + F_{ii'} + L_j + \underline{Y_k} + \underline{\delta z} + LY_{jk} + PL_{ij} + PL_{i'j} + FL_{ii'j}$ | E10 | **I** | **I** | $\kappa$ (eBLUP) |
| $\underline{PY_{ik}} + \underline{PY_{i'k}} + \underline{FY_{ii'k}} + \underline{PLY_{ijk}} + \underline{PLY_{i'jk}} + \underline{FLY_{ii'jk}} + \epsilon_{ijk}$ | E10G | **G<sub>M</sub>** | **G<sub>M</sub>** | $\kappa$ (GEBV) |
| $\hat{y}_{ijk} = \underline{P_i} + \underline{P_{i'}} + \underline{F_{ii'}} + L_j + \underline{Y_k} + \underline{\delta z} + LY_{jk} + PL_{ij} + PL_{i'j} + FL_{ii'j}$ | E11 | - | - | $\hat{P}_{i,i'}$ (eBLUE) |
| $\underline{PY_{ik}} + \underline{PY_{i'k}} + \underline{FY_{ii'k}} + \underline{PLY_{ijk}} + \underline{PLY_{i'jk}} + \underline{FLY_{ii'jk}} + \epsilon_{ijk}$ | | | | |
| $\hat{y}_{ijk} = \underline{\beta_f w_{ii'}} + F_{ii'} + L_j + \underline{Y_k} + \underline{\delta z} + LY_{jk} + PL_{ij} + PL_{i'j} + FL_{ii'j}$ | E12 | - | - | Direct<sup>c</sup> |
| $\underline{PY_{ik}} + \underline{PY_{i'k}} + \underline{FY_{ii'k}} + \underline{PLY_{ijk}} + \underline{PLY_{i'jk}} + \underline{FLY_{ii'jk}} + \epsilon_{ijk}$ | | | | |
| $\hat{y}_{ih} = \underline{G_i} + E_h + \underline{EG_{ih}}$ (1) | E13 | **I** | **I** | $\hat{P}_{i,i'}$ (eBLUE) |
| $\hat{y}_{ih} = \underline{P_i} + \underline{P_{i'}} + F_{ii'} + \underline{\lambda_E \hat{E}_h} + P\hat{E}_{ih} + P\hat{E}_{i'h} + F\hat{E}_{ii'h} + \epsilon_{ih}$ (2) | | | | |
| $\hat{y}_{ih} = \underline{G_i} + E_h + \underline{EG_{ih}}$ (1) | E14 | - | - | Direct |
| $\hat{y}_{ih} = \underline{\beta_f w_{ii'}} + F_{ii'} + \underline{\lambda_E \hat{E}_h} + F\hat{E}_{ii'h} + \epsilon_{ih}$ (2) | | | | |

<sup>a</sup> $P_i$ and $P_{i'}$ are the general combining ability effects of pureline parents used in crossing blocks, and $F_{ii'}$ stands for specific combining ability effects. Model terms $G$ , $L$ , $Y$ , and $E$ , are the main effects of genotypes (lines), locations, years, and environments ( $h = 1, \dots, H$ ), respectively. The response variables $\hat{y}_{ijk}$ and $\hat{y}_{ih}$ are the vectors of eBLUE values of genotypic means. The covariates $z$ and $t_k$ , with regression coefficients $\delta$ and $\beta_t$ , are the mapping variables of non-genetic trend linked to the frequency of locations (Equation 2), and a continuous covariate for the calendar year, respectively. The covariate $\hat{E}$ is the predicted eBLUP values of $E$ . The residual $\epsilon_{ijk}$ ( $\epsilon_{ih}$ ) was included when stage-wise models were not used. The numbers in parentheses (1, 2) in the model's description represent steps to fit the models. Models in step one (1) were adjusted only with check varieties.
<sup>b</sup> VCOV modeled for $P_i$ and $P_{i'}$ were identity (**I**) and the additive genomic relationship matrix (**G<sub>M</sub>**).
<sup>c</sup> RGG is directly estimated from $\beta_f w_{ii'}$ , where $\beta_f$ is the regression coefficient of the genetic trend for parental lines, and $w_{ii'}$ is the year parents were used in crossing blocks (i.e., the $F_1$ was generated).

#### 4.2.3.1 Estimation models with crossing information

These models represent the inclusion of a type of parents’ general (*P_i_*) and specific (*F_ii_*′) combining abilities to estimate RGG. We say it is a “type of combining ability” because diallel mating designs were not used in the simulator. Also, because only data from BT3 in the next cycle of development is available, we can actually estimate combining abilities of breeding lines to produce the best 0.4 percent of lines created for the next cycle of line development. The analytical models are the following: Model E10 estimates *P_i_* as random effects (Mö hring *et al*. 2011), whereas Model E11 as fixed effects. Model E10G included G_M_. Shrunken or GEBV’s were then calculated as 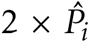 (Isik *et al*. 2017) and used to indirectly estimate RGG. For Models E10 and E10G, RGG was calculated from ***κ***_(*s*)_. Model E12 obtained direct estimates of RGG by replacing the main effects of breeding lines by *^βf^*(*s*)*^wi^*(*s*), where *β_f_*(*s*) is the RGG regression coefficient, and *w_i_*(*s*) the year the *i^th^* breeding line was used for crossing. Note only one term *β_f_*_(*s*)_ *w_i_*_(*s*)_ was included in this model given breeding lines were not used for crosses in multiple years, so that *w_i_*_(*s*)_ is equivalent for *P_i_*and *P_i_*′ . Models E13 and E14 are similar to E7 and E8 (Table 6), respectively, where the check cultivars were used to predict environmental effects in the first step (Table 7).

### 4.2.4 Statistical comparisons of analysis models

#### 4.2.4.1 Covariance modeling

Models E0, E1, *. . .*, and E6, considered GEI independent effects with homogeneous variance, whereas their counterparts E0V, E1V, *. . .*, and E6V, assumed correlated effects with heterogeneous variances. We hypothesize there is no substantial difference in estimating RGG due to variance-covariance (VCOV) modeling. We formally tested our hypothesis by performing an analysis of variance (ANOVA) of estimated RGG as a function of the simulated breeding program (i.e., simulation run) and evaluated models (Figure A6). The average estimated RGG (i.e., the slope *β_R_*) of each model (*β_RE_*_0_, *β_RE_*_0*V*_, *. . .*, *β_RE_*_6_, *β_RE_*_6*V*_) was computed, and the pairwise differences (*β_RE_*_0_ − *β_RE_*_0*V*_, *β_RE_*_1_ − *β_RE_*_1*V*_, *. . .*, *β_RE_*_6_ − *β_RE_*_6*V*_) tested with the Tukey method with *α* = 0.05 adjusted for multiplicity (Lenth 2022, functions *lstrends* and *pairs*). The same procedure was applied to compare independent versus correlated genotypic effects, and direct versus indirect estimates of RGG.

#### 4.2.4.2 Bias and linearity

Assuming 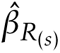 (Equation 4) is an estimator of *β_T_*_(*s*)_ (Equation 3), the estimation bias or error is defined as 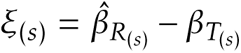. Models were evaluated based on the average 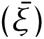 and relative 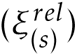 bias across simulation runs, as well as the root mean squared error (*RMSE*), computed as follows:

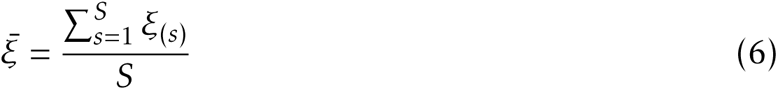

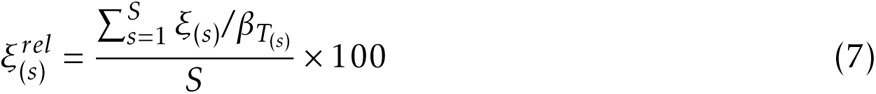

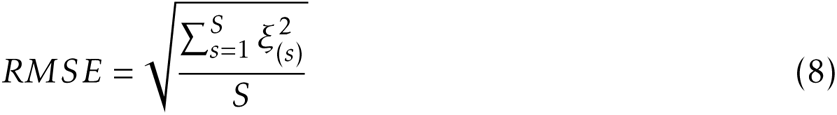

The same criteria hold for 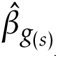, the direct estimator of RGG. In addition to bias and RMSE, linearity was accessed on the premise that if there is continuous genetic progress due to the selection of a (additive) quantitative trait, a linear trend between years of line development operation and genotypic means in MET should be evident. For indirect models where marginal genotypic (line) effects are estimated/predicted, we formally tested for linearity with to the sieve-bootstrap version of the Student’s t-test (Lyubchich and R. Gel 2022; Noguchi *et al*. 2011). The null hypothesis of no trend versus the alternative hypothesis of a linear trend was assessed on *α* = 0.05 with Bonferroni correction, and the proportion of statistically significant trends across simulation runs was reported. For direct models, we report the distribution of the two-tailed p-values from the estimated z-ratio (point estimate / standard error) of the slope (*β_g_, β_t_, β_f_*) as an indication of its significance. This distribution was compared to the random sample scenario (B2-R).

#### 4.2.4.3 Expected bias of evaluated analytic models

Estimating RGG from advanced MET without information from lines used in crosses (Section 4.2.2) is biased by design in most breeding programs. In our simulations, all experimental lines selected for recycling in BT3 advanced to PYT (Figure 1). However, not all experimental lines in PYT (or URT) were selected for crosses from the BT3. Thus, when pedigree information is not available, the estimated RGG from advanced trials is a mixture of experimental lines used for crossing, and experimental lines that were not used. In this case, if phenotypic data equal genotypic data (i.e., no error or GEI), the expected bias is defined as the difference between simulated RGG (*β_T_*_(*s*)_) and the true trend from MET 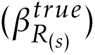. The regression slope 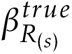 was calculated from Equation 4 by replacing ĝ*_i_*_(*s*)_ by true genetic values. Results from the expected bias are reported with the acronym “Expected”. Note the expected bias is zero for Models E10, *. . .*, E14 (Table 7), where the pedigree is known and RGG was estimated from breeding lines.

## 4.3 Estimation of RGG from empirical data

Historical soybean seed yield data from advanced MET evaluated from 1989 to 2019 was analyzed. The dataset contains 39,006 data points, 4,257 experimental genotypes derived from multiple public breeding programs, and 591 environments in the United States and Canada, for maturity groups II and III. We report results considering all experimental lines were derived from a single line development project. The dataset can be obtained from the R package SoyURT. Refer to Krause *et al*. (2022) for more details.

## 5 Results

### 5.1 Simulation overview

In total, ∼1.03 trillion data points were simulated in MET across 1,350 breeding programs. The number of different sampled locations ranged from 28 to 41 across all simulations. The sample size (i.e., the number of experimental lines excluding checks) in each breeding program ranged from 5,000-18,723 in BT1; 500-1,872 in BT2; 100-374 in BT3; 21-75 in PYT; and 4-15 in URT. Trial-level estimates of heritabilities were consistent with the type of trial (Figure A7). REML estimates of variances are also reported (Figure A8). The average true simulated RGG (bu/ac^−1^/yr^−1^) was 0.44 for A1 and A2, 0.41 for B1 and B2, 0.55 for B2-M, and zero for B2-R (Figure 3, “Simulated”).

**Figure 3:**
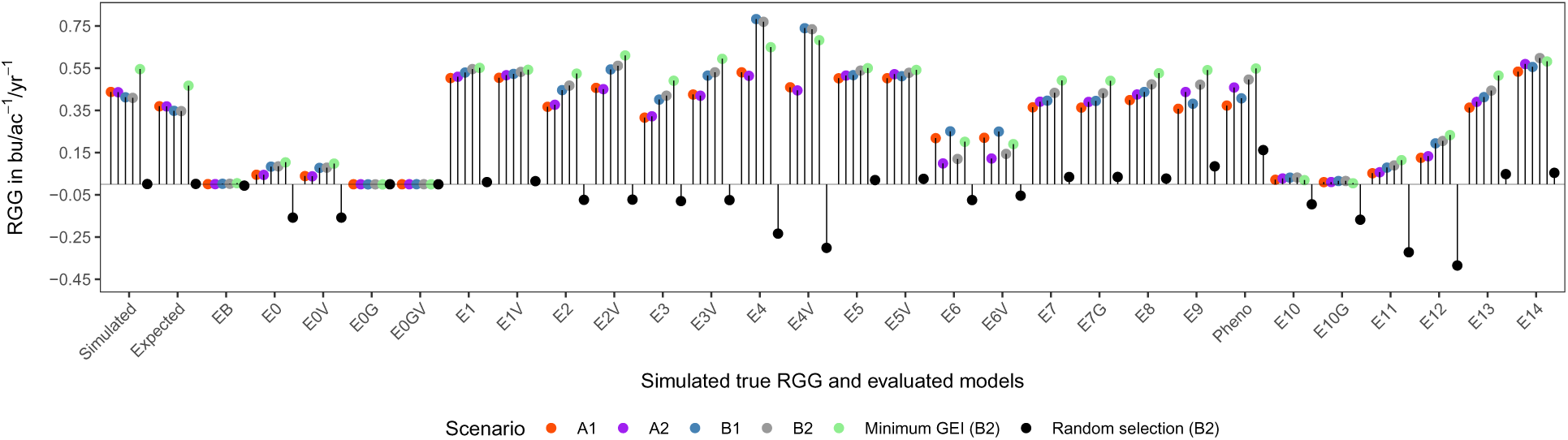
Average true simulated 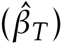, expected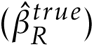, and estimated RGG 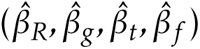.

### 5.2 Covariance modeling

Across all simulation models, the inclusion of covariance models for GEI significantly affected contrasts of estimated RGG values. Three contrasts were statistically significant when a random sample was used to create a new cycle of breeding (B2-R). The inclusion of the genomic relationship matrix (G_M_) was statistically significant for the contrast E10-E10G in B2-R, and E0-E0G was significant for all B simulation models. The direct versus indirect estimation of RGG resulted in significant differences between estimated RGG values for most simulation models; the primary exception was that the contrast between E7 and E8 was not large for any of the simulation models (Table 8).

**Table 8:** Adjusted Tukey P-values for the contrasts between variance-covariance modeling (VCOV) and direct versus indirect (DI) estimation of RGG.

| Contrast | Simulation models (scenarios) |  |  |  |  |  | Type |
| --- | --- | --- | --- | --- | --- | --- | --- |
|  | A1 | A2 | B1 | B2 | B2-M | B2-R |  |
| E0 - E0V | 1.00 | 1.00 | 1.00 | 1.00 | 1.00 | 1.00 | VCOV |
| E0 - E0G | 0.48 | 0.67 | < 0.01 | < 0.01 | < 0.01 | < 0.01 | VCOV |
| E1 - E1V | 1.00 | 1.00 | 1.00 | 1.00 | 1.00 | 1.00 | VCOV |
| E2 - E2V | < 0.01 | < 0.01 | < 0.01 | < 0.01 | < 0.01 | 1.00 | VCOV |
| E3 - E3V | < 0.01 | < 0.01 | < 0.01 | < 0.01 | < 0.01 | 1.00 | VCOV |
| E4 - E4V | < 0.01 | < 0.01 | 0.71 | 0.98 | 0.99 | 0.01 | VCOV |
| E5 - E5V | 1.00 | 1.00 | 1.00 | 1.00 | 1.00 | 1.00 | VCOV |
| E6 - E6V | 1.00 | 1.00 | 1.00 | 1.00 | 1.00 | 1.00 | VCOV |
| E7 - E7G | 1.00 | 1.00 | 1.00 | 1.00 | 1.00 | 1.00 | VCOV |
| E10 - E10G | 1.00 | 1.00 | 1.00 | 1.00 | 1.00 | < 0.01 | VCOV |
| E1 - E2 | 0.00 | 0.00 | 0.00 | 0.01 | 1.00 | 0.00 | DI |
| E1V - E2V | 0.35 | 0.03 | 1.00 | 1.00 | 0.03 | 0.00 | DI |
| E7 - E8 | 0.94 | 0.97 | 0.80 | 0.92 | 0.98 | 1.00 | DI |
| E11 - E12 | 0.00 | 0.00 | 0.00 | 0.00 | 0.00 | 0.00 | DI |
| E13 - E14 | 0.00 | 0.00 | 0.00 | 0.00 | 0.04 | 0.00 | DI |

### 5.3 Relative bias and overall performance

Analytic Models EB, E0G, E0GV, and E10G, were not able to estimate RGG (Figure 3). Results from these models and from B2-R will not be included in the reported summary statistics below. Raw estimates of RGG are available in Figures A9, A10, A11, A12, A13, and A14, as well as the estimated bias without standardization in Figure A15.

All models demonstrated some degree of bias. On average, a smaller bias was observed for simulation models with complex GEI effects. The expected relative bias for models without information from breeding lines had an average value of -15.21% (Figures 4 and A16). Models with information from breeding lines are expected to have no bias, but did not outperform models that only considered advanced trials (Figures 3, 4, 5). Including G_M_ to account for correlated genetic effects did not improve the estimates of RGG. Across simulation runs, the average value of the diagonal values of G_M_ was 1.91, and zero for the off-diagonal. Within years, these values were 1.91 and 0.49, respectively (data not shown).

**Figure 5:**
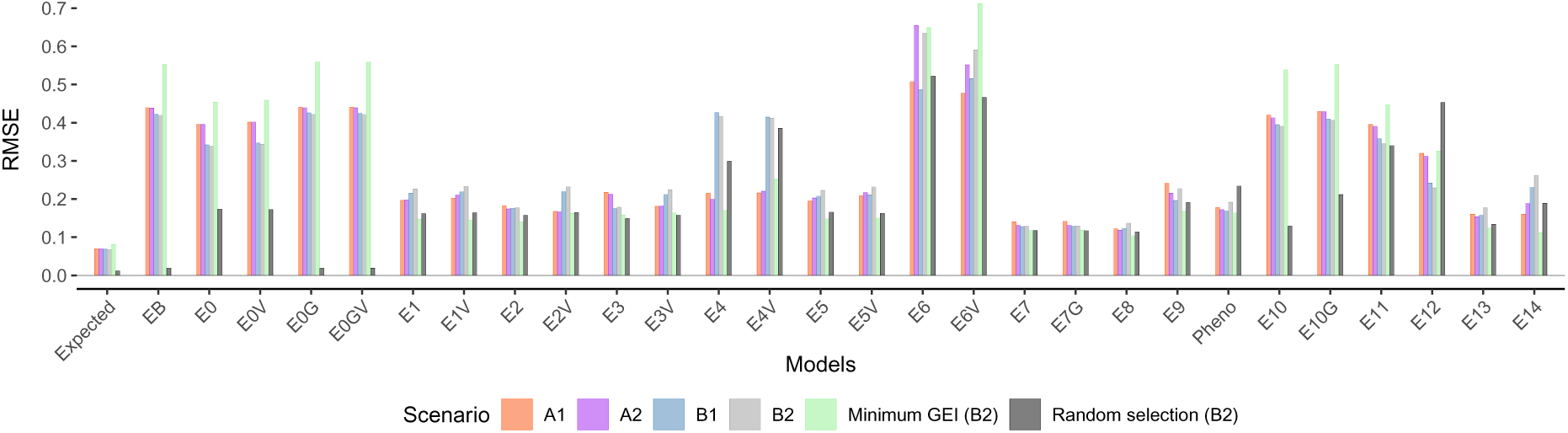
Root mean squared error (RMSE).

On average, 15 models presented less than ±5% relative bias for at least one simulation model. Models E2, E7, E7G, E8, and E13 had less than ±18% relative bias across all simulation models. Directional relative bias (i.e., under or overestimating true RGG on average) was observed for Models E1, E2V, E4, E4V, E5, E6, E6V, E10, E11, E12, and E14, and very similarly for Models E1V, E3, E3V, and E5V (Figure 4). Across simulation models and excluding very few negative estimates (Table A1), the range of estimated RGG from Models E1, E2V, and E7, contained the true simulated RGG in 58% of the simulations (Figure A17). For these models, the relative bias ranged from -16.82% to 40.73%, which represents biases of ±7.41 kg/ha^−1^/yr^−1^ or ±0.11 bu/ac^−1^/yr^−1^.

Estimates of RGG using raw phenotypic data (“Pheno”) resulted in relative bias across simulations from -14.64% to 23.23% (Figure 4), with a similar *RMSE* as other analytic models (Figure 5). However, in B2-R, where there is no RGG, the average estimated gain was 0.16 bu/ac^−1^/yr^−1^ (Figure 3). Simulation models A2, B2, B2-M, and B2-R, included a positive non-genetic trend. Although in this work we are not investigating the estimation of the rate of non-genetic gain (i.e., we were trying to isolate it from RGG), by comparing the relative bias in scenarios A1 versus A2, and B1 versus B2, it is evident some analytic models successfully isolated it (Figures 3 and 4).

### 5.4 Linearity

For Models E0V, E1, E1V, E7, and E7G (advanced MET), the proportion of statistically significant linearity ranged from 0.80 to 0.99, with an average value of 0.90. As expected, true simulated genetic values from breeding lines were always statistically significant. In B2-R, Models E0, E0V, E4, E4V, E5, E5V, E10G, and E11, presented an average value of 0.63, similar to when raw phenotypic values were considered (Figure 6). For models in which RGG was directly estimated, the distribution of the p-values from the z-ratio statistic showed it could be used to indicate there was RGG, except for Models E6, E6V, and E12 (Figure 7).

**Figure 6:**
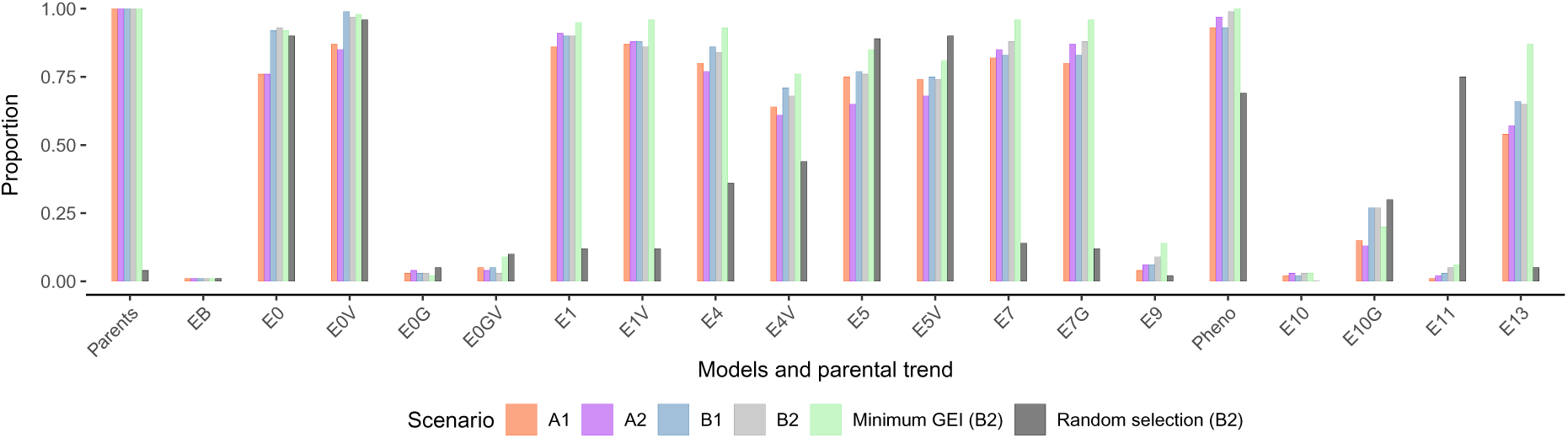
The proportion of statistically significant linear trends for models that indirectly estimate RGG.

**Figure 7:**
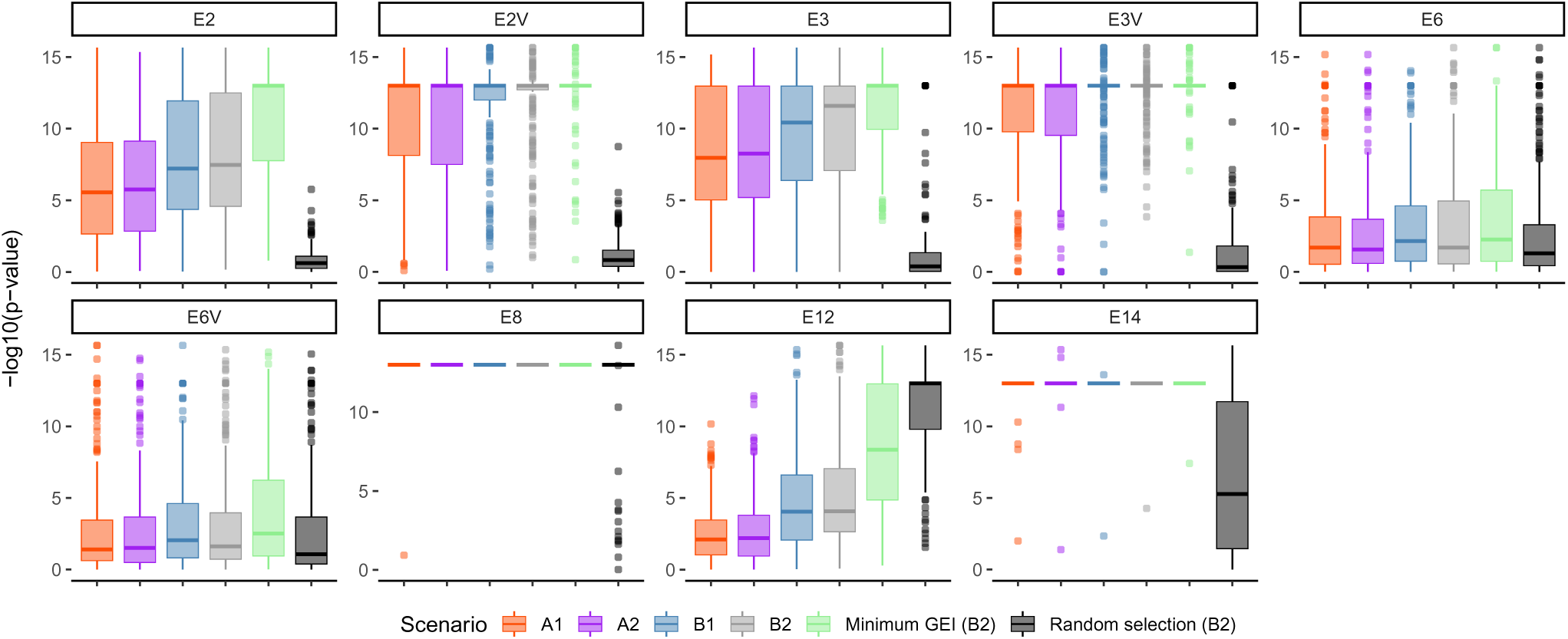
Distribution of the -log10(p-values) from the z-ratio statistic of direct RGG estimation.

### 5.5 Estimates of RGG from empirical data

Based on the simulation results, we used models E1, E2V, and E7 to estimate RGG for the empirical data. For Model E2V, several covariance structures were evaluated, and the best-fit model was selected (Table A2). The point estimates of RGG ranged from 0.27 to 0.59 bu/ac^−1^/yr^−1^. For Models E1 and E7, the p-values from the linearity test were statistically significant, as well as for the z-ratio of the estimated slope *β_g_* (i.e., the direct estimate of RGG) from Model E2V (Table 9). Therefore, although imprecise, there is strong evidence that RGG from public soybean breeding programs has been positive for maturity groups II and III in the period from 1989 to 2018.

**Table 9:** Estimated RGG with 95% confidence interval (bu/ac^−1^/yr^−1^) and linearity for the empirical soybean dataset.

| Model | RGG | Linearity |
| --- | --- | --- |
| E1 | 0.59 (0.58, 0.61) | < 0.01 |
| E2V | 0.27 (0.20, 0.35) | < 0.01 |
| E7 | 0.42 (0.40, 0.43) | < 0.01 |

## 6 Discussion

Analytical methods to estimate RGG have been a long-term challenge to animal and plant breeders (Rizzo *et al*. 2022; Garrick 2010; Eberhart 1964). In the plant breeding community, several reasons underlie its adoption, such as the need to evaluate novel breeding strategies, and as a metric to quantify the programs’ efficiency (Rutkoski 2019a,b). Herein, we proposed RGG be defined as the accumulation of beneficial alleles in breeding (parental) lines across years of breeding operation. This definition considers the intricate nature of cultivar development programs while remaining consistent with the original concept of genetic gain (Falconer 1960; Kempthorne 1957).

In parallel to recurrent selection (Jenkins 1940), cultivar development programs also consist of evaluation, selection, and reproduction; however, there are relevant distinctions. There is no single cycle zero from which all genotypes are recurrently derived. Indeed, within both public and commercial plant breeding programs, there are many geographically distributed breeding projects that began with unique sets of founders, and there is an exchange of lines across projects within the programs. Consequently, genetic changes are due not only to selection, but also to migration and drift. In this work, we addressed RGG in a closed system, i.e., there was no exchange or introgression of external germplasm. If a line from an external breeding program is included in the crossing nursery, then our RGG definition still applies as long as an adjusted genotypic mean (i.e., eBLUE value) is provided and included in the RGG estimation.

Distinguishing RGG from genotypic and/or commercial gain will help avoid misleading interpretations. Genotypic gain is the RGG plus the expression of non-additive genetic effects such as epistatic (Pavlicev *et al*. 2010; Hansen and Wagner 2001) and dominance deviations. Hence, for a trait not only controlled by additive effects (e.g., Garcia *et al*. 2008), the genotypic gain is expected to be higher than the RGG. Commercial gain is the genotypic gain delivered in the farmer’s field. Examples of commercial gain are estimates from era trials (e.g., Cooper *et al*. 2020; Bruce *et al*. 2019) when only farmer-grown cultivars are considered. The essential distinction between RGG, genotypic, and commercial gain, is that only RGG is relevant to genetic improvement by the breeding project. Also, both genotypic and commercial gains are related to the concept of commercial heterosis, in which the performance of the released hybrid is compared over a commercial check cultivar (Labroo *et al*. 2021).

Another metric called “yield gain, advances”, or simply “genetic trend”, has been used to quantify the driving factors of yield increase of staple crops worldwide (Prasanna *et al*. 2022; Grassini *et al*. 2013) or in specific countries (Rizzo *et al*. 2022; Guo *et al*. 2022; Fischer *et al*. 2022). For example, Rizzo *et al*. (2022) collected maize field-trial data over 14 years (2005-2018) from the state of Nebraska (USA), and concluded that climate and agronomy represented 87% of the yield gains in high-yield irrigated environments, leaving 13% for the genetic contribution. The 13% genetic contribution can be interpreted as an unweighted average of the commercial gain. It is unweighted in the sense that individual commercial programs breeding for Nebraska have a specific contribution to the reported gain, given each program has released a number of hybrids with an average lifespan of three years according to their market share. For that reason, the yearly yield gain is a composition of commercial gain within and across breeding programs. Consequently, careful consideration should be given when linking reported genetic trends with RGG.

The main emphasis of this study was to estimate RGG from advanced MET. We choose linear mixed models for estimators because they are commonly used in data analysis of MET (Krause *et al*. 2020; Dias *et al*. 2018; Isik *et al*. 2017) and are well-known in the plant and animal breeding communities. The underlying distributional assumptions of linear mixed models are that random effects have an expected value of zero, and are realizations of independent and multivariate Gaussian distributions with positive-definite variance matrices (Gumedze and Dunne 2011; Henderson *et al*. 1959; Henderson 1950, 1949). By assuming the random effects have an expected value of zero, both the average (Isik *et al*. 2017) and sum (Searle 1997) of predicted values for the random factor are zero. This analytic constraint, however, does not assure the eBLUP value of a newly developed genotype will be numerically higher than that of an older genotype when historical MET data is analyzed using G×L×Y models (Figure A21A). Thus, our benchmark Model EB did not provide positive values for RGG.

An alternative parametrization for the G×L×Y benchmark model was computed with the cumulative sum of the average eBLUP values of all genotypes according to the year of first testing, (e.g., the PYT for the available public soybean breeding data and our simulations). This strategy is connected to the expected gain from selection when all genotypes that are under selection are analyzed together (i.e., the same model). In this case, the expected/predicted gain can be directly calculated with the average of the eBLUP values from selected individuals. The rationale is that, when likelihood-based estimators of variance components are used (e.g., REML), the regularization parameter lambda in the BLUP predictor is the ratio of residual and random term variance estimates. Hence, for the main effects of genotypes, the shrinkage of the genetic term is inversely proportional to the heritability/repeatability of the trait (Xavier *et al*. 2016). The practical application of ***κ*** relies on the assumption that the observed experimental genotypes in MET are a realization of a multivariate random variable with covariance depending on genetic relationships. This assumption does not require reference to a specific base population with idealized properties (Piepho *et al*. 2008). Furthermore, in parallel to recurrent selection, the first cycle is arbitrarily set by the first available year in the MET data, thus representing the initial value of ***κ***.

A second alternative parameterization of benchmark model G×L×Y was to model the combination of location and years, i.e., environments. The idea underlying this approach is from Diers *et al*. (2018) and Montes *et al*. (2022). These authors analyzed phenotypic data from a soybean nested association panel using stage-wise models. In the first-stage model, the eBLUP values of incomplete blocks within locations were predicted using a unique identifier in the dataset, and in the second-stage model, were used as a fixed covariate. We modified their first-stage model to obtain the eBLUP values of environmental effects 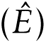 based on evaluations of check cultivars, which were then considered a fixed covariate to represent both E and GEI effects. We hypothesized that if environmental effects can be successfully captured by check cultivars in the first-stage (Figure A22), then unbiased estimates of genetic effects could be obtained from the second-stage model.

While this second modeling strategy successfully captured RGG and GEI effects, it significantly inflates the estimates of genotypic variance, and consequently, the estimates of heritability (Figures A21B and A23). For example, estimates of genotypic variance and heritability using empirical soybean MET data changed respectively from 6.30 (bu/ac)^2^ and 0.47, in the benchmark Model EB, to 31.11 (bu/ac)^2^ and 0.94 with Model E7. A similar outcome can be achieved by dropping the interaction terms GL, GY, and GLY from the benchmark model, where the estimated genotypic variance changed from 6.30 (bu/ac)^2^ to 27.01 (bu/ac)^2^. These results suggest the genotypic variance in Models E7 and E7G were inflated by the GL, GY, and GLY variances associated with the experimental genotypes. Thus, while checks account for variability among environments of MET, the GEI effects from experimental genotypes are still confounded with the genotypic variability.

The aforementioned inflation due to GEI applies to other mixed models used for analyzing data from MET. For example, the “EBV” model from Rutkoski (2019b) only accounted for genotypic main effects and successfully captured RGG with a biased (inflated) genotypic variance. In this case, RGG estimates from eBLUP values 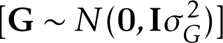, estimated breeding values 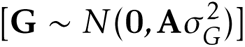, or genomic estimated breeding values 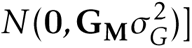, will not produce large differences, as shown in Models E7 and E7G. Nonetheless, for annual and breeding line selections, it is well-known that under normality, equal variance, and independence, BLUP will minimize the mean squared error of predicted values (Piepho *et al*. 2008; Robinson 1991) and hence maximize the correlation between true and predicted genotypic values (Searle *et al*. 1992; Searle 1974; Henderson 1963). Thus, although RGG from MET can not be directly estimated from eBLUP values of the G×L×Y model, the use of BLUP is likely to increase gains from selection if data is correctly modeled (Hartung *et al*. 2023; Smith *et al*. 2005).

Every model assessed to estimate RGG in the simulations demonstrated some degree of bias. For example, the average bias was similar between simulation models A1 and A2 (simple GEI effects), and in B1 and B2 (complex GEI effects), but were relatively different between the A and B simulations. The inclusion of a positive, cumulative, non-genetic gain did not result in major bias when comparing simulation models A1 versus A2, and B1 versus B2. Given simulation results suggest bias is associated with the simulation model, ideally, RGG should be estimated with models with small values for root mean squared error. Moreover, recognizing that all evaluated models were biased, directional bias is desirable in the sense that it is consistent. Directional bias can be seen as insurance that the estimated RGG from empirical datasets is likely under or overestimated.

The observed bias was also related to the manner in which RGG was defined and to the structure of the simulated breeding program. The simulator was constructed to mimic public soybean breeding programs responsible for maturity zones II and III in the USA. These programs evaluate nearly homozygous lines in replicated field trials for four to five years (Figure 1). The available empirical and simulated data reflect the last two years of MET. We assumed for simulation that breeding lines were selected in BT3 after three years of trials, and then were used in the crossing nursery as well as advanced to a regional PYT. Thus, the PYT stage could include lines that were not used in the crossing nursery. Consequently, estimated RGG that only uses data from PYT and URT, and has no information about breeding lines, represents a biased sample of lines relative to the breeding lines that are used to create the next cycle of progeny. Our results show that the average bias was -15.21% for the simulated conditions, indicating that, on average, the samples of lines included in the PYT and URT underestimate RGG.

The sampling bias is also related to sample size. In general, a larger sample size is desirable for obtaining more precise estimates and reducing the variance of the estimator. We focused on estimating RGG from 30 years of advanced MET and could not obtain an unbiased, robust estimator. Results considering only 10 years of data revealed a much larger bias and root mean squared error (Figures A18, A19, A20), as also observed by Rutkoski (2019b). The simulated breeding programs in this work as well as from Rutkoski (2019b) assumed small breeding projects, relative to proprietary breeding projects. Proprietary breeding programs have a much larger network of trials, which could easily represent a 3 to 6-fold increase in the amount of data. A question for further investigation is what is the optimal (or minimum) number of locations and years from advanced MET needed to obtain unbiased estimates of RGG using linear mixed models.

A pattern of missing data also can introduce bias in trend (Hartung *et al*. 2023) and REML variance estimation (Hartung and Piepho 2021; Aguate *et al*. 2019; Piepho and Mohring 2006). We only considered data from two or three years of MET, excluding the information from local breeder trials. We excluded these because empirical data from local breeders trials historically have not been published nor recorded in accessible databases. Further research is needed to investigate if this exclusion introduced bias in estimates of RGG. Furthermore, simulated breeding lines were selected in BT3 with BLUP models that consider all available data (BT1, BT2, BT3). It is worthwhile investigating if these predicted values could lead to less biased estimates of RGG. This approach would also be informative regarding the predicted genetic gain as previously discussed. In addition, early generations can also be used to estimate RGG (Cowling *et al*. 2023).

Computing RGG from raw phenotypes, without statistical modeling, can indicate positive RGG when no genetic gain was actually delivered. One might argue that when the data is balanced the arithmetic average and (generalized) least-squares yield numerically equivalent estimates. The first issue is that data from MET are rarely balanced within years, and largely unbalanced across years. Thus, RGG estimates from raw phenotypes are completely confounded with non-genetic effects. As emphasized by Hartung *et al*. (2023), careful consideration should be given in selecting the best-fit, proper, model for the dataset being analyzed. Great attention should be given to evaluating covariance modeling for non-genetic effects given it played an important role for most models. Metrics such as the Akaike and Bayesian information criteria (Akaike 1974; Schwarz 1978), as well as the proportion of genetic variance explained by FA models (Smith *et al*. 2015), should always be considered.

Both direct and indirect estimators of RGG provided useful information. For the theoretical development of direct estimation see Piepho *et al*. (2014). The inclusion of additive genomic relationships to account for correlated genetic effects did not improve their estimates of RGG. This result likely occurs due to genotypes exhibiting little to no correlation across years of MET. Diallel-based models were also evaluated assuming pedigree was available, so RGG was estimated directly from breeding lines. These models did not outperform models that only considered advanced trials, and hence there is no clear advantage in considering this modeling approach. When pedigree is available, an alternative to compute RGG would be to use breeding lines *per se* performance. We did not test for this approach, but results from Rutkoski (2019b) showed unbiased estimates could not be obtained.

Overall, the best-performing models were E1, E2V, and E7. Since none of the models yielded unbiased estimates of RGG, the most suitable strategy would be to account for the range of the estimated values. This would increase the likelihood of capturing the true RGG. Using this strategy, we report estimates of RGG that range from 18.12 to 39.60 kg/ha^−1^/yr^−1^ (0.27 to 0.59 bu/ac^−1^/yr^−1^) for 30 years of empirical soybean MET. The modeling was done assuming that there is only one set of BTs from a single cultivar development project. It should be pointed out that there are actually multiple variety development projects working in maturity zones II and III, so that there are multiple sets of BTs and further work using island models (Ramasubramanian and Beavis 2021) are needed to provide better interpretation of results (including maintenance of genetic variability) from MET. Lastly, if the goal is to determine if there is evidence for RGG, regardless of bias, our linearity measure based on the simulation results is useful. Further investigation is needed if RGG is expected to be nonlinear (Ramasubramanian and Beavis 2021; Bulmer 1971; Eberhart 1964).

## 7 Conclusion

We evaluated several linear mixed models to estimate RGG using advanced, routine MET. We approach the research question by simulation and propose a novel RGG definition that considers the intricate nature of cultivar development programs while remaining consistent with the original concept of genetic gain. Our results suggest it is not possible to accurately estimate RGG using data from two years of MET, such as are available in the public soybean breeding programs. Consequently, the evaluated estimators should not be used to compare breeding programs or quantify the relative efficiencies of proposed breeding systems. If the goal is only to determine whether there was RGG, the linearity metric is useful. Therefore, as also concluded by Rutkoski (2019b), there was no ideal model to estimate RGG. Lastly, in addition to the practical and theoretical results applied to soybean genetic improvement, the analyses performed in this study can be applied to quantitative traits evaluated in any diploid crop undergoing phenotypic evaluations in MET.

## 8 Declarations

Conflicts of interest: None declared.

## 9 Author contribution statement

MDK and WDB conceived the research; MDK designed the simulator, models, performed the statistical analyses, and wrote the first drafts of the manuscript; HPP and KOGD provided insights into the methodology; HPP revised numerous drafts of the manuscript; AKS provided knowledge on the structure of a public soybean breeding programs; and WDB and AKS were responsible for acquiring funding to support the research. All authors approved the final version of the manuscript.

## 10 Data and code availability

The simulator and evaluated models are publicly available on GitHub (https://github.com/mdkrause/RGG). The soybean empirical data is available in the R package SoyURT

(https://github.com/mdkrause/SoyURT).

## 11 Acknowledgments

Our sincere thanks to Dr. R Chris Gaynor for providing R functions to simulate GEI effects with a compound symmetry model, and the Iowa State University (ISU) Research IT team for providing efficient computational resources.

## Appendices

## A Additional figures

**Figure A1:**
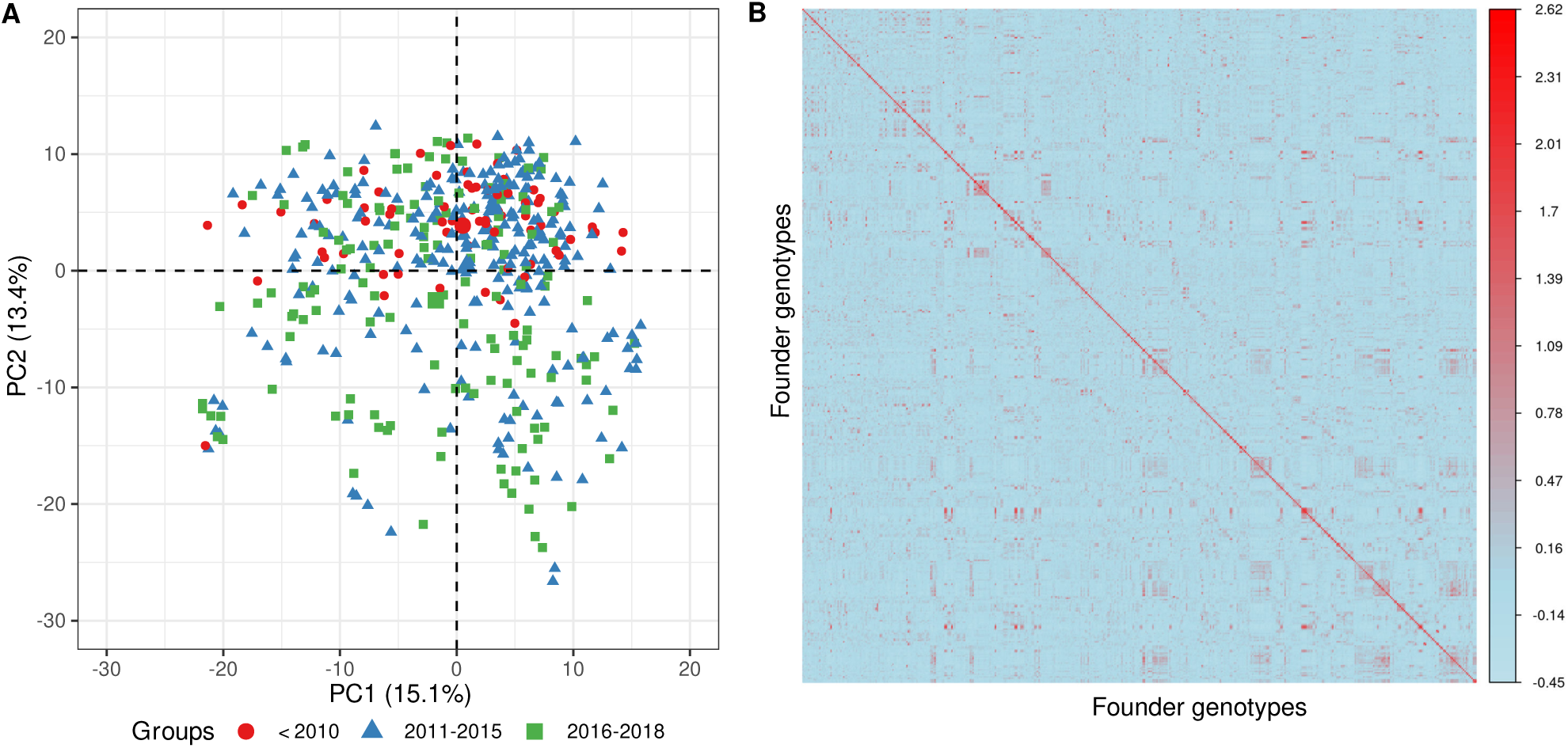
Principal component analysis of the additive genomic relationship matrix (G_M_) estimated from the population of experimental genotypes used as founders in the simulation (A), and a heatmap of the G_M_ matrix (B).

**Figure A2:**
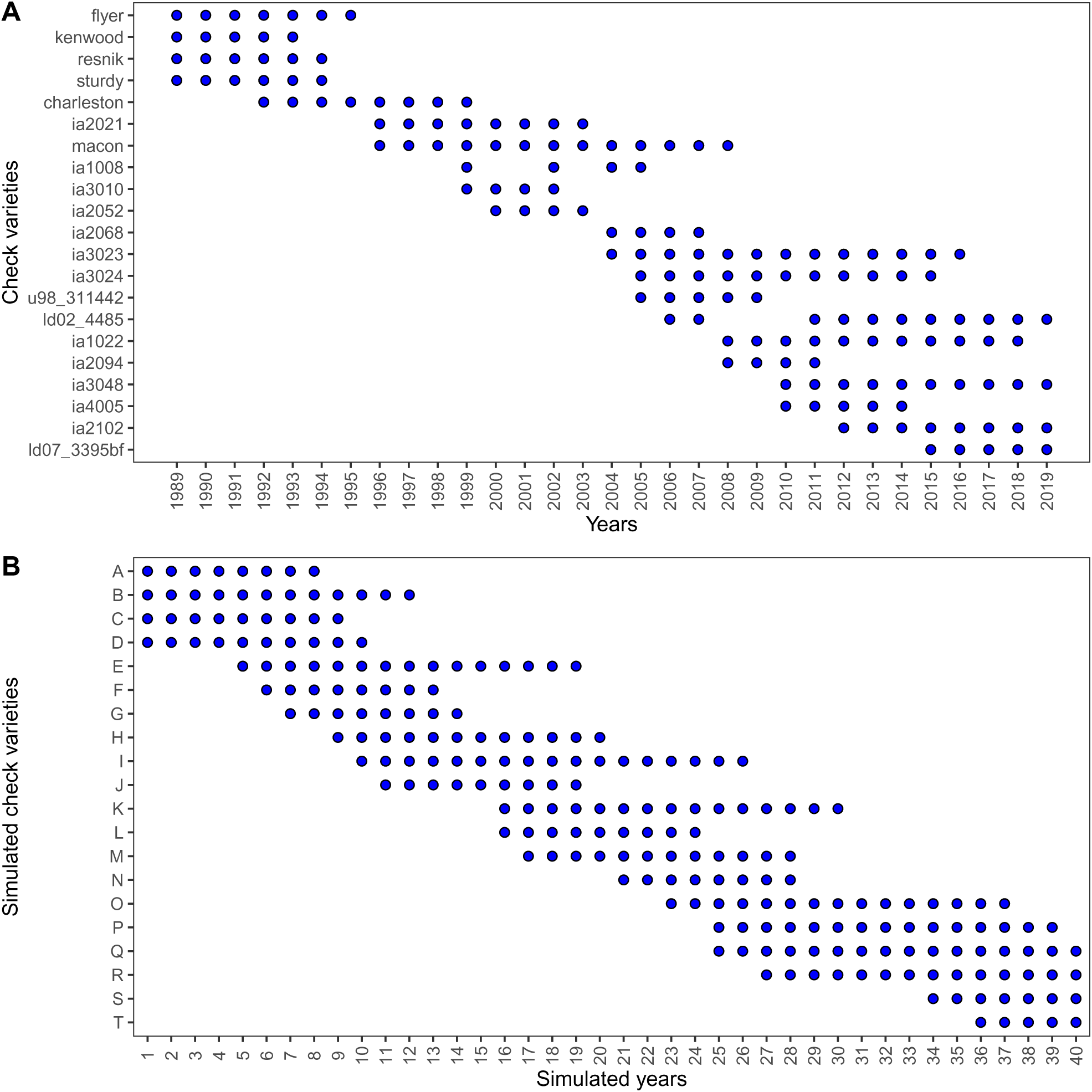
Frequency of empirical (A) and simulated (B) check cultivars.

**Figure A3:**
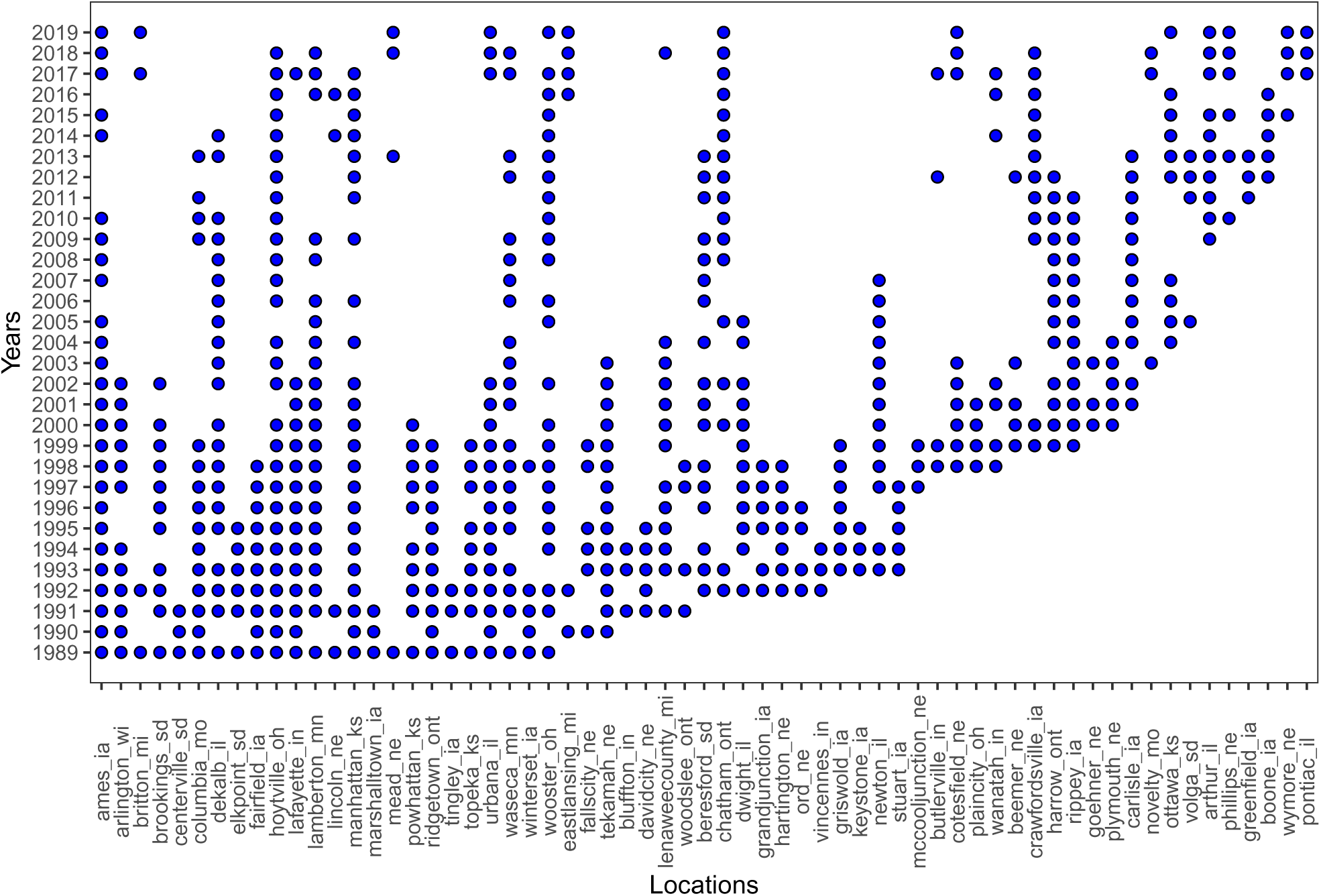
Frequency of locations from empirical data.

**Figure A4:**
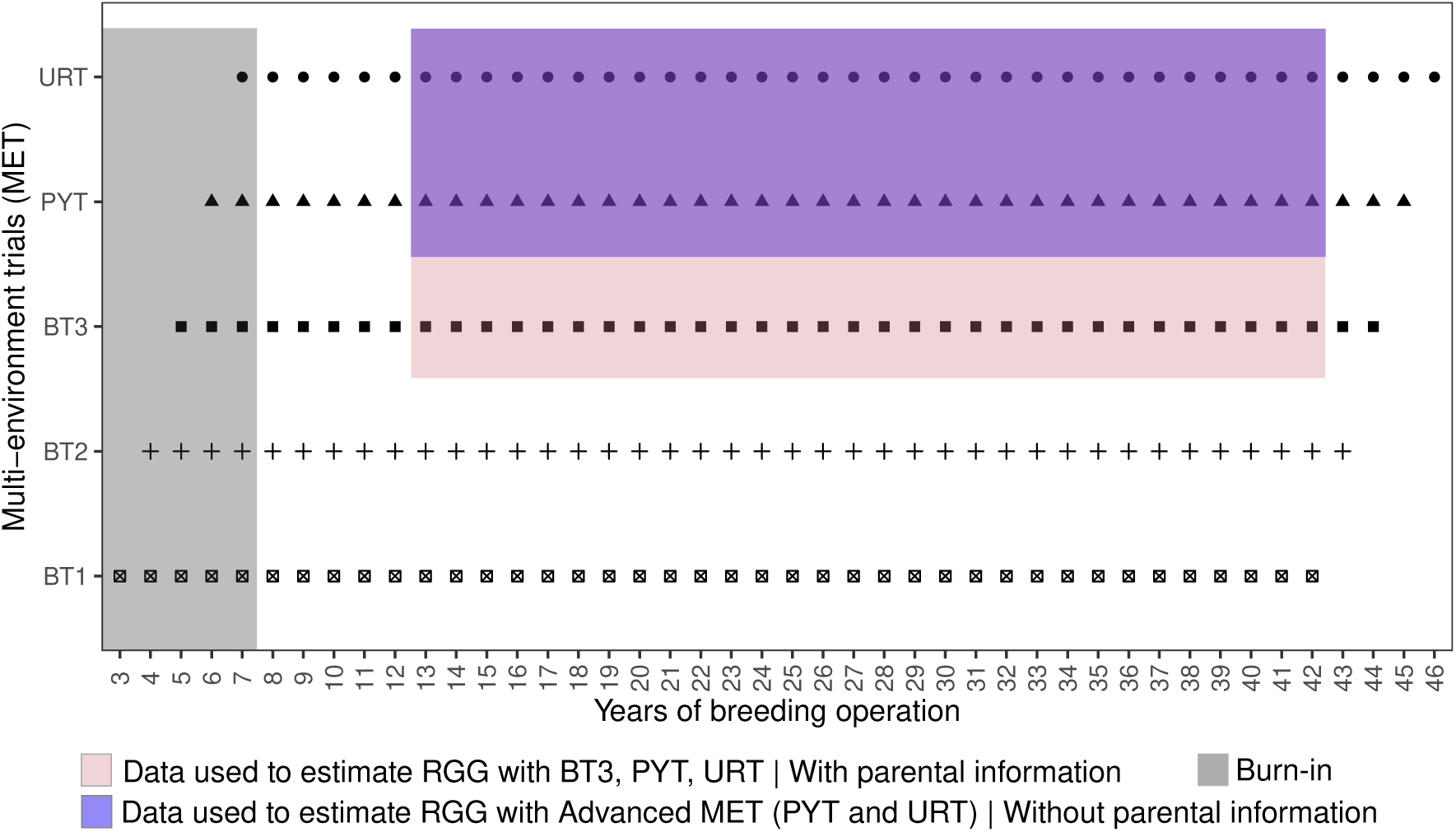
The structure of the simulated MET. Colors represent the source of data used to estimate RGG (with or without information from breeding lines).

**Figure A5:**
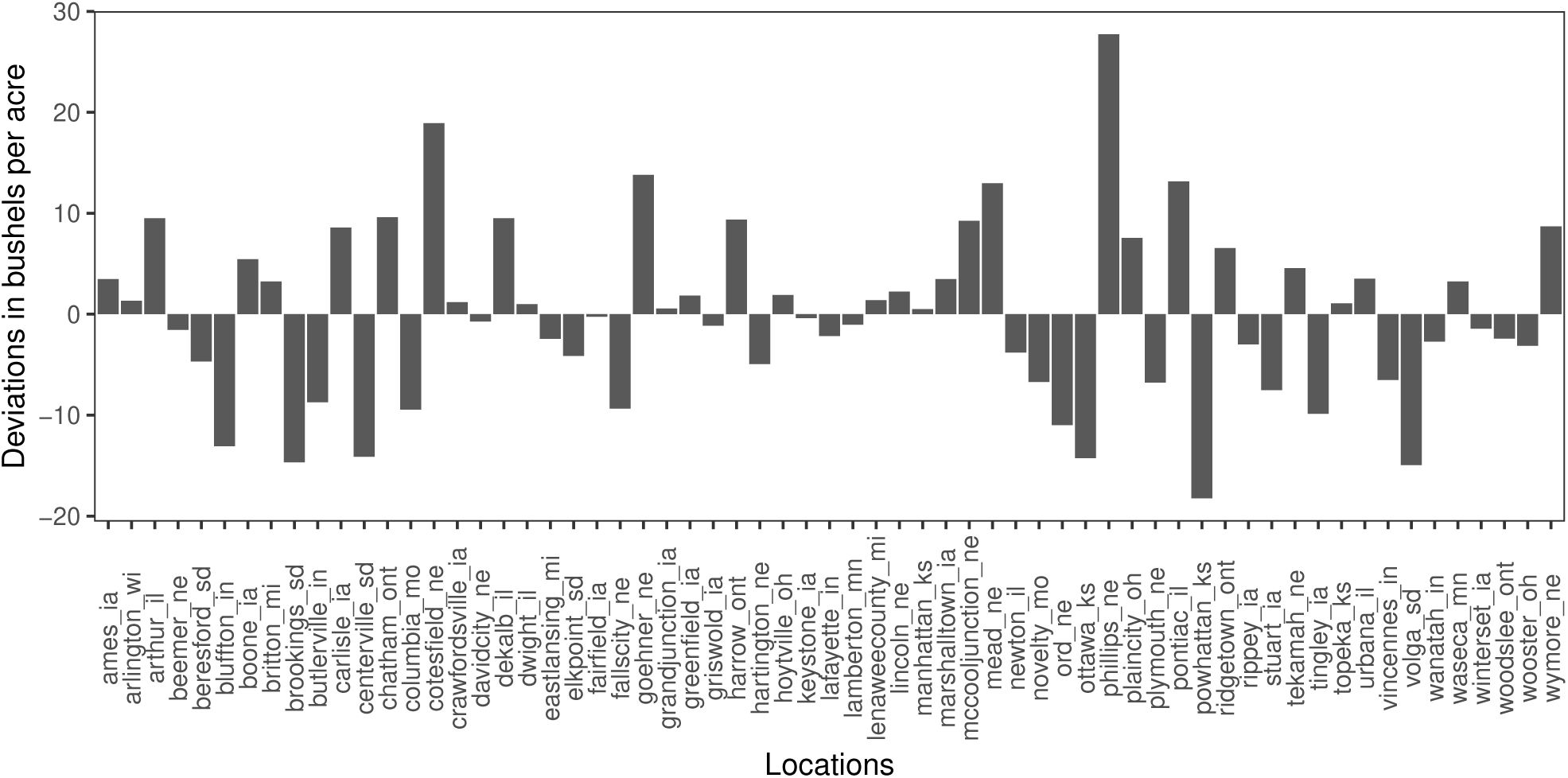
Estimated marginal location values from empirical data used in the simulator.

**Figure A6:**
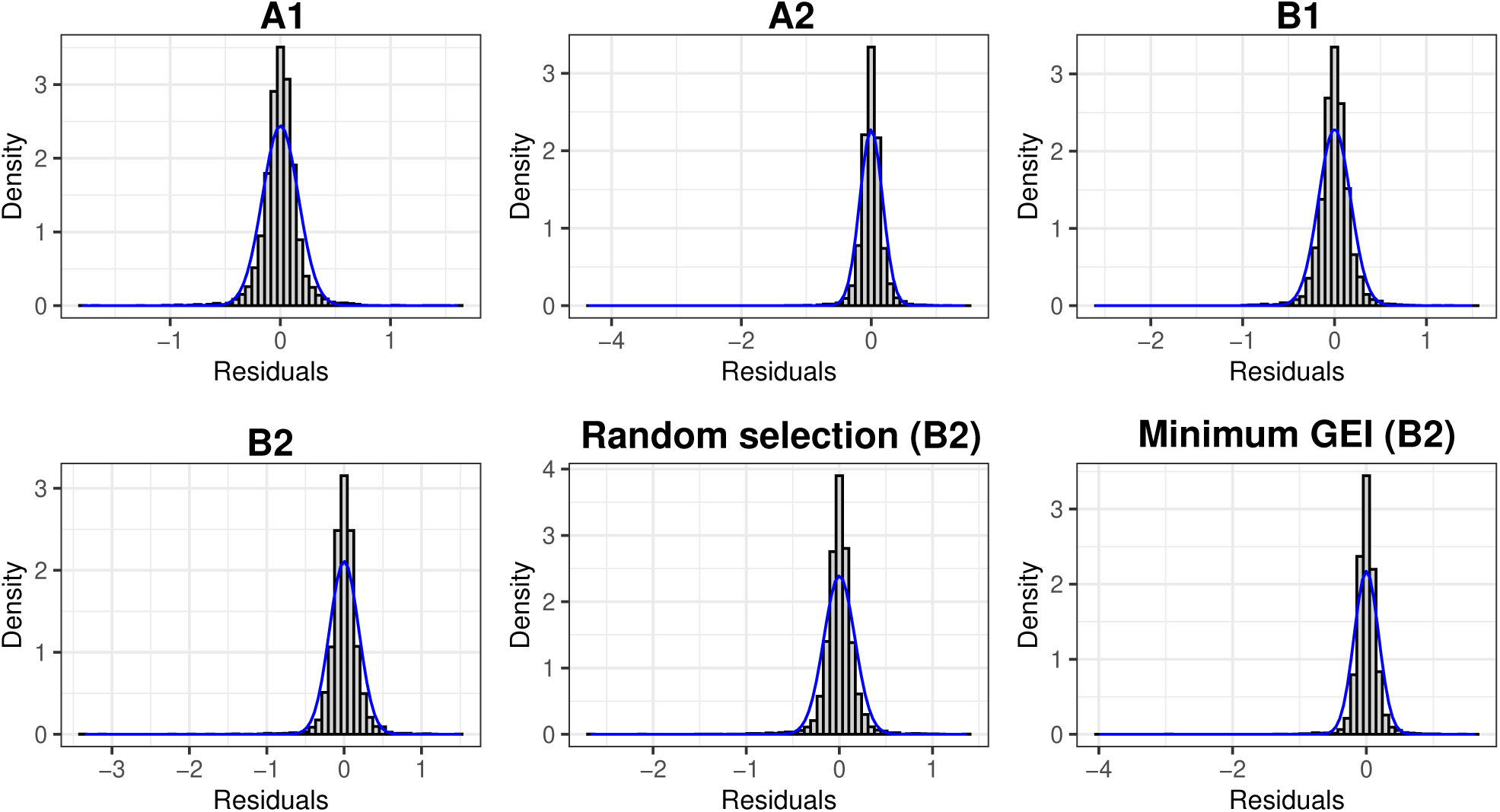
Histogram of estimated residuals from the analysis of variance performed to evaluate the average slope (RGG) of tested models (Tables 6 and 7).

**Figure A7:**
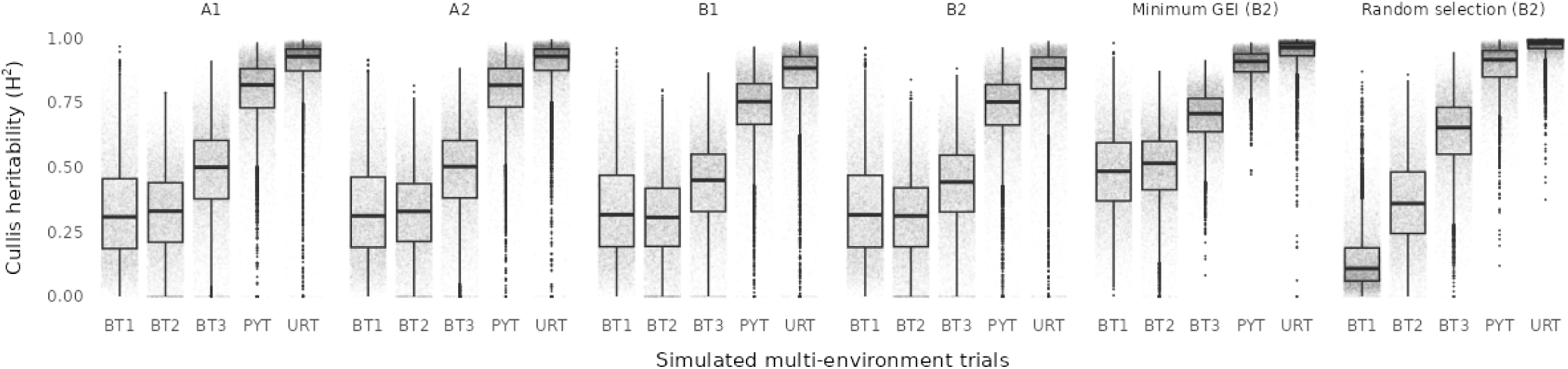
Estimates of generalized heritability (Cullis et al. 2006) for each simulation model and trial.

**Figure A8:**
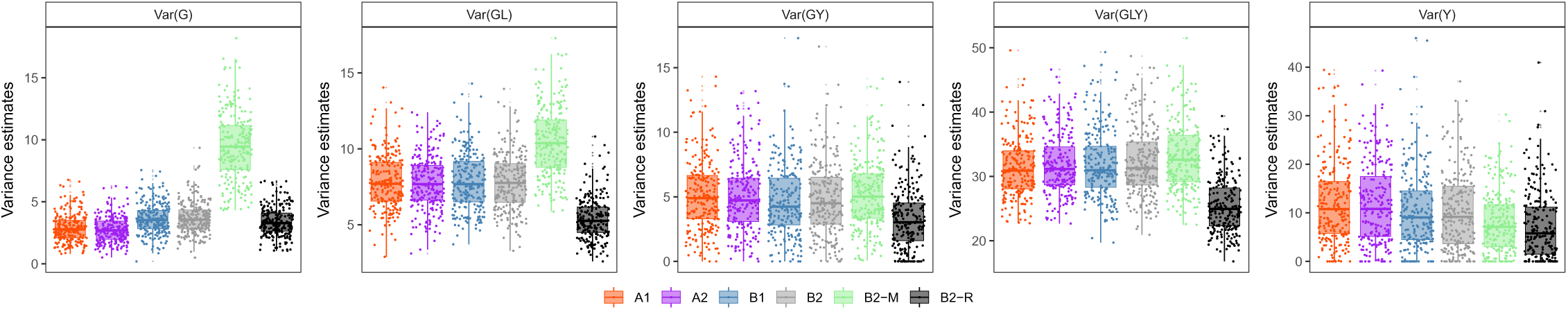
REML estimates of genotypic (G), year (Y), G × Location (L), G × Y, and G × L × Y variances for each simulation scenario. Data were analyzed with Model 1 and included BT3, PYT, and URT.

**Figure A9:**
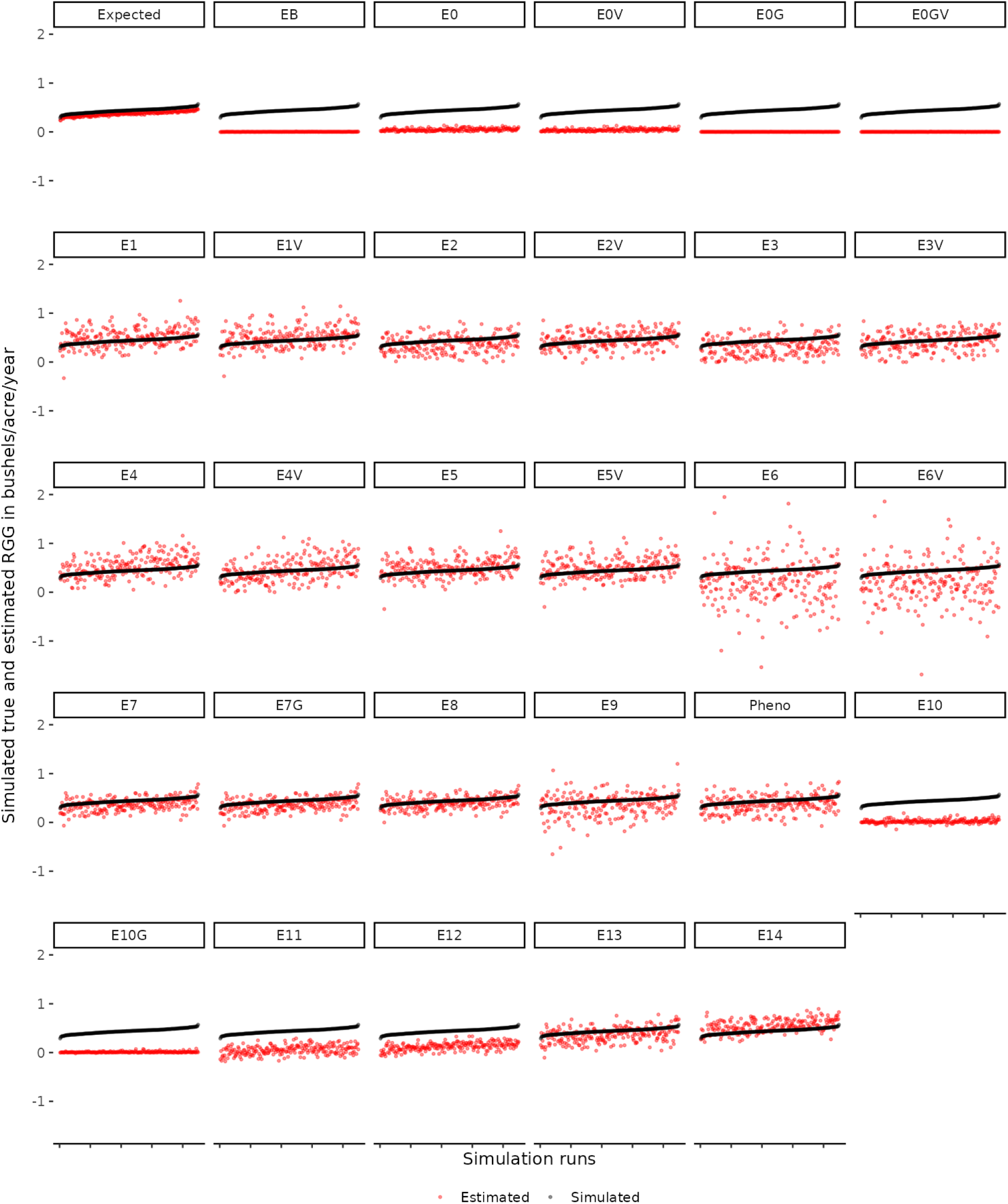
True (black) and estimated (red) RGG for evaluated models in scenario A1 (Tables 1 and 2). Check Section 4.2.4.3 for details on the “Expected” facet.

**Figure A10:**
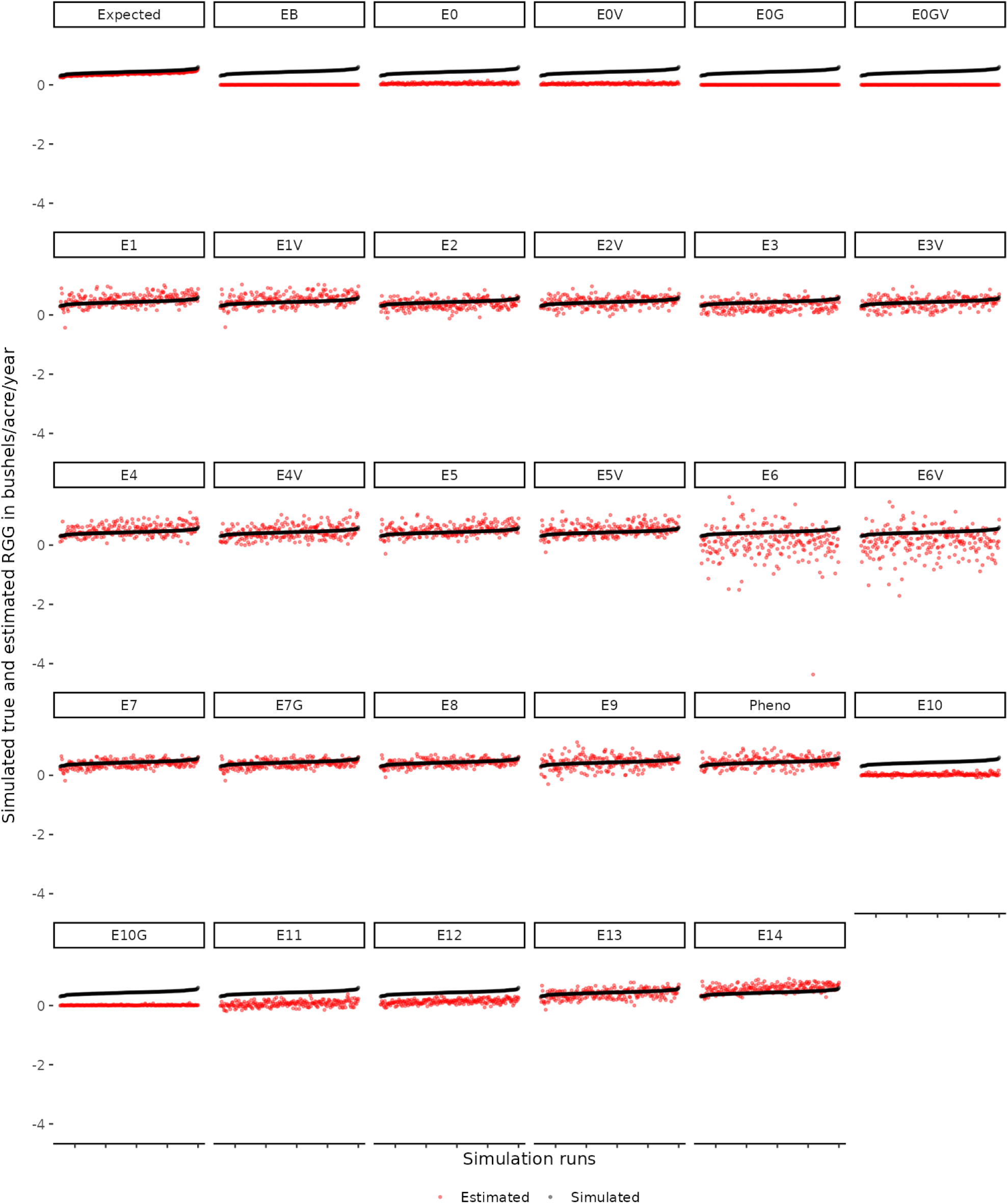
True (black) and estimated (red) RGG for evaluated models in scenario A2 (Tables 1 and 2). Check Section 4.2.4.3 for details on the “Expected” facet.

**Figure A11:**
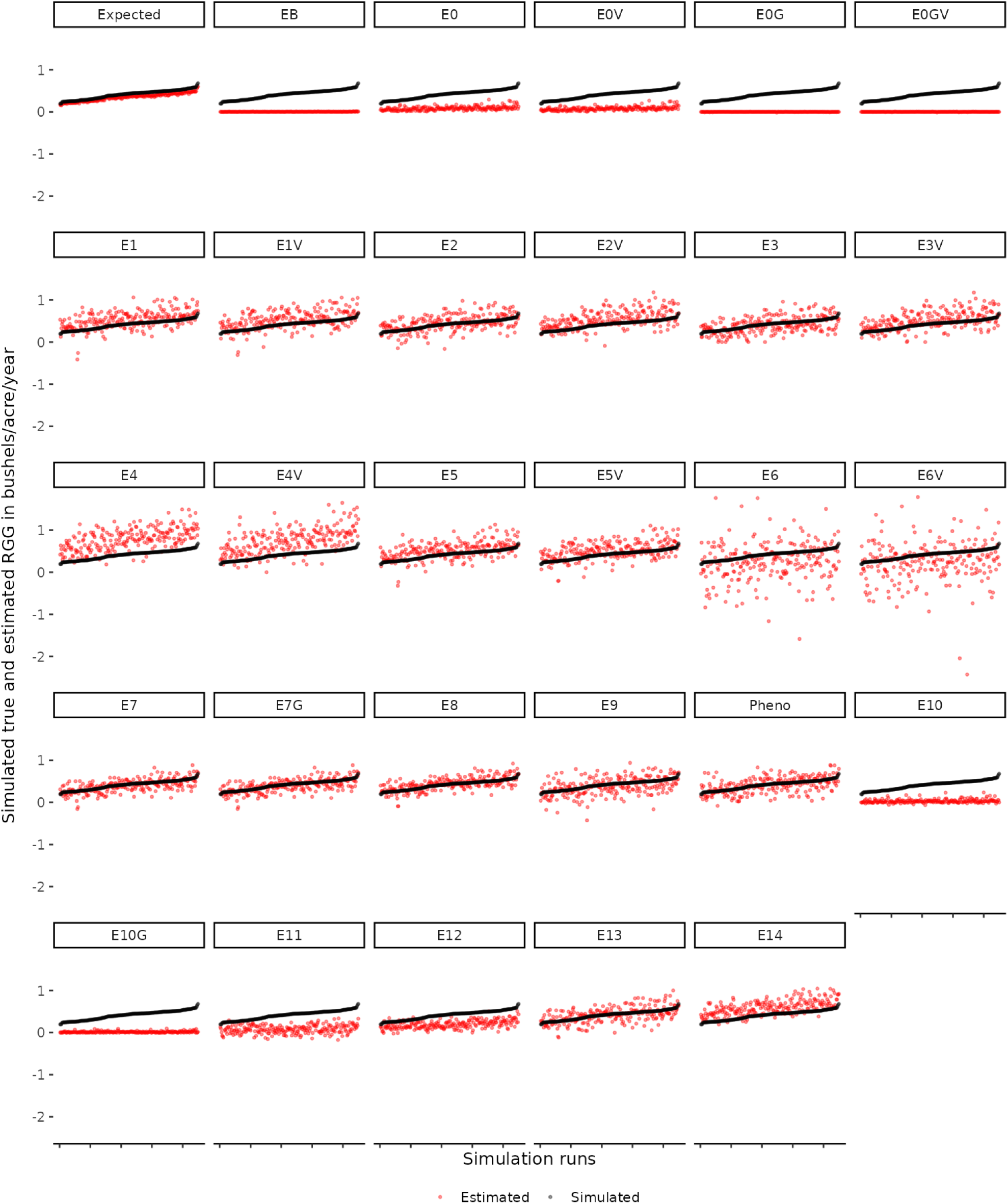
True (black) and estimated (red) RGG for evaluated models in scenario B1 (Tables 1 and 2). Check Section 4.2.4.3 for details on the “Expected” facet.

**Figure A12:**
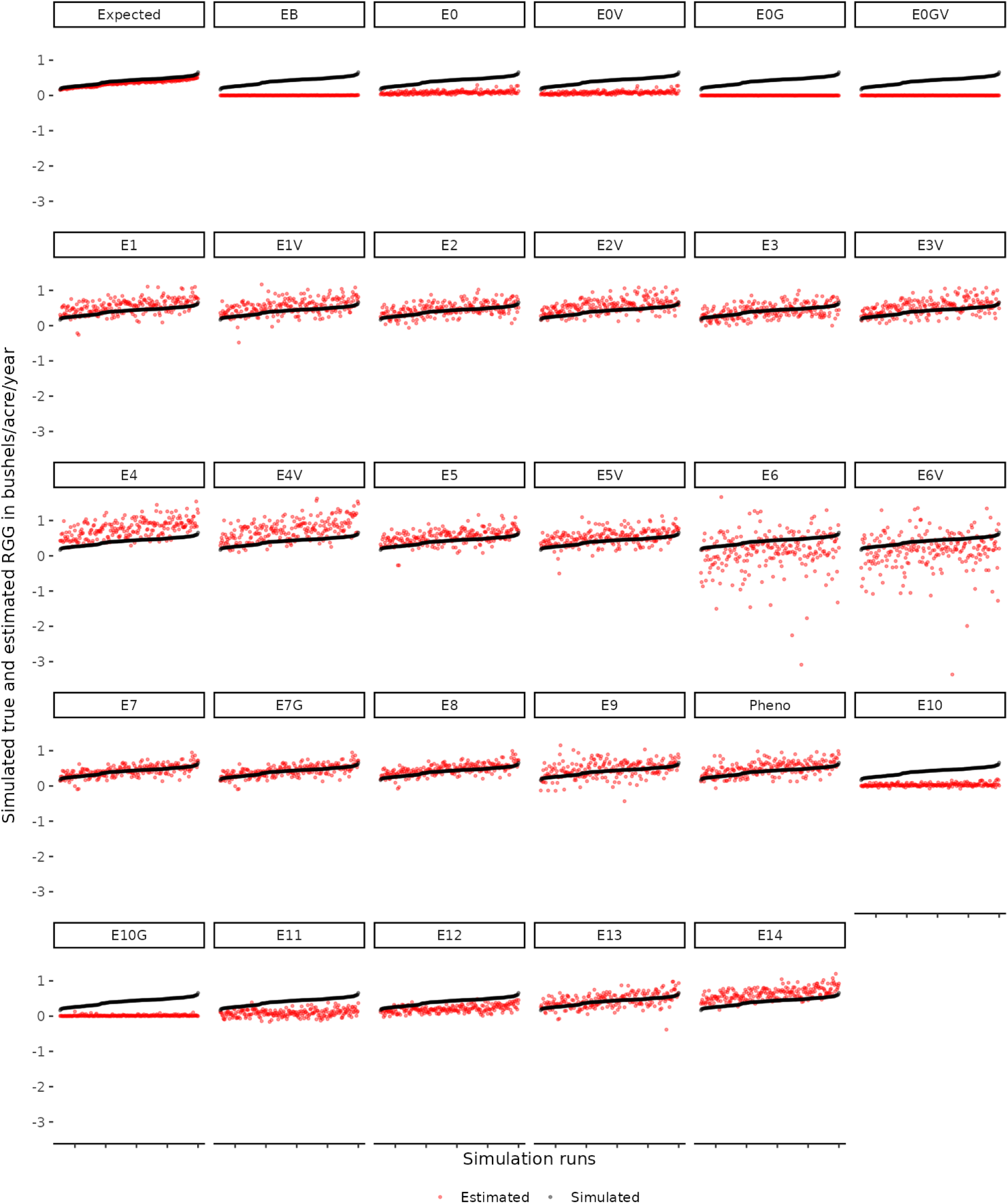
True (black) and estimated (red) RGG for evaluated models in scenario B2 (Tables 1 and 2). Check Section 4.2.4.3 for details on the “Expected” facet.

**Figure A13:**
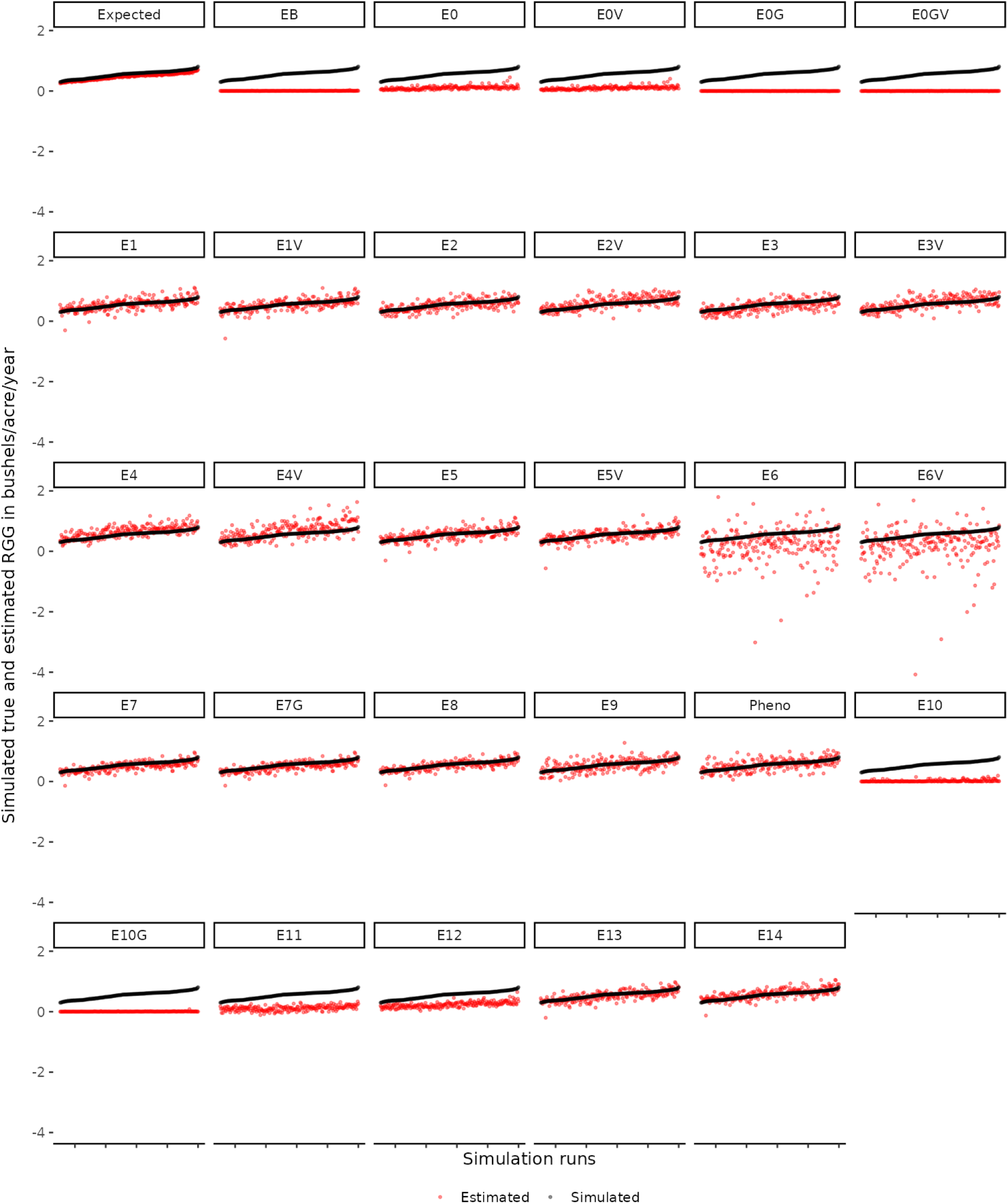
True (black) and estimated (red) RGG for evaluated models in scenario B2-M (Tables 1 and 2). Check Section 4.2.4.3 for details on the “Expected” facet.

**Figure A14:**
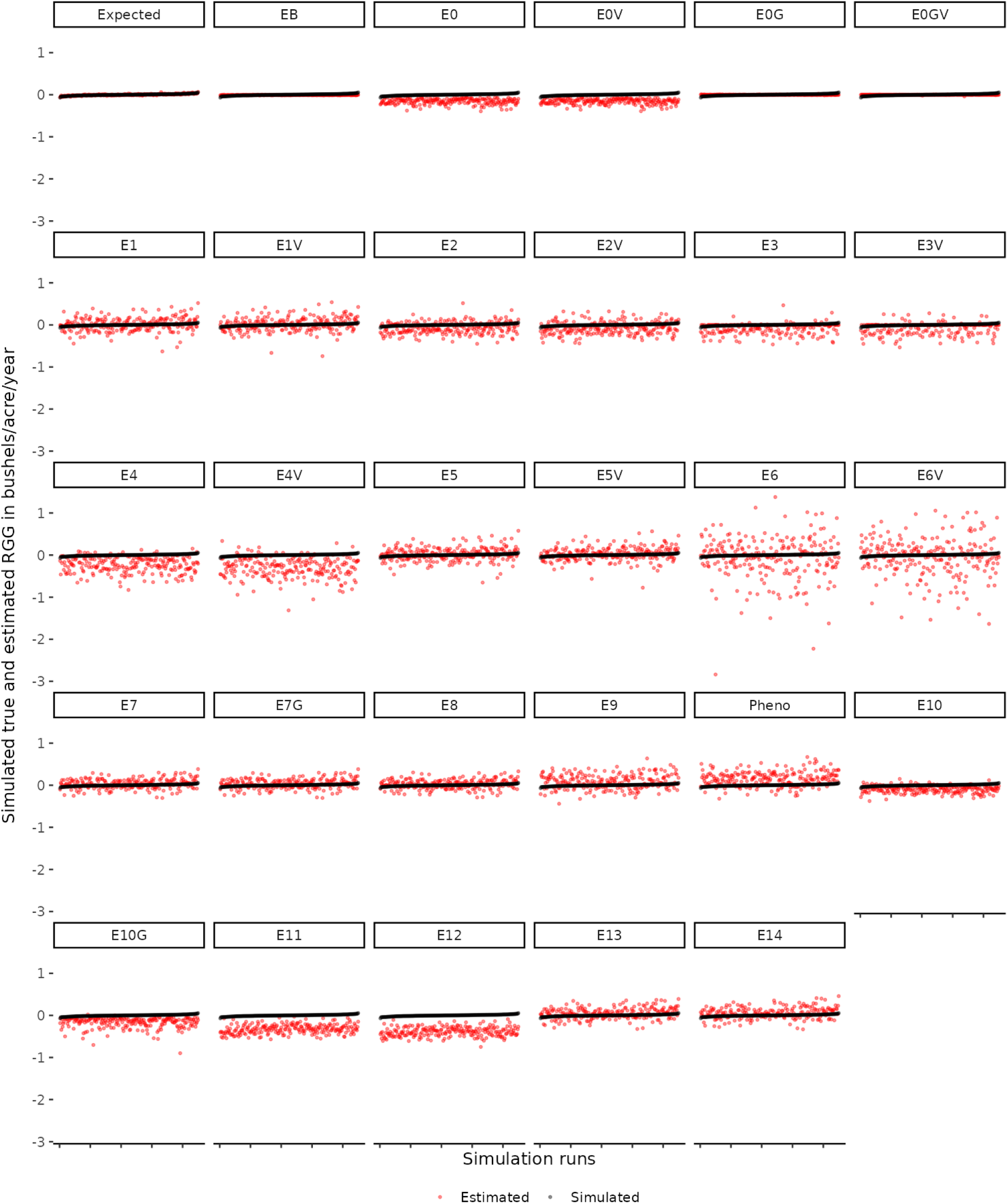
True (black) and estimated (red) RGG for evaluated models in scenario B2-R (Tables 1 and 2). Check Section 4.2.4.3 for details on the “Expected” facet.

**Figure A15:**
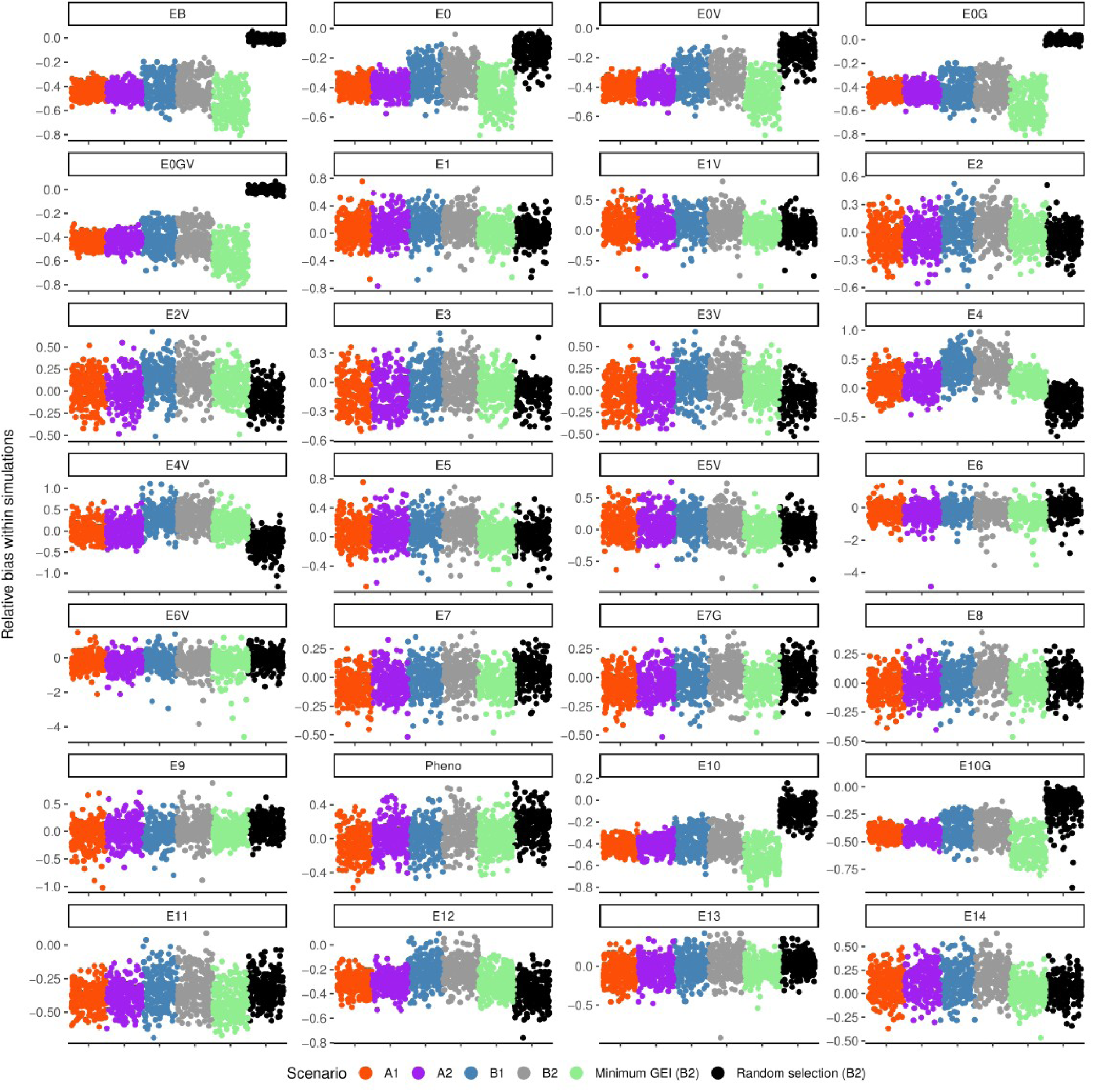
Estimated bias 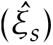 for each simulation run (*s* = 1*, . . .,* 225).

**Figure A16:**
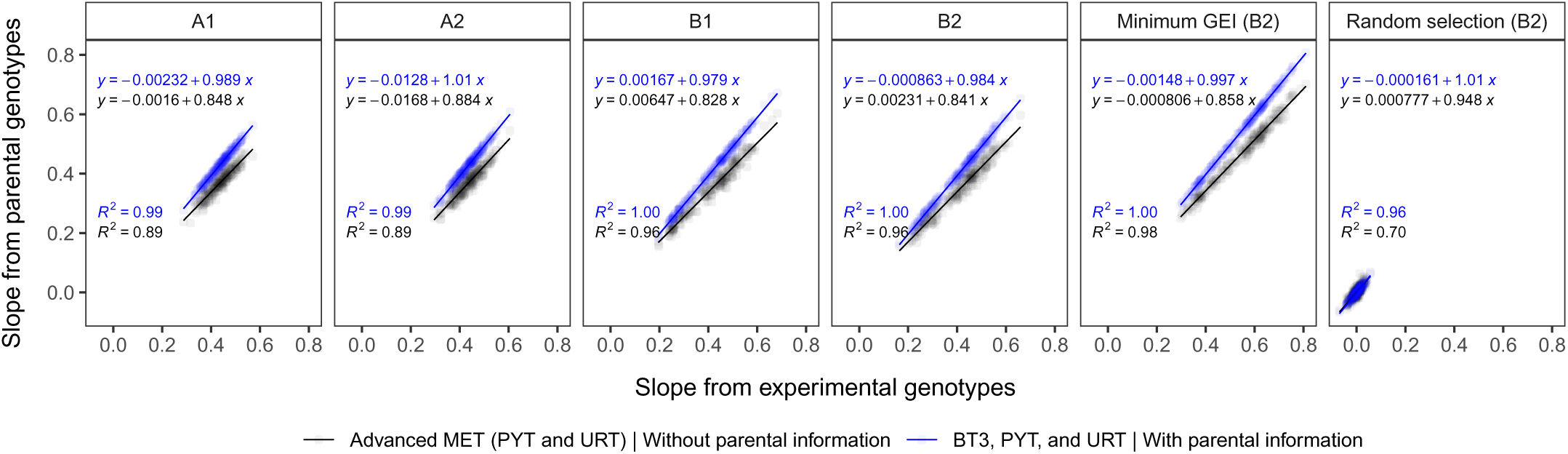
Linear regression (*y*) between true simulated genetic values of parental and experimental genotypes. The *R*^2^ represents the coefficient of determination from the fitted (blue/black) regression lines. The slope on the y-axis is *β_T_*_(*r*)_ from Equation 3, and on the x-axis is *β_R_*_(*r*)_ from Equation 4.

**Figure A17:**
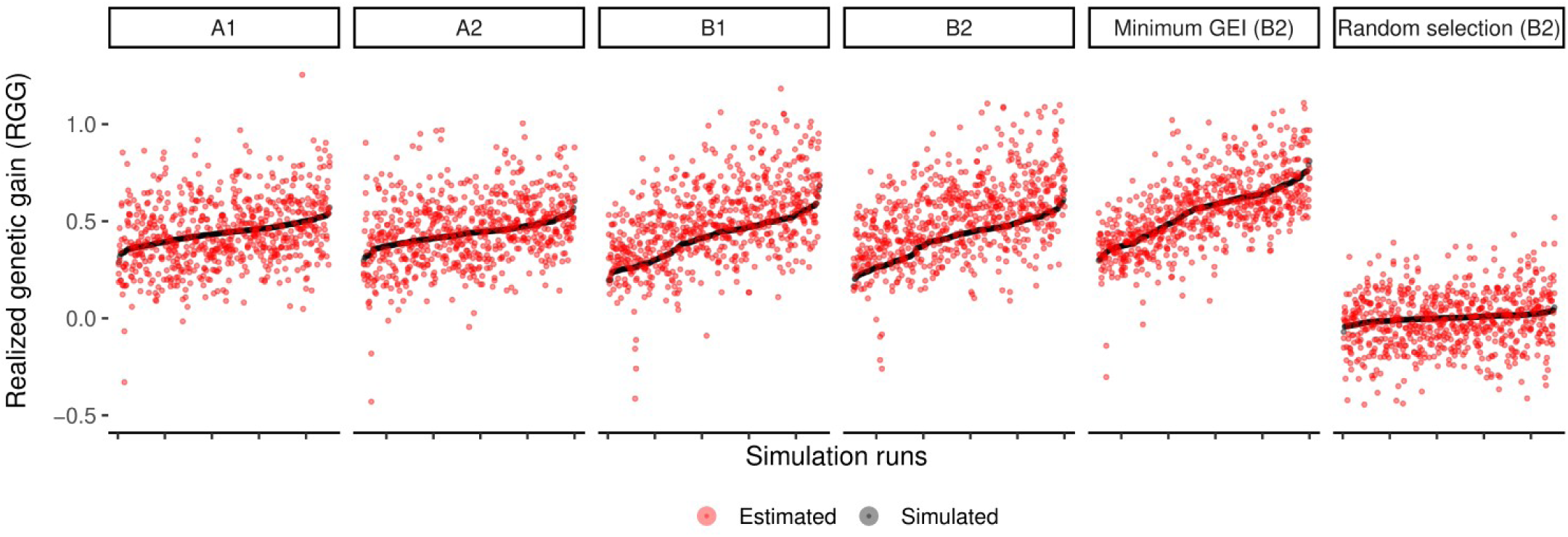
Scatter plot of simulated (black) and estimated (red) RGG values from Models E1, E2V, and E7 (Table 6).

**Table A1:** Number of positive (P) and negative (N) estimates of RGG.

| Scenarios | A1 |  | A2 |  | B1 |  | B2 |  | B2-M |  | B2-R |  |
| --- | --- | --- | --- | --- | --- | --- | --- | --- | --- | --- | --- | --- |
| Model | P | N | P | N | P | N | P | N | P | N | P | N |
| Simulated | 225 | 0 | 225 | 0 | 225 | 0 | 225 | 0 | 225 | 0 | 123 | 102 |
| EB | 135 | 90 | 140 | 85 | 181 | 44 | 184 | 41 | 219 | 6 | 7 | 218 |
| E0 | 223 | 2 | 224 | 1 | 225 | 0 | 225 | 0 | 225 | 0 | 0 | 225 |
| E0V | 224 | 1 | 224 | 1 | 225 | 0 | 225 | 0 | 225 | 0 | 0 | 225 |
| E0G | 64 | 161 | 69 | 156 | 62 | 163 | 80 | 145 | 67 | 158 | 93 | 132 |
| E0GV | 80 | 145 | 78 | 147 | 97 | 128 | 96 | 129 | 57 | 168 | 94 | 131 |
| E1 | 224 | 1 | 224 | 1 | 223 | 2 | 223 | 2 | 223 | 2 | 119 | 106 |
| E1V | 224 | 1 | 224 | 1 | 222 | 3 | 222 | 3 | 224 | 1 | 123 | 102 |
| E2 | 224 | 1 | 221 | 4 | 220 | 5 | 224 | 1 | 225 | 0 | 71 | 154 |
| E2V | 224 | 1 | 223 | 2 | 224 | 1 | 225 | 0 | 225 | 0 | 66 | 159 |
| E3 | 224 | 1 | 224 | 1 | 223 | 2 | 225 | 0 | 225 | 0 | 54 | 171 |
| E3V | 224 | 1 | 225 | 0 | 225 | 0 | 225 | 0 | 225 | 0 | 56 | 169 |
| E4 | 225 | 0 | 225 | 0 | 225 | 0 | 225 | 0 | 225 | 0 | 16 | 209 |
| E4V | 225 | 0 | 225 | 0 | 225 | 0 | 225 | 0 | 225 | 0 | 19 | 206 |
| E5 | 224 | 1 | 224 | 1 | 223 | 2 | 223 | 2 | 223 | 2 | 123 | 102 |
| E5V | 224 | 1 | 224 | 1 | 222 | 3 | 222 | 3 | 224 | 1 | 124 | 101 |
| E6 | 163 | 62 | 146 | 79 | 171 | 54 | 153 | 72 | 163 | 62 | 109 | 116 |
| E6V | 169 | 56 | 149 | 76 | 176 | 49 | 159 | 66 | 170 | 55 | 110 | 115 |
| E7 | 224 | 1 | 224 | 1 | 223 | 2 | 222 | 3 | 224 | 1 | 141 | 84 |
| E7G | 224 | 1 | 224 | 1 | 223 | 2 | 222 | 3 | 224 | 1 | 140 | 85 |
| E8 | 224 | 1 | 224 | 1 | 223 | 2 | 223 | 2 | 224 | 1 | 137 | 88 |
| E9 | 211 | 14 | 218 | 7 | 219 | 6 | 218 | 7 | 225 | 0 | 154 | 71 |
| Pheno | 223 | 2 | 225 | 0 | 224 | 1 | 224 | 1 | 225 | 0 | 189 | 36 |
| E10 | 174 | 51 | 182 | 43 | 188 | 37 | 184 | 41 | 216 | 9 | 29 | 196 |
| E10G | 203 | 22 | 208 | 17 | 221 | 4 | 221 | 4 | 220 | 5 | 2 | 223 |
| E11 | 158 | 67 | 157 | 68 | 182 | 43 | 184 | 41 | 202 | 23 | 0 | 225 |
| E12 | 207 | 18 | 212 | 13 | 221 | 4 | 223 | 2 | 225 | 0 | 0 | 225 |
| E13 | 224 | 1 | 224 | 1 | 222 | 3 | 222 | 3 | 224 | 1 | 150 | 75 |
| E14 | 225 | 0 | 225 | 0 | 225 | 0 | 225 | 0 | 224 | 1 | 151 | 74 |

**Figure A18:**
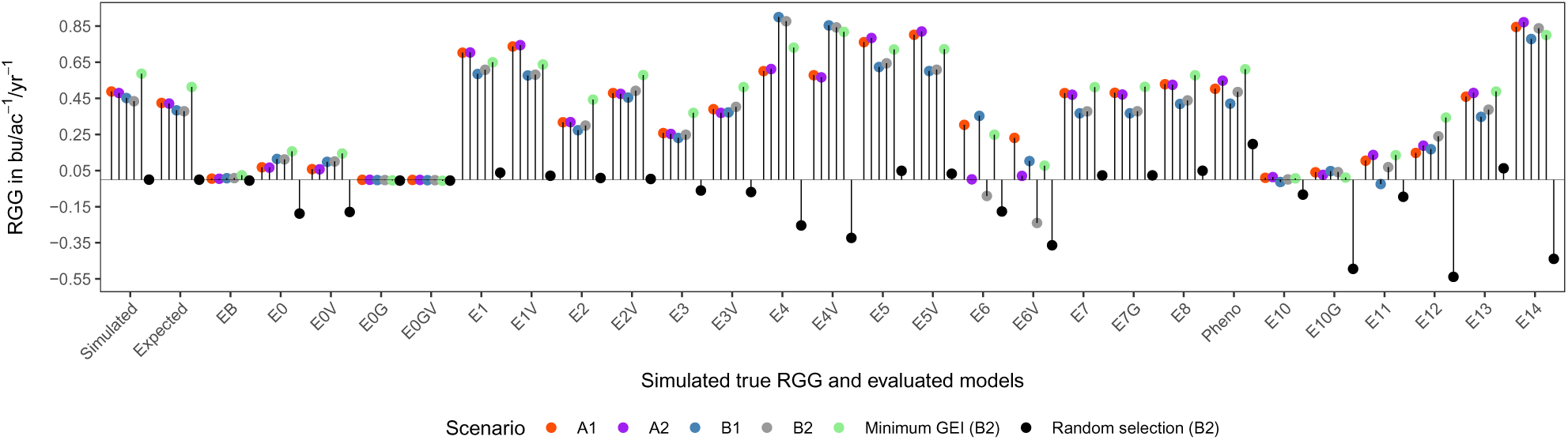
Average true simulated 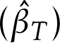, expected 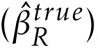, and estimated RGG 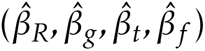, considering 10 years of MET. Model E9 (Table 6) was not included.

**Figure A19:**
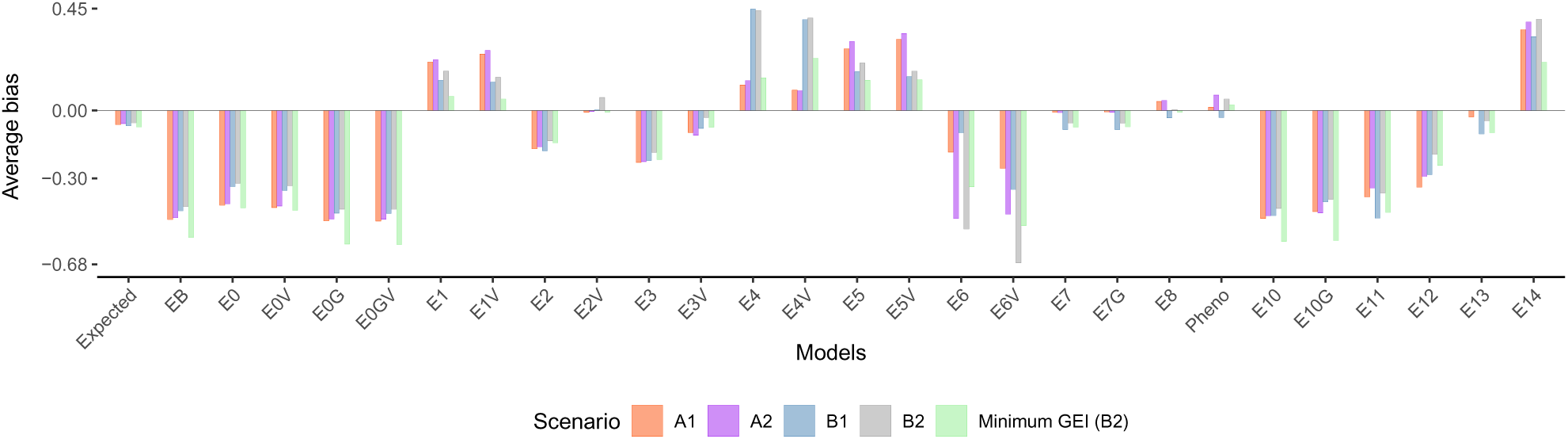
Average bias in bu/ac^−1^/yr^−1^ considering 10 years of MET. Model E9 (Table 6) was not included.

**Figure A20:**
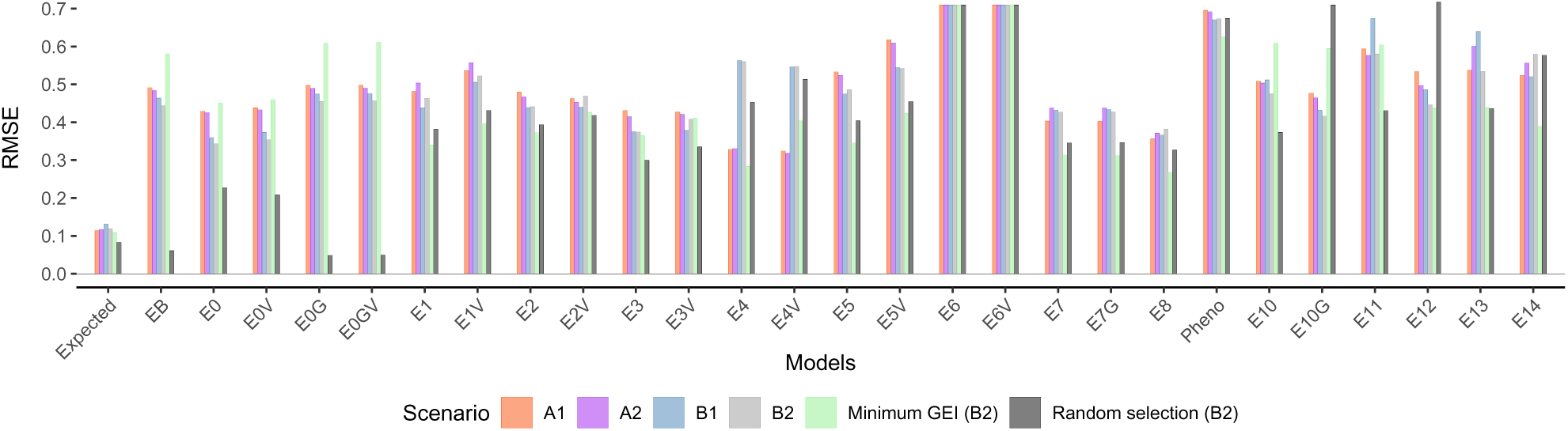
Root mean squared error (RMSE) considering 10 years of MET. Values were bounded at 0.72 units to enhance visualization. Model E9 (Table 6) was not included.

**Table A2:** Goodness-of-fit for models considered to estimate the RGG from the soybean empirical data. Fixed effects are underlined, and the RGG was estimated with model terms highlighted in gray. The numbers in parentheses (1, 2) in the model’s description represent steps to fit the models. Models in step one (1) were adjusted only with check cultivars. Models E2, E2_1_, *. . .*, E2_6_, were evaluated according to the Akaike information criterion (Akaike 1974, AIC), and the RGG reported with the selected model. For further details on model terms and covariance modeling, check Table 6.

| Models |  |  |  |  |  |
| --- | --- | --- | --- | --- | --- |
| Description | ID | VCOV |  |  | AIC |
| | | G | $\Sigma_{gl}$ | $\Sigma_{gy}$ | |
| $\hat{y}_{ijk} = \underline{G_i} + L_j + \underline{Y_k} + \underline{\delta z} + GL_{ij} + GY_{ik} + LY_{jk} + GLY_{ijk}$ | E1 | **I** | **I $\otimes$ I** | **I $\otimes$ I** | 159411.7 |
| $\hat{y}_{ijk} = \underline{\beta_g r_i} + L_j + \underline{\beta_t t_k} + \underline{\delta z} + GL_{ij} + GY_{ik} + LY_{jk} + GLY_{ijk}$ | E2 | - | **I $\otimes$ I** | **I $\otimes$ I** | 175764.9 |
|  | E2V<sub>1</sub> | - | **D $\otimes$ I** | **D $\otimes$ I** | 174366.1 |
|  | E2V<sub>2</sub> | - | **FA<sub>1</sub> $\otimes$ I** | **FA<sub>1</sub> $\otimes$ I** | 172889.9 |
|  | E2V<sub>3</sub> | - | **FA<sub>2</sub> $\otimes$ I** | **FA<sub>2</sub> $\otimes$ I** | 172582.2 |
|  | E2V<sub>4</sub> | - | **FA<sub>3</sub> $\otimes$ I** | **FA<sub>2</sub> $\otimes$ I** | 172380.3 |
|  | E2V<sub>5</sub> |  | **FA<sub>3</sub> $\otimes$ I** | **FA<sub>3</sub> $\otimes$ I** | 172399.0 |
|  | E2V<sub>6</sub> |  | **FA<sub>4</sub> $\otimes$ I** | **FA<sub>2</sub> $\otimes$ I** | 172326.3 |
| $\hat{y}_{ih} = \underline{G_i} + E_h + EG_{ih}$ (1)<br>$\hat{y}_{ih} = \underline{G_i} + \underline{\lambda_E \hat{E}_h} + G\hat{E}_h$ (2) | E7 | **I** | - | - | 211842.9 |

**Figure A21:**
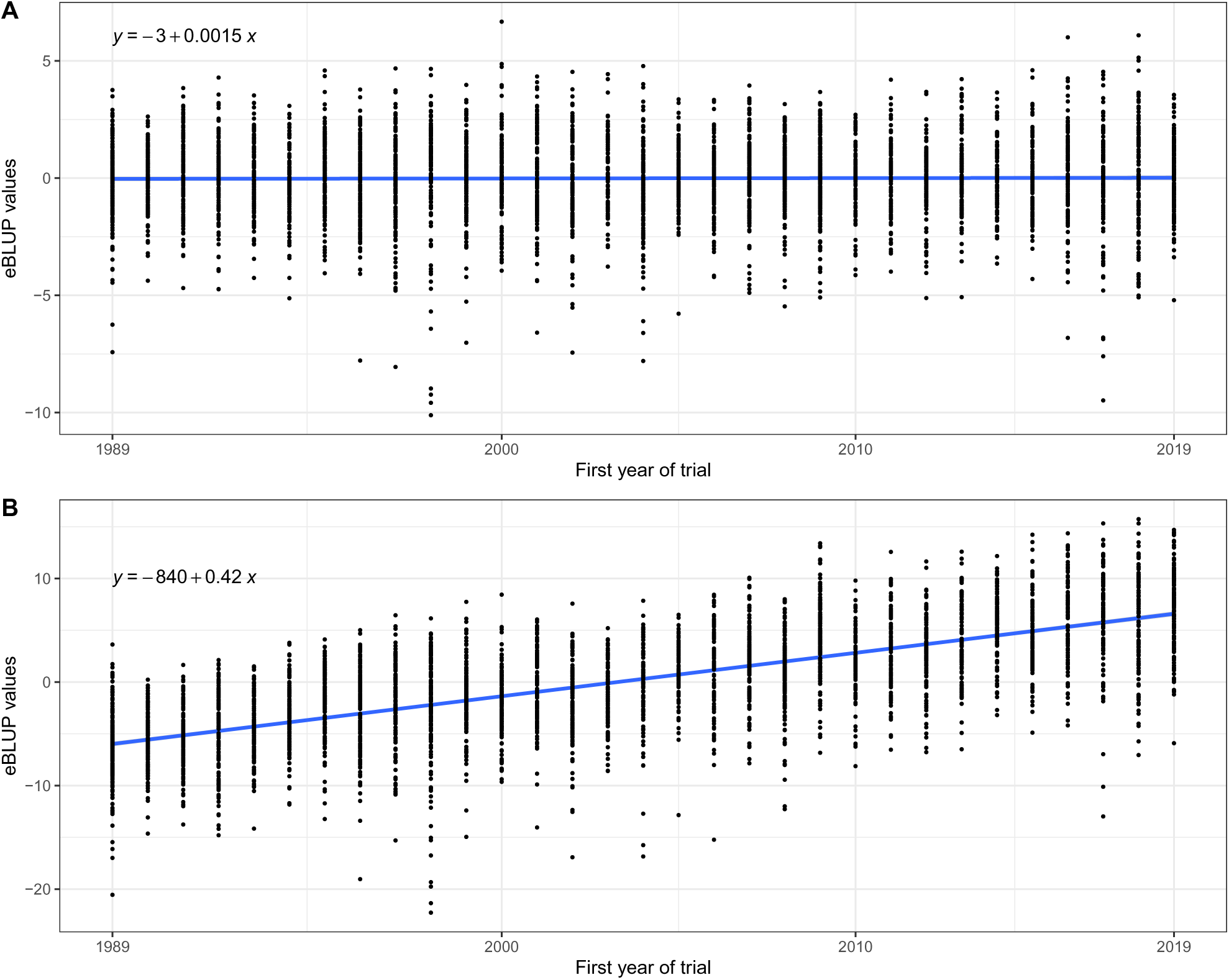
Predicted eBLUP values from the benchmark Model E0 (A) and from Model E7 (B) for the soybean empirical data. The blue line is from the fitted regression of the predicted values as a function of the first year of testing. For details on the models check Table 6.

**Figure A22:**
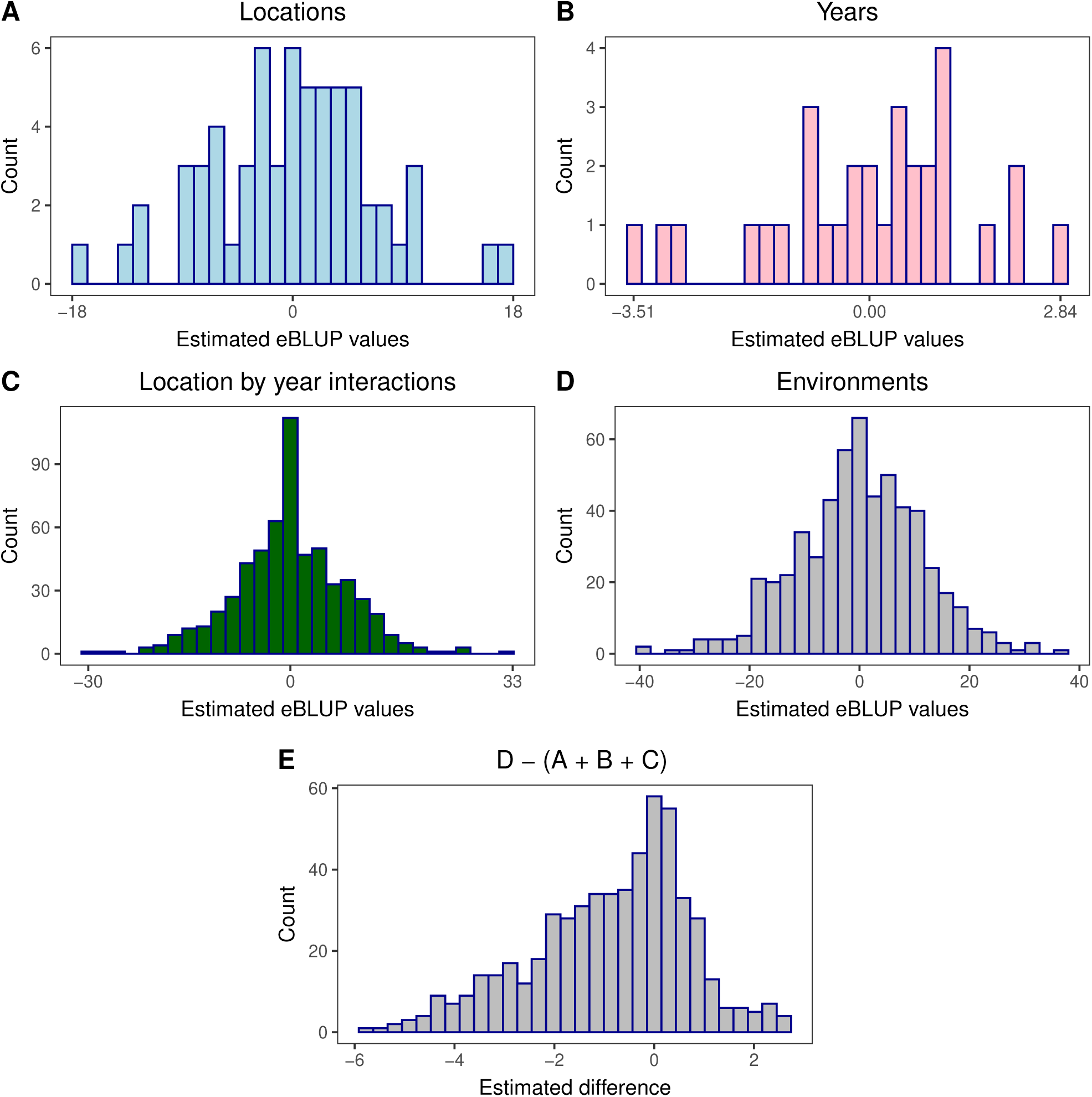
Estimated eBLUP values of locations (A), years (B), location by year interaction effects (C), and of environmental (location-year combination) effects (D) from check cultivars for the empirical data. These predicted values in D were used in Model E7 to calculate RGG for the empirical data.

**Figure A23:**
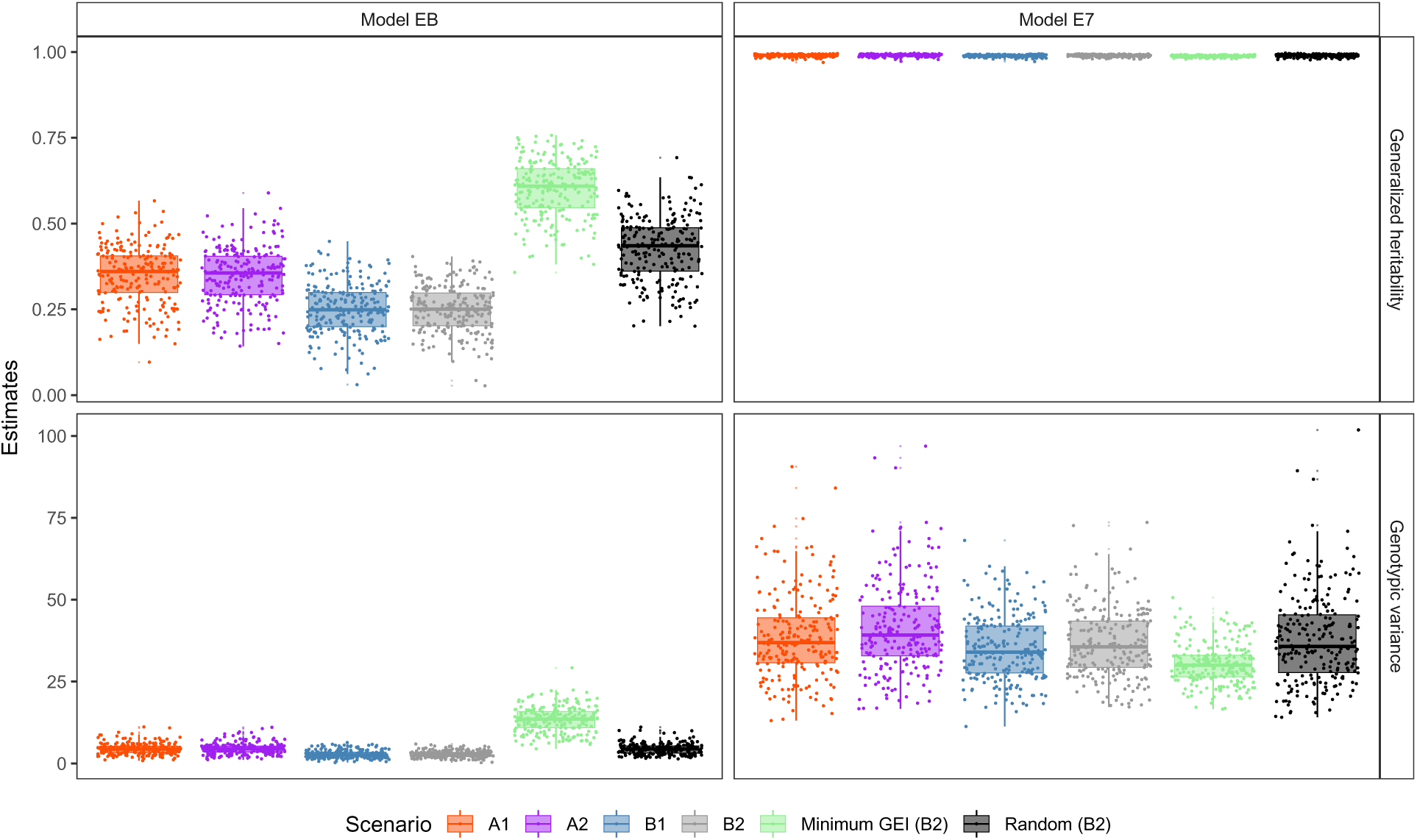
REML estimates of genotypic variance and generalized heritability (Cullis *et al*. 2006) for benchmark Model EB and E7 (Table 6).

## B Trait simulation in AlphaSimR

Additive (or total) QTL effects for the simulated traits in AlphaSimR (Gaynor *et al*. 2021) are simulated in two steps. The first one consists of sampling the effects either from a Standard Normal or Gamma distribution ((the user must define_r_its shape parameter). The second step applies a scaling factor 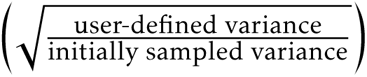to match the user-defined additive (or total) variance. Then, an intercept (user-defined) is added to the simulated (and now scaled) QTL effects. This step mimics the introduction of a trait mean reference to which trait the user is simulating. In the case of a purely additive trait, the variance of the simulated breeding values in the founder population will match the user-defined additive variance. For further details, check the vignette “Traits in AlphaSimR” (https://cran.r-project.org/web/packages/AlphaSimR/vignettes/ traits.pdf, accessed on 09/07/2022).

## C Phenotypic models used in the simulator for parental selection, yearly selections, and for single-trial analyses

Parental genotypes were selected in BT3 after three years of trials. Data from BT1, BT2, and BT3 was combined, and the following linear mixed model was considered:

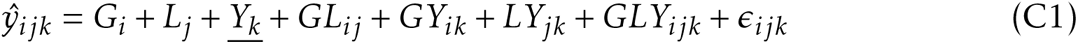

where 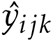 is the vector of eBLUE values of genotypic means for the *ijk^th^* genotype × location × year combination estimated from single-trial analyses, and *G_i_*, *L_j_*, and *Y_k_*, are the main effects of the *i^th^* genotype, *j^th^* location, and *k^th^* years, respectively, followed by all two and three-way interaction effects. All model terms but *L_j_* (underlined) were considered as random effects. The error variance 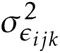 of *ɛ_ijk_* is assumed to be known from the analyses of the individuals trials (Model C3), where the residual variance matrix is a diagonal matrix with elements equal to 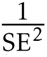, where SE is the estimated standard error (Smith *et al*. 2001a; Frensham *et al*. 1997). Yearly selections and the analyses of single-trials were performed with the following models, respectively:

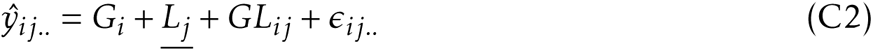

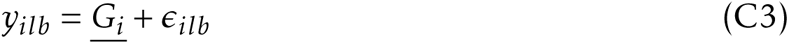

where *y_ilb_* is the observed yield in a plot level at any given trial. The eBLUE values of genotypic 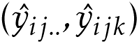 means were estimated with weighted least squares, so that 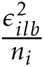, where *n_i_* is the number of plots of the *i^th^* genotype in the *b^th^* trial. Thus, genotypes have heterogeneous SE within trials due to the insertion of missing data, i.e., a random proportion from zero to 12% of the phenotypic data was considered missing data within trials. A basic explanation of stage-wise modeling is given in Appendix F.

## D Simulation of correlated additive QTL effects for GL, GY, and GLY

In terms of variance components, the genotype by environment interactions (GEI) is composed of genotype × location 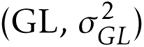, genotype × year 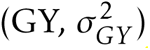, and genotype × location × year 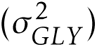 interaction variances (Krause et al. 2022). Both 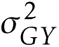 and 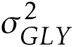 are non-static (unrepeatable) sources of variation, while the static portion of the 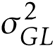 is repeatable across years (Yan 2016). Krause *et al*. (2022) investigated the GEI patterns in a historical soybean dataset composed of 39,006 phenotypic records from 1989-2019. This data comprises trials conducted by public plant breeders in the Northern Region of the U.S., and it embraces 4,257 experimental genotypes, 63 locations, and 31 years. The modeling scenarios B1 and B2 (Section 4.1.6.1) were designed to capture the structure of this historical data as best as possible.

The following algorithm is a heuristic approach built to simulate correlated additive QTL effects in multi-environment trials (MET) according to GL and GY effects. For example, if 1,000 QTLs were simulated and the goal is phenotyping in *L* = 10 locations across *Y* = 2 years, the algorithm will initially sample 10 unique variance components for GL and two unique variance components for GY. The variances for individual locations and years are sampled from a mixture of three and six Log-Logistic distributions, respectively (Tables 1 and E1). For more details on how these empirical probability distributions of variance components were obtained, please refer to Krause *et al*. (2022). The initial diagonal variance-covariance (VCOV) matrices for GL (Σ*_GL_*) and GY (Σ*_GY_*) are as follows:

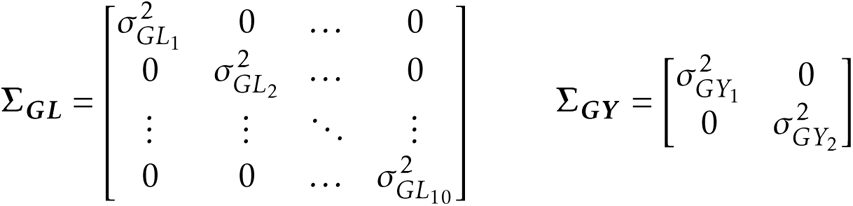

These variance components (i.e., diagonal matrices) are then set relative to the predefined additive genetic variance 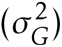 of the simulated trait. For example, if the predefined 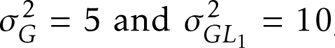, then the variance of GL effects represents two times the magnitude of the additive genetic variance, or 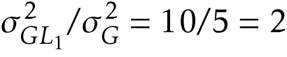. As a reminder, scenarios A define a unique 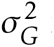 from a point estimate of variance, and scenario B sample from empirical distributions a unique 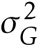 in each simulation run 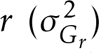. Next, the relative variances are multiplied by the variance of the simulated additive effects (*γ*), as follows:

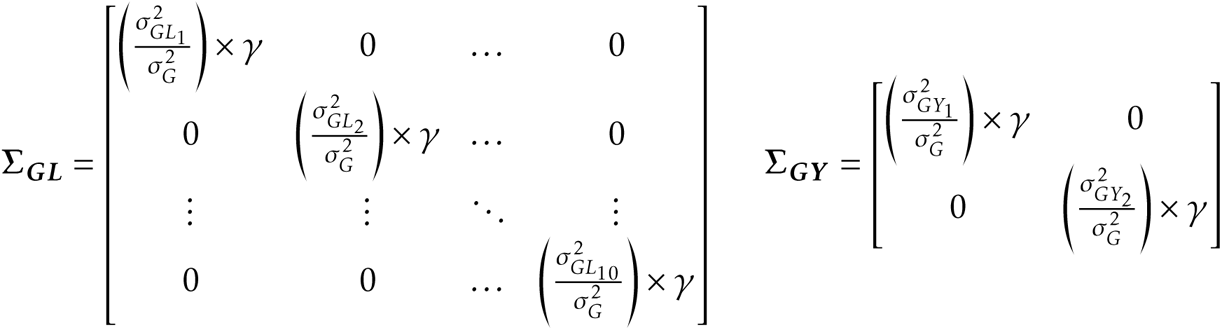

Which are equivalent to:

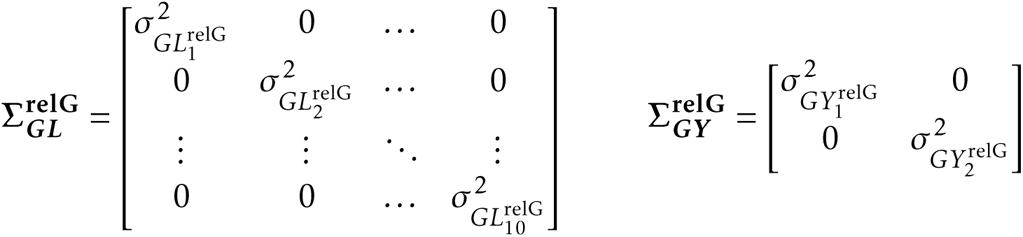

Where the “relG”stands for relative to the genetic variance. The VCOV matrices 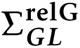 and 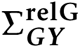 are now modified to include covariances between locations and years, receptively. Genetic correlation between 63 locations and 31 years was obtained by Krause *et al*. (2022). Briefly, the authors fitted 20 linear mixed models to the soybean historical dataset (see Table 1 in their publication) and selected the best-fit one according to some evaluation criteria. Correlations were then calculated from the best-fit model which included factor-analytic (FA) covariance structures for both GL (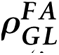, Figure D2) and GY (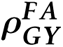 Figure D1) model terms. With the pre-sampled variances (i.e., the diagonal values from 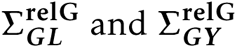) and given the correlations (*ρ_l,l_*′, *ρ_y,y_*′) are known, the covariances among any pair of years and/or locations were computed as 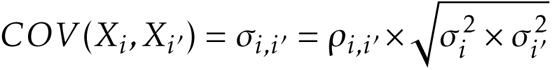 Thus,

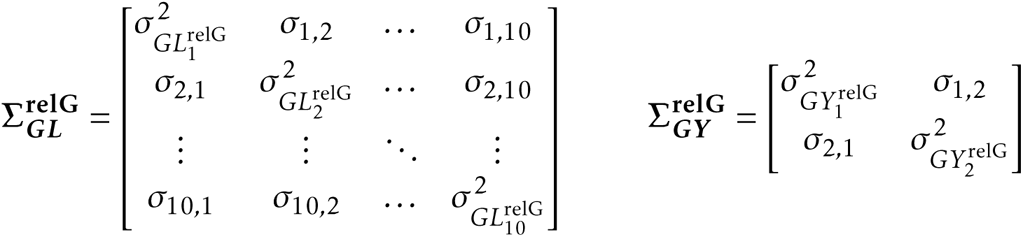

Correlated additive QTL effects can now be drawn from a Multivariate Normal Distribution according to the computed covariance matrices above. So, the distributions of the QTL effects for GL and GY are as follows:

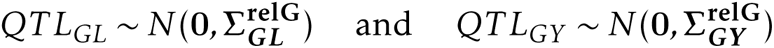

And the final matrices of QTL effects are of the following form:

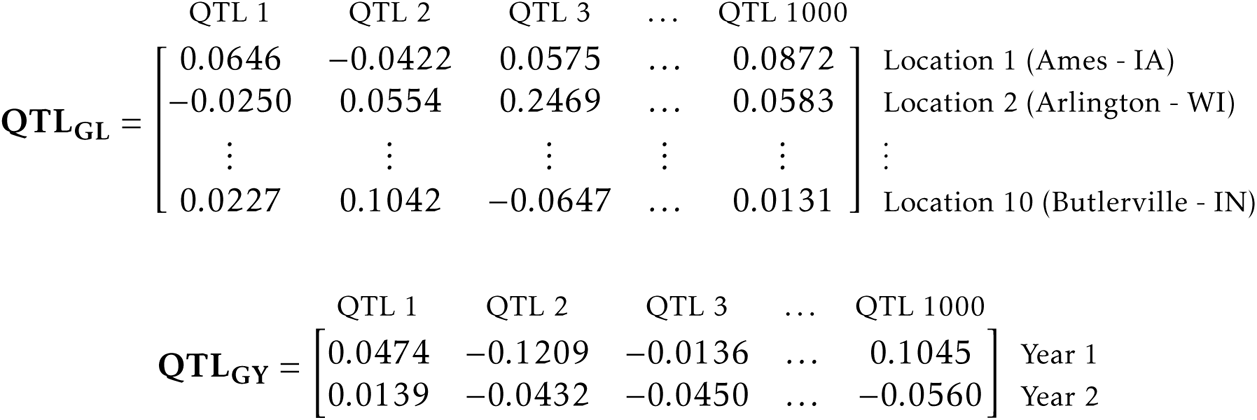

The original correlation matrices for GL and GY obtained by Krause *et al*. (2022) from FA models contained 63 locations and 31 years, respectively. For this work, a unique simulation run never explored more than 63 locations across the 42 years of the simulated breeding program. Hence, there was enough empirical information in 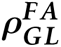 (Figure D2) to simulate correlated GL effects. On the other hand, for the GY effects, the correlation matrix is of dimension 31 × 31 (Figure D1). To overcome this issue, our algorithm symmetrically augments 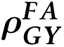 according to the desired number of years. Briefly, the algorithm explores the empirical correlations to simulate the first 31 years of MET. New correlations are then simulated by adding noise, in a highly controlled manner (Hardin *et al*. 2013, function noisecor), to the empirical values. If the resulting covariance matrix is not positive-definite (PD), it computes the nearest PD matrix while keeping the pre-sampled variances (diagonal values) constant (Bates and Maechler 2021, function nearPD). Examples of the sampled QTL effects are shown below in Figures D3 and D4.

Correlated additive QTL effects for scenarios B1 and B2 were computed as described up to this point. For scenarios A1 and B1, the simple effects/variance scenarios, independent QTL effects were drawn according to a compound symmetry structure as follows:

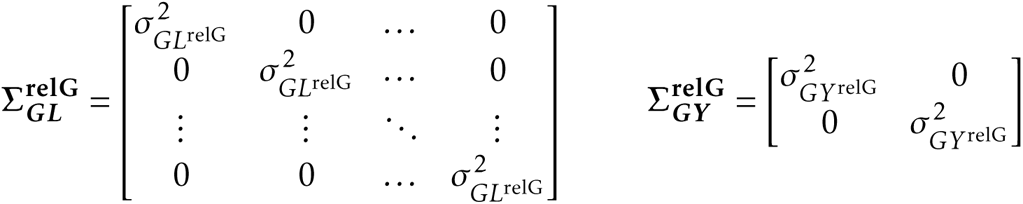

Where there is a unique variance for GL and GY, set relative to the genetic/additive variance. Now it is straightforward to demonstrate how GLY was simulated for the simple effects scenarios (A’s) based on the following covariance matrices:

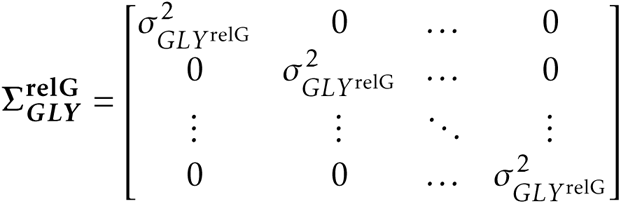

And finally for the complex effects scenarios (B’s):

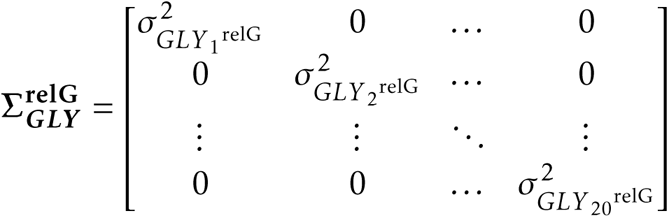

**Figure D1:**
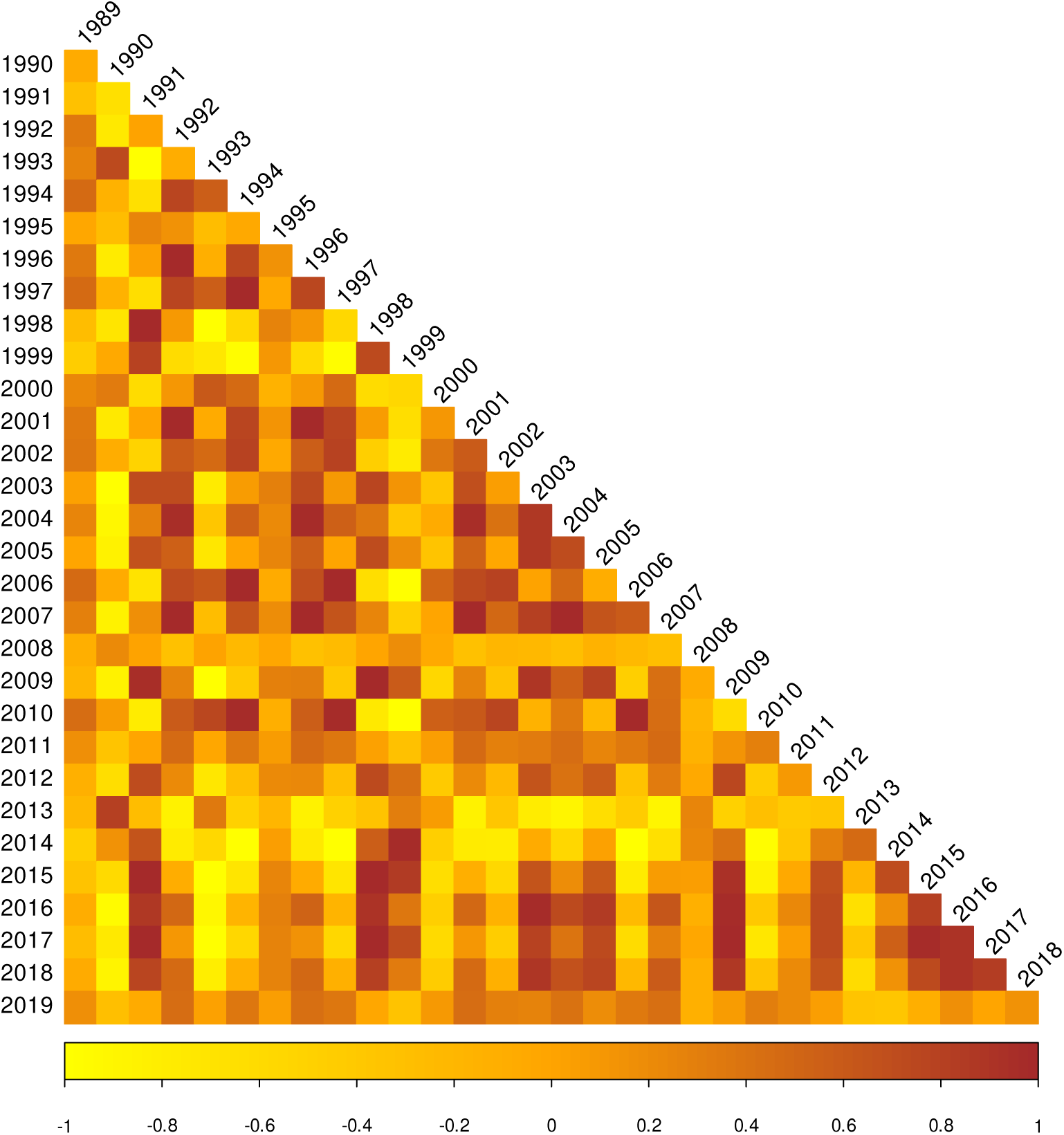
Phenotypic correlations between years 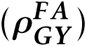 estimated from FA multiplicative model. For details, check Model M3-18 from Krause *et al*. (2022).

**Figure D2:**
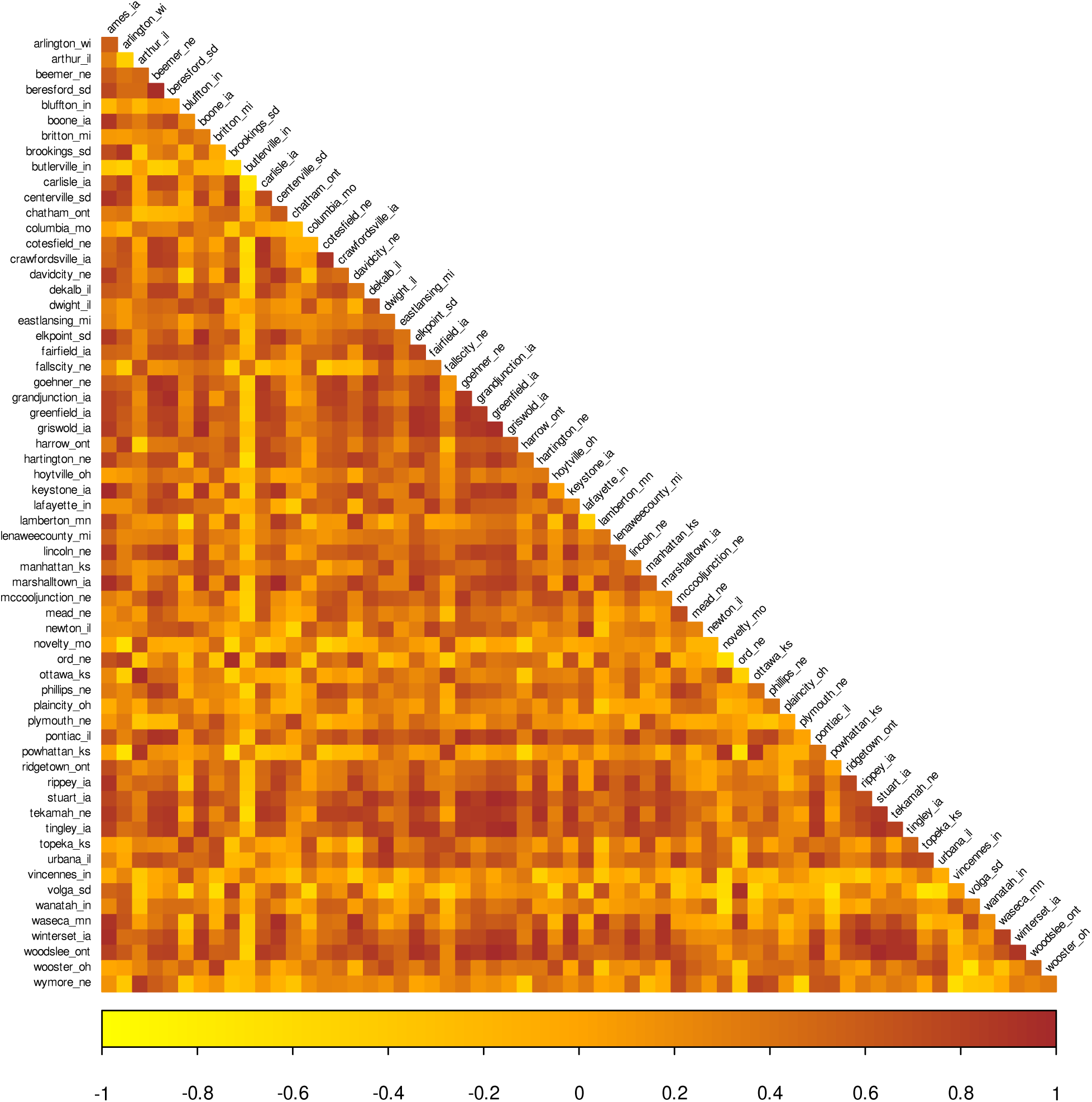
Phenotypic correlations between locations 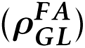 estimated from FA multiplicative model. For details, check Model M3-18 from Krause et al. (2022).

**Figure D3:**
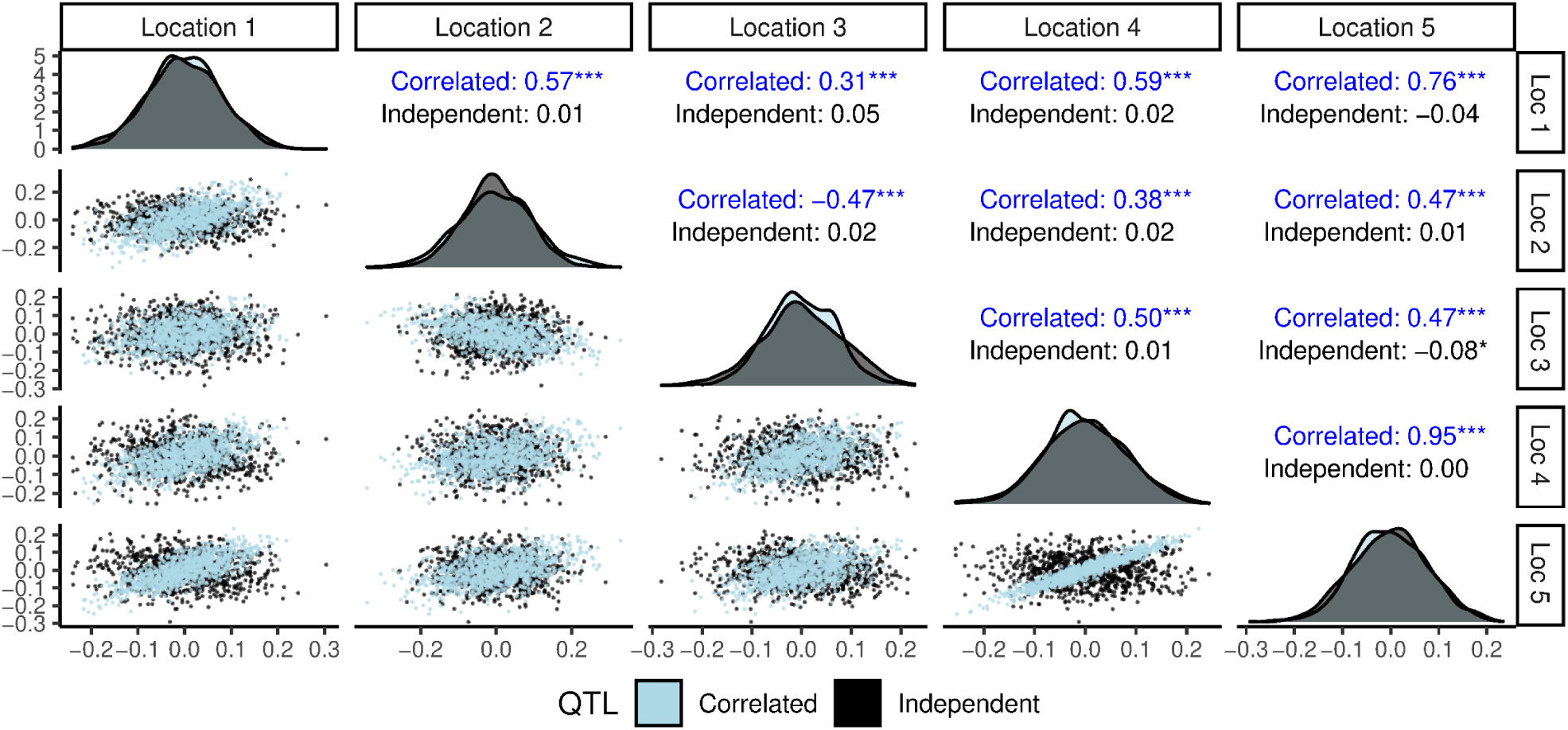
An example of sampling of QTL effects according to correlated or independent genotype × location interaction effects. Pearson correlation coefficients followed by their significant tests are plotted in the upper diagonal.

**Figure D4:**
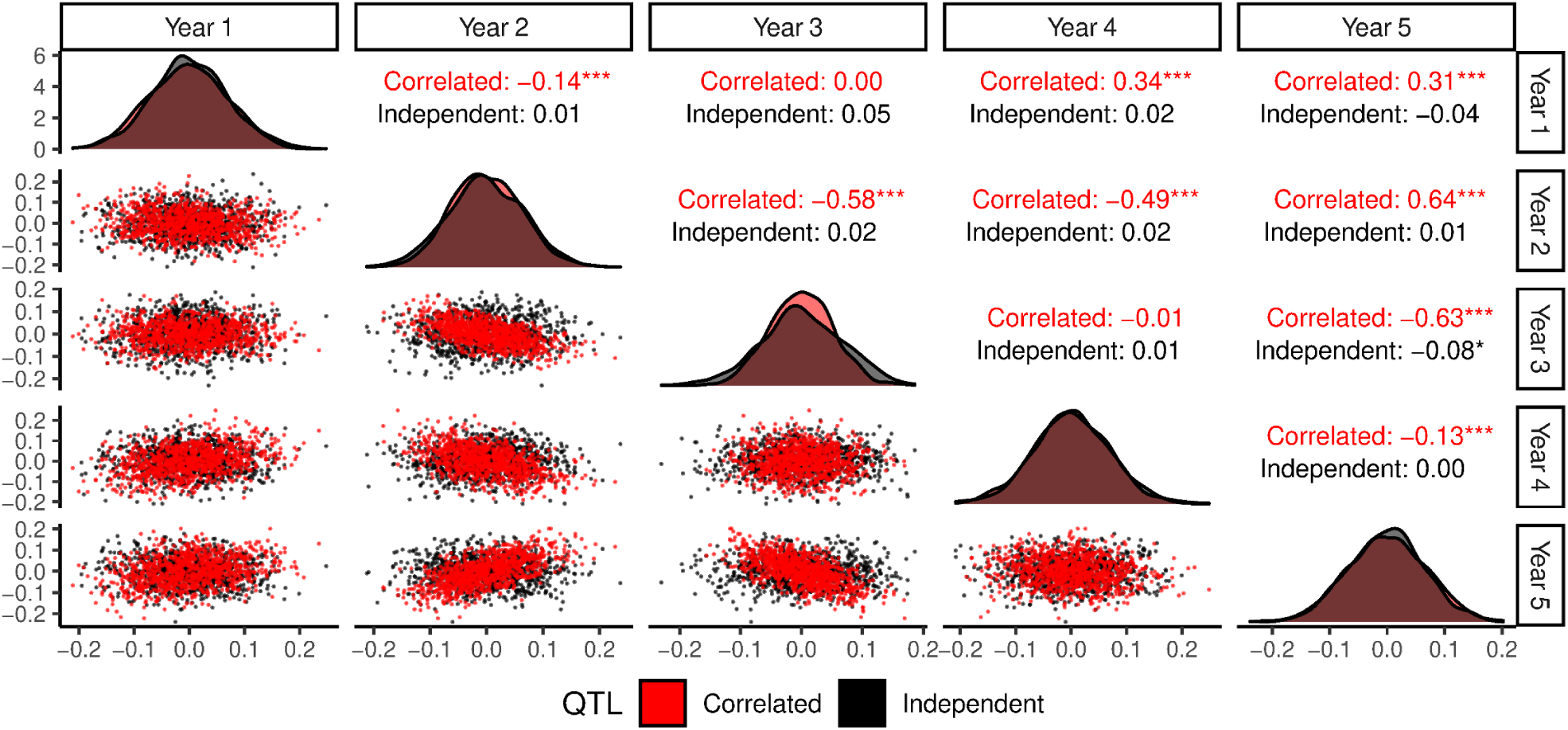
An example of sampling of QTL effects according to correlated or independent genotype × year interaction effects. Pearson correlation coefficients followed by their significant tests are plotted in the upper diagonal.

## E Empirical density functions of variance components

**Table E1:** Maximum Likelihood estimates of parameters for the best-fit univariate and multivariate probability distributions for empirical distributions obtained using jack-knife resampling. Estimates of residual variance 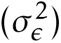 were obtained from trials conducted from 1989 to 2019. Original Table 3 from Krause *et al*. (2022).

| Variance<sup>a</sup> | Distribution | Number of distributions | Parameters<sup>b</sup> |  |  |
| --- | --- | --- | --- | --- | --- |
| | | | $\hat{w}_i$ | $\hat{\alpha}_i$ | $\hat{\beta}_i$ |
| $\sigma_G^2$ | Log-Logistic | 5 | 0.30 | 125.98 | 3.54 |
|  |  |  | 0.14 | 119.87 | 6.68 |
|  |  |  | 0.08 | 44.03 | 7.23 |
|  |  |  | 0.01 | 88.03 | 6.18 |
|  |  |  | 0.47 | 192.69 | 7.53 |
| $\sigma_{GL}^2$ | Log-Logistic | 3 | 0.63 | 122.27 | 4.23 |
|  |  |  | 0.17 | 32.00 | 3.12 |
|  |  |  | 0.20 | 92.56 | 7.62 |
| $\sigma_{GY}^2$ | Log-Logistic | 6 | 0.02 | 258.41 | 3.77 |
|  |  |  | 0.31 | 172.61 | 3.91 |
|  |  |  | 0.20 | 89.05 | 3.51 |
|  |  |  | 0.30 | 94.97 | 2.06 |
|  |  |  | 0.02 | 32.81 | 2.64 |
|  |  |  | 0.15 | 57.79 | 3.01 |
| $\sigma_{GLY}^2$ | Gamma | 5 | 0.05 | 4078.04 | 0.0024 |
|  |  |  | 0.31 | 7755.00 | 0.0011 |
|  |  |  | 0.20 | 5306.62 | 0.0025 |
|  |  |  | 0.28 | 31922.50 | 0.0003 |
|  |  |  | 0.16 | 7301.02 | 0.0020 |
| $\sigma_\epsilon^2$ | Log-Logistic | 1 | 1 | 2.56 | 17.03 |

<sup>a</sup> Genotypic ( $\sigma_G^2$ ), genotype × location ( $\sigma_{GL}^2$ ), genotype × year ( $\sigma_{GY}^2$ ), and genotype × location × year ( $\sigma_{GLY}^2$ ) variance components.
<sup>b</sup> Estimates of weight parameters ( $w_i$ ) sums to one, and both Gamma and Log-Logistic distributions include a shape ( $\alpha_i$ ) and scale ( $\beta_i$ ) parameter.

## F Stage-wise analysis

All models were obtained via stage-wise analysis with weights equal to 1*/SE*^2^, where *SE* is the standard error of genotypic means (Smith *et al*. 2001b; Frensham *et al*. 1997). Stage-wise models break down the full analysis (i.e., single-stage analysis) into multiple steps, by carrying to the next step the estimated means and some measure of precision (Piepho *et al*. 2012; Smith *et al*. 2001a; Frensham *et al*. 1997).

## G Model E9

Model E9 (Table 6) is based on the methodology proposed by Vencosvsky *et al*. (1986) and later modified/applied by Brisson *et al*. (2010), Oury *et al*. (2012), and Bornhofen *et al*. (2018). This model takes advantage of both experimental genotypes and checks replicated between consecutive pairs of years: (*i*) an initial model is fitted within years to predict eBLUP values of genotypic means (*g_ik_*); and (*ii*) the *g_ik_* values are then corrected for the year effect according to the “reference year (*R*)”. For example, in a MET dataset composed of 10 years, if *R* = year 5, the corrections for the year effects will be performed like 1 ← 2 ← 3 ← 4 ← 5 → 6 → 7 → 8 → 9 → 10, a forward-backward process along years. If *R* = 1, it is only a forward process (1 → · · · → 10), and if *R* = 10, a backward process (1 ← · · · ← 10). The algorithm is described with the following equations:

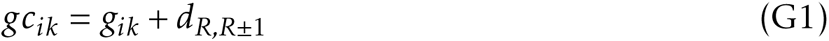

where *gc_ik_* is the corrected value for genotype *i* in year *k*, and *d_R,R_*_±1_ is the correction term obtained by Equation G2:

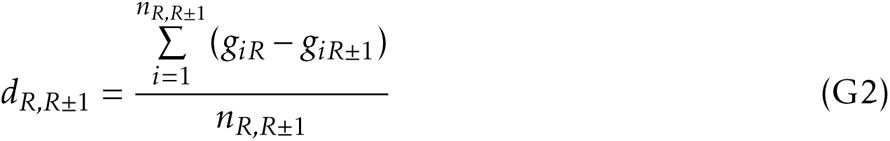

where *n_R,R_*_±1_ is the number of common genotypes in years *R* and *R* ± 1. The ± sign states the correction can go forward or backward across years. In the following year (*R* ± 2), the procedure is the same:

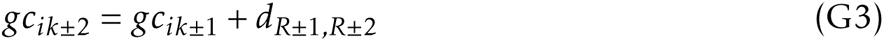

where *d_R_*_±1*,R*±2_ is:

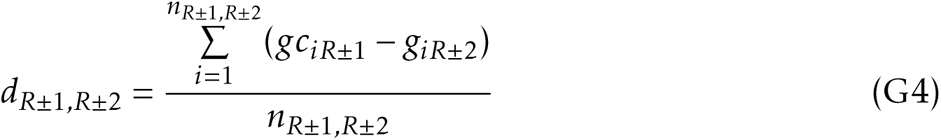

and so on for the following years. The estimated “year effects” 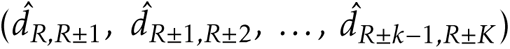 are calculated relatively to the reference year, *R*. In our simulations, the reference year was randomly chosen among the five years with the highest generalized measure of heritability (Cullis *et al*. 2006, *H*^2^), calculated as:

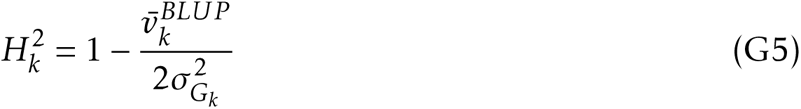

where 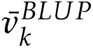 is the average pairwise genotypic prediction error variance and 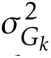 is the genotypic variance estimated from residual maximum likelihood (Patterson and Thompson 1971, REML). In step (*i*), before correction, the intercept (*µ*) was added to the eBLUP values. *µ* was parameterized to represent the average least-square value of the locations observed that year.

